# Efficient Detection and Classification of Epigenomic Changes Under Multiple Conditions

**DOI:** 10.1101/864124

**Authors:** Pedro L. Baldoni, Naim U. Rashid, Joseph G. Ibrahim

**Affiliations:** Department of Biostatistics, University of North Carolina at Chapel Hill, Chapel Hill, North Carolina, USA

**Keywords:** ChIP-seq, Differential Peak Call, Epigenomics, Hidden Markov Model, Mixture Model

## Abstract

Epigenomics, the study of the human genome and its interactions with proteins and other cellular elements, has become of significant interest in recent years. Such interactions have been shown to regulate essential cellular functions and are associated with multiple complex diseases. Therefore, understanding how these interactions may change across conditions is central in biomedical research. Chromatin immunoprecipitation followed by massively-parallel sequencing (ChIP-seq) is one of several techniques to detect local changes in epigenomic activity (peaks). However, existing methods for differential peak calling are not optimized for the diversity in ChIP-seq signal profiles, are limited to the analysis of two conditions, or cannot classify specific patterns of differential change when multiple patterns exist. To address these limitations, we present a flexible and efficient method for the detection of differential epigenomic activity across multiple conditions. We utilize data from the ENCODE Consortium and show that the presented method, mixNBHMM, exhibits superior performance to current tools and it is among the fastest algorithms available, while allowing the classification of combinatorial patterns of differential epigenomic activity and the characterization of chromatin regulatory states.

## 1. Introduction

Epigenomics, the study of the human genome and its interactions with proteins and other cellular elements, has become of significant interest in recent years. Such interactions have been shown to regulate essential cellular functions such as gene expression and DNA packaging (Kim et al., 2018), resulting in downstream phenotypic impact. Therefore, the interrogation of how these interactions may change across conditions, such as cell types or treatments, is of marked interest in biomedical research. Several landmark articles have identified specific genomic regions of changing (differential) epigenomic activity between conditions as drivers of cell differentiation (Creyghton et al., 2010), cancer progression (Varambally et al., 2002), and a number of human diseases (Portela and Esteller, 2010). Within differential regions, the delineation of specific patterns of change across conditions is also of interest, for example classifying the gain-of-or loss-of-activity in genomic loci due to treatment (Clouaire et al., 2014). The identification of specific combinations of processes acting locally may also be informative, such as for segmenting the genome into regulatory states (Kundaje et al., 2015). To quantify local epigenomic activity, a common high-throughput assay is chromatin immunoprecipitation followed by massively parallel sequencing (ChIP-seq). ChIP-seq experiments begin with cross-linking DNA and proteins within chromatin structures, which are then fragmented by sonication in a particular sample. DNA fragments bound to the protein of interest are isolated by chromatin immunoprecipitation, which are then sequenced via massively parallel high-throughput sequencing to generate short sequencing reads pertaining to the original fragments. Sequences are then mapped onto a reference genome through sequence alignment to determine their likely locations of origin. Genomic coordinates containing a high density of mapped reads, often referred to as enrichment regions (peaks), indicate likely locations of protein-DNA interaction sites, and all other regions are referred to as background regions. This local read density is often summarized by counting the number of reads mapped onto non-overlapping windows of fixed length tiling the genome (window read counts), forming the basis for downstream analyses. Across multiple conditions, regions exhibiting enrichment in at least one condition, but not across all conditions, indicate the presence of differential activity pertaining to the protein-DNA interaction of interest.

To date, many differential peak callers (DPCs) have been proposed (Song and Smith, 2011; Stark and Brown, 2011; Shen et al., 2013; Chen et al., 2015; Lun and Smyth, 2015; Allhoff et al., 2016). However, several challenges affect their ability to accurately detect regions of differential activity from the wide range of ChIP-seq experiments (Section 2). First, differential regions may be both short or broad in length, causing difficulty for methods optimized for a particular type of signal profile (Stark and Brown, 2011; Chen et al., 2015). Second, methods that pool experimental replicates together (Song and Smith, 2011) often exhibit more false positive calls compared to methods that jointly model replicates from each condition (Steinhauser et al., 2016). Third, the analysis of ChIP-seq data is often subject to complex biases that may vary across the genome, as differences in local read enrichment may depend on the total read abundance in a given region. DPCs that solely rely on sample-specific global scaling factors or control subtraction methods (Stark and Brown, 2011; Shen et al., 2013; Chen et al., 2015; Allhoff et al., 2016) may be prone to detecting spurious differences due to the lack of non-linear normalization methods (Lun and Smyth, 2015). Reflecting these limitations, a recent comparison of DPCs demonstrated that current methods tend to detect either a large number of short peaks (low sensitivity) or exhibit a high number of false positive calls (low specificity) in ChIP-seq experiments with broad regions of enrichment (Steinhauser et al., 2016). Moreover, few methods are able to simultaneously test for differential activity across three or more conditions (Chen et al., 2015; Lun and Smyth, 2015), or can classify specific differential combinatorial patterns. Altogether, these limitations can impact the drawing of accurate insights from modern epigenomic studies.

Here, we propose an efficient and flexible statistical method to identify differential regions of enrichment from epigenomic experiments with diverse signal profiles and collected under common multi-replicate, multi-condition settings. Our method overcomes the limitations of current DPCs with three major features. First, it uses a hidden Markov model (HMM) to account for the diversity in differential enrichment profiles that may result from short and broad epigenomic ChIP-seq data sets. Second, it captures specific differential combinatorial patterns through a novel finite mixture model emission distribution within the HMM’s differential state. Each mixture component pertains to a particular differential combinatorial pattern that is formed by the presence or absence of local enrichment across conditions, where a generalized linear model (GLM) is used to model the specific differential combinatorial pattern while accounting for sample- and window-specific normalization factors via offsets. Third, it enables the simultaneous detection and classification of epigenomic changes under three or more conditions, a novelty not yet available in any other DPC algorithm. The presented method offers additional benefits over current HMM-based DPC algorithms (Song and Smith, 2011; Allhoff et al., 2016) that include a GLM-based framework with an embedded mixture model, which allows the modeling of covariates of interest as well as the inclusion of model offsets for non-linear normalization, and a fast and accurate parameter estimation scheme via rejection-controlled EM algorithm (RCEM).

## 2. Data

Histones are proteins that interact and condense DNA in eukaryotic cells into structural units called nucleosomes. Multiple types of enzymatic modifications may be applied to histones, resulting in changes in local DNA packaging and chromatin accessibility mediated by nucleosomes (Bannister and Kouzarides, 2011). In turn, cellular processes such as gene transcription, gene silencing, DNA repair, replication, and recombination are also affected. Proteins that interact with DNA and alter its functional properties are often referred to as epigenomic marks. For example, the trimethylation of histone H3 at lysines 36 and 27 (H3K36me3 and H3K27me3) are two types of histone modifications that tend to occur in genomic loci containing actively transcribed and repressed genes (Liu et al., 2016), respectively, and exhibit broad enrichment profiles. These marks have been investigated in cancer studies, where their absence is often observed in multiple cancer types (Wei et al., 2008). As a result, H3K36me3 and H3K27me3 are considered to be key prognostic indicators in patients with breast, ovarian, and pancreatic cancer. EZH2, a major component of the polycomb complex PRC2 that catalyzes the methylation of H3K27me3 (Margueron and Reinberg, 2011), is another example of a protein with experimental signal characterized by broad enrichment domains and co-occurs with the activity of H3K27me3.

Using ChIP-seq data pertaining to histone modifications H3K27me3, H3K36me3, and the enhancer EZH2 from the ENCODE Consortium, we find that current DPCs have difficulty in accurately detecting broad regions of differential enrichment between several common cell lines (Figure 1). In line with previous findings (Steinhauser et al., 2016), we observe that even current DPCs designed for broad data (Song and Smith, 2011; Allhoff et al., 2016) tend to detect either overly fragmented differential peaks or call regions exhibiting no difference in experimental signal between conditions as differential (Figure 1A). The low specificity and sensitivity of such methods may impair the biological interpretation of the resulting peak calls in downstream analyses. Methods that rely on candidate peaks may also exhibit a compromised performance due to the limitations of single-sample peak callers in broad data (Stark and Brown, 2011; Chen et al., 2015). In addition, most current DPCs restrict their application to the analysis of two experimental conditions. For methods that are tailored for the analysis of three or more conditions, the classification of specific differential combinatorial patterns across conditions (or across various epigenomic processes) is still an open problem. The classification of such patterns would allow researchers to, for example, quantify treatment responses on the epigenomic level (Clouaire et al., 2014), or identify sets of processes working together to regulate local chromatin state. We find that the performance of such methods exhibit low sensitivity and specificity in calling differential regions in broad marks (Figure 1B).

**Figure 1:**
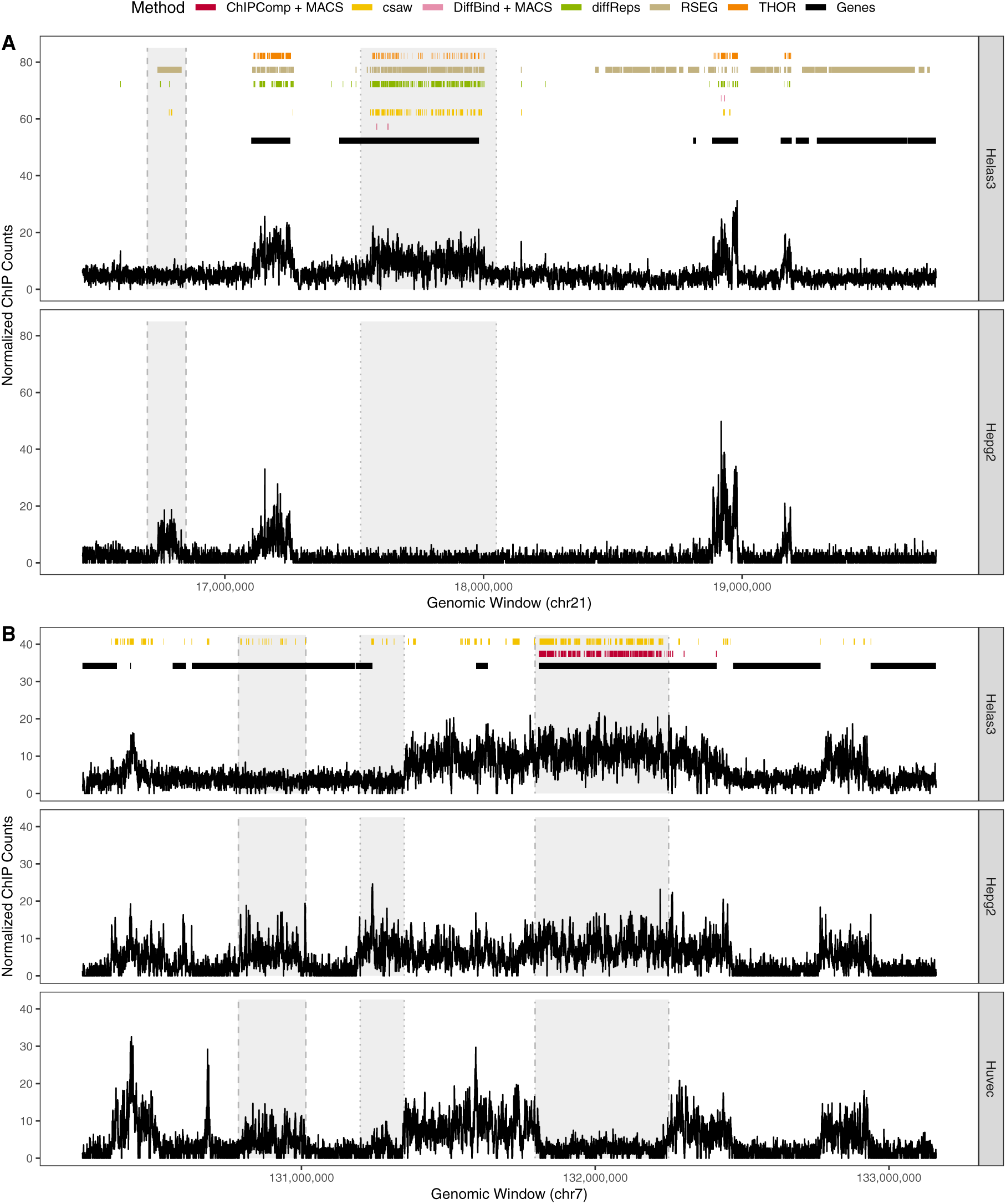
Performance of current DPC methods on calling differential enrichment regions in broad marks under a false discovery rate control of 0.05. (A): Differential peak calls between cell lines Helas3 and Hepg2 for the H3K36me3 histone modification. (B): Differential peak calls between cell lines Helas3, Hepg2, and Huvec for the H3K27me3 histone modification. Only ChIPComp, csaw, and DiffBind are designed for DPC under three or more conditions. Shaded regions indicate observed differential enrichment, and each vertical line type bordering each region represents a different combinatorial pattern of enrichment across cell lines. Optimal DPCs would call broad peaks inside shaded regions and no peaks outside them. This figure appears in color in the electronic version of this article.

We assessed the performance of our model on ChIP-seq experiments characterized by broad peaks (H3K36me3, H3K27me3, and EZH2) and short peaks (H3K27ac, H3K4me3, and the transcription factor CTCF). In simulations (Section 4) and in data sets from the ENCODE Consortium (Landt et al., 2012), we show that our model addresses the issues of the current peak callers in broad data (Section 5.1), while being flexible for short peaks (Section 5.2) and comparable to the fastest DPCs regarding the computation time. We show that our method can also be utilized for genomic regulatory state segmentation when studying multiple types of epigenomic processes from a single condition or cell line (Section 5.3). The Web Appendices A1 and A2 present the data accession codes and the data pre-processing steps, respectively. Code implementing the method and to replicate the presented results are available in the Web Appendices A3 and A4, respectively.

## 3. Methods

### 3.1 Statistical Model

Let *Y*_*hij*_ denote the random variable pertaining to the ChIP read count for genomic window *j* from sample *i* of condition *h*, where *j* = 1, …, *M, i* = 1, …, *n*_*h*_, *h* = 1, …, *G*, and let *y*_*hij*_ be the observed count. Here, *n*_*h*_ is the number of samples in condition *h* and 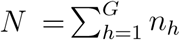 is the total number of samples across the *G* conditions. At the *j*^*th*^ window, let 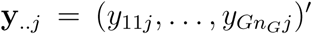 denote the *N* × 1 vector of ChIP window read counts across all samples and conditions, and let **y** = (**y**.′_.1_, …, **y**.′._*M*_)′ denote the corresponding *NM* × 1 vector of window read counts spanning all windows, samples, and conditions. We assume that each window belongs to one of three possible hidden states: consensus background (state 1), differential (state 2), and consensus enrichment (state 3). Windows exhibiting low (high) enrichment across all conditions will be modeled by an emission distribution pertaining to the consensus background (enrichment) state. Windows exhibiting enrichment under at least one condition, but not all conditions, will be modeled by an emission distribution pertaining to the differential state. If *G* conditions are of interest, there are *L* = 2^*G*^ − 2 possible differential combinatorial patterns of enrichment and background across conditions at a given window. The emission distribution pertaining to the differential state models all *L* possible differential combinatorial patterns via a mixture model with mixture proportions ***δ*** = (*δ*_1_, …, *δ*_*L*_)′, such that 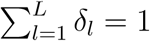 (Web Figure 1).

To model transitions between states, we assume a single latent discrete time stationary Markov chain 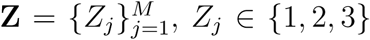, with state-to-state transition probabilities ***γ*** = (*γ*_11_, *γ*_12_, …, *γ*_33_)′ and initial probabilities ***π*** = (*π*_1_, *π*_2_, *π*_3_)′, such that 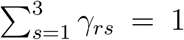 and 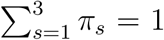 for *r* ∈ {1,2,3}. To facilitate the notation, let *f*_*r*_(**y**_..*j*_|***Ψ***_*r*_) denote the emission distribution corresponding to the *r*^*th*^ hidden state, where **Ψ** = (***π***′, ***γ***′, ***δ***′, ***Ψ***′)′ denotes the vector of all model parameters, 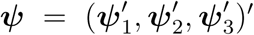 denotes each state’s set of emission distribution-specific parameters, and 𝒵 denotes the set of 3^*M*^ possible state paths of length *M*. Then, the likelihood function pertaining to the proposed HMM may be written as

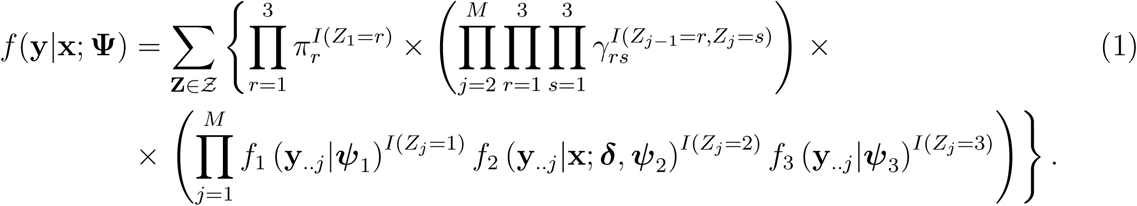

Here, **x** is a fixed *G* × *L* design matrix enumerating each of the *L* possible differential combinatorial patterns in terms of the presence or absence of enrichment across each of the *G* conditions, only in the emission distribution of the differential state.

We assume that read counts pertaining to genomic windows from the consensus background (*r* = 1) and consensus enrichment (*r* = 3) states follow a Negative Binomial (NB) distribution with state-specific parameters ***Ψ***_*r*_ = (*µ*_(*r,hij*)_, *ϕ*_*r*_)′, with mean *µ*_(*r,hij*)_ and variance *µ*_(*r,hij*)_(1 + *µ*_(*r,hij*)_*/ϕ*_*r*_). Assuming independence of read counts across experiments and samples, conditional upon the HMM state, the emission distribution of the consensus background and consensus enrichment states, respectively, can be written as

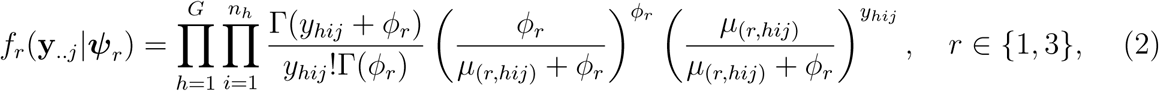

with *y*_*hij*_ ∈ {0, 1, 2, …}, such that log(*µ*_(1,*hij*)_) = *β*_1_ + *u*_*hij*_, log(*ϕ*_1_) = *λ*_1_, log(*µ*_(3,*hij*)_) = *β*_1_ + *β*_3_ + *u*_*hij*_, and log(*ϕ*_3_) = *λ*_1_ + *λ*_3_. The offset *u*_*hij*_ adjusts for technical artifacts and allows the non-linear normalization of the signal profile across genomic windows, conditions, and samples (Web Appendix B2). When *u*_*hij*_ = 0, *β*_1_ and *λ*_1_ represent the log-mean and log-dispersion, respectively, of read counts pertaining to consensus background state windows, whereas *β*_3_ and *λ*_3_ represent the difference in log-mean and log-dispersion of read counts from consensus enrichment state windows relative to consensus background state windows. For windows belonging to the differential state (*r* = 2), we assume that the corresponding read counts are modeled by a *L*-component finite mixture model with mixture components that follow a Negative Binomial distribution, where each component corresponds to a particular differential combinatorial pattern. To define these patterns, let us consider the sets *S*_1_, …, *S*_*L*_ that delineate the subset of the *G* conditions that are enriched in each of the *L* differential combinatorial patterns. For instance, if *G* = 3, the sets *S*_1_ = {1}, *S*_2_ = {2}, *S*_3_ = {3}, *S*_4_ = {1, 2}, *S*_5_ = {1, 3}, and *S*_6_ = {2, 3} define the six possible differential combinatorial patterns of enrichment and background across three conditions. That is, the set *S*_1_ denotes enrichment in only the first condition and background in all others, whereas the set *S*_6_ denotes enrichment in conditions 2 and 3 and background in condition 1. The presence or absence of enrichment in each of the *L* sets is encoded into each column of **x** = (**x**_1_, …, **x**_*L*_), such that **x**_*l*_ = (*x*_1*l*_, …, *x*_*Gl*_)′, and *x*_*hl*_ = *I*(*h* ∈ *S*_*l*_) for *l* = 1, …, *L* and *h* = 1, …, *G*. That is, **x**_*l*_ is the *G* × 1 vector of binary indicator variables denoting which subset of conditions are enriched in pattern (mixture component) *l*. A graphical illustration of our proposed model is provided in Web Figure 1.

Let ***Ψ***_2_ denote the state-specific parameter vector pertaining to the differential state and let ***Ψ***_(2,*l*)_ denote the set of parameters pertaining to the *l*^*th*^ mixture component. Assuming independence of read counts across conditions and samples, conditional upon the differential HMM state, the finite mixture model emission distribution can be written as

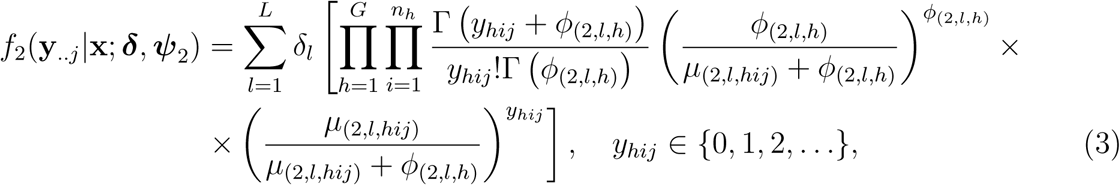

where *µ*_(2,*l,hij*)_ and *ϕ*_(2,*l,h*)_ are the mean and dispersion, respectively, pertaining to read counts originating from window *j* and sample *i* in condition *h* from the mixture component *l*. We assume that log(*µ*_(2,*l,hij*)_) = *β*_1_ + *β*_3_*x*_*hl*_ + *u*_*hij*_ and log(*ϕ*_(2,*l,h*)_) = *λ*_1_ + *λ*_3_*x*_*hl*_. That is, in the mixture component *l*, we utilize the same consensus background (consensus enriched) log-mean and log-dispersion from (2) in all conditions that are specified by **x**_*l*_ to be background (enriched) in the *l*^*th*^ differential combinatorial pattern. There are several advantages to such a parametrization for the differential emission distribution. For example, it ensures that windows exhibiting differential enrichment across conditions share means and dispersions that are common between the consensus background and consensus enrichment states, a reasonable assumption that significantly increases computational efficiency. Utilizing a mixture model as the differential state emission distribution avoids the computational burden that would come from assuming separate hidden states for each of the *L* differential combinatorial patterns, particularly as *G* increases. We evaluate the strength of these assumptions through multiple simulations and a real data benchmarking analysis in Sections 4 and 5.

Two novel features result from our proposed approach that are relevant to the context of differential enrichment detection from ChIP-seq experiments. By using a modified version of the Expectation-Maximization (EM) algorithm to estimate the model parameters, we are able not only to detect differential enrichment regions across multiple conditions, but we can also classify various differential combinatorial patterns of enrichment within broad and short differential enrichment domains. With state-specific parameters, the current implementation of the method allows the direct modeling of continuous covariates (e.g. input controls; Web Appendix A3), for which a state-level testing of their effects on the read count distribution could be performed. In a simulation study and in real data analyses, however, we did not observe a significant improvement in performance in differential peak detection after accounting for the effect of input controls (Web Appendix B2), a fact that has also been observed by others (Lun and Smyth, 2015).

### 3.2 Estimation

To simplify the parameter estimation in (4), we introduce another set of latent variables 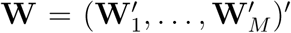, such that **W**_*j*_ = (*W*_*j*1_, …, *W*_*jL*_)′ for *j* = 1, …, *M*. We assume that **W** is a sequence of independent random vectors such that **W**_*j*_|(*Z*_*j*_ = 2) ∼ Multinomial(1, ***δ***) and **W**_*j*_|(*Z*_*j*_ = *r*) = 0 with probability 1 if *r* = {1, 3}. Under this setup, one may define the data generating mechanism when *Z*_*j*_ = 2 (differential state) and *W*_*jl*_ = 1 (*l*^*th*^ differential combinatorial pattern) such that read counts pertaining to genomic window *j* are sampled from *f*_(2,*l*)_ given ***Ψ***_(2,*l*)_ and **x**_*l*_. Let 𝒲 denote the set of *L*^*M*^ possible combinations of latent vectors **W**. Hence, the likelihood function of the observed data (1) can be rewritten as

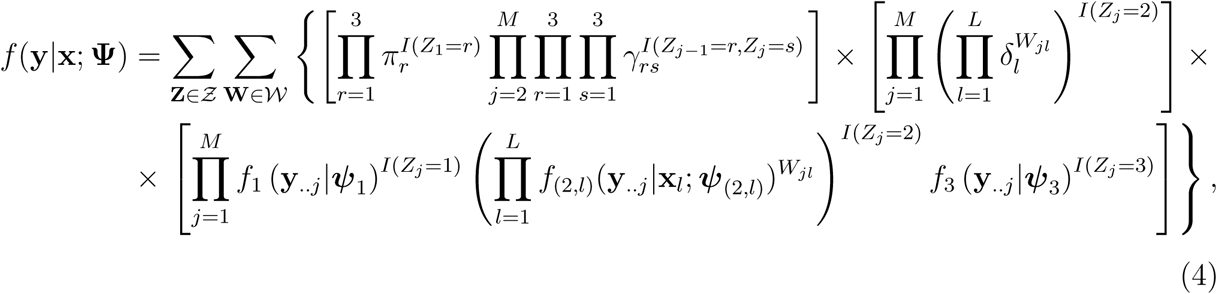

where *f*_(2,*l*)_(**y**_..*j*_|**x**_*l*_; ***Ψ***_(2,*l*)_) is defined as in (3). In the *t*^*th*^ step of the EM algorithm, the *Q* function of the complete data log-likelihood can be written as

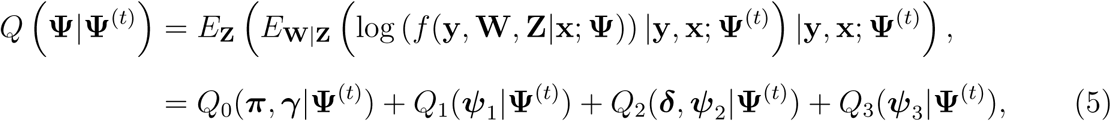

where *Q*_0_(***π, γ***|**Ψ**^(*t*)^), *Q*_1_(***ψ***_1_|**Ψ**^(*t*)^), *Q*_2_(***δ, Ψ***_2_|**Ψ**^(*t*)^), and *Q*_3_(***ψ***_3_|**Ψ**^(*t*)^) are defined in (A.1). In the E-step of the EM algorithm, we compute the posterior probabilities from (5). The quantities (*Pr Z*_*j*_ = *r*|**y, x**; **Ψ**^(*t*)^) and *Pr* (*Z*_*j*−1_ = *r, Z*_*j*_ = *s*|**y, x**; **Ψ**^(*t*)^), defined in (A.2), can be calculated through the Forward-Backward algorithm (see Appendix) and 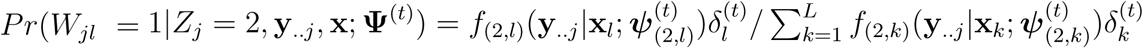 for *l* = 1, …, *L*.

The *Q* function is maximized with respect to the parameters **Ψ** = (***π***′, ***γ***′, ***δ***′, *β*_1_, *β*_3_, *λ*_1_, *λ*_3_)^′^ during the M-step of the algorithm. Estimates for the initial and transition probabilities can be directly calculated as 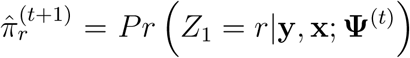 and 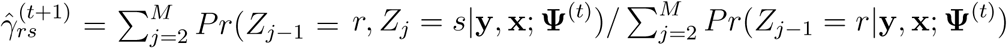, respectively, restricted to 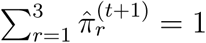 and 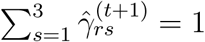, for *r* ∈ {1, 2, 3}. We perform conditional maximizations to compute estimates of the remaining model parameters (***δ***′, *β*_1_, *β*_3_, *λ*_1_, *λ*_3_)′. First, mixture proportions can be estimated as 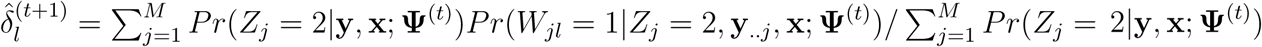. Estimating (*β*_1_, *β*_3_, *λ*_1_, *λ*_3_)′ from (5) can be seen as obtaining parameter estimates from a series of weighted NB regression models with shared mean and dispersion parameters. We jointly estimate these quantities via the algorithm BFGS (Fletcher, 2013).

The estimation scheme is robust to situations where certain differential combinatorial patterns of enrichment are rare (Figure 5). This unique characteristic results from the fact that ChIP-seq experiments often provide enough data (usually *M >* 10^7^ non-overlapping windows of 250 bp fixed size for the human reference genome) to estimate the parameters (*β*_1_, *β*_3_, *λ*_1_, *λ*_3_)′, which are shared across all *L* mixture components and HMM states. If pruning differential combinatorial patterns of the differential mixture component is of interest, the optimal number of mixture components *L*^*^, *L*^*^ *< L*, can be selected via the Bayesian Information Criterion (BIC) for HMMs. We observed that selecting the optimal number of mixture components based on BIC agrees with the pruning of rare differential combinatorial patterns that we would not biologically expect to observe in real data (Web Appendix B3).

**Figure 2:**
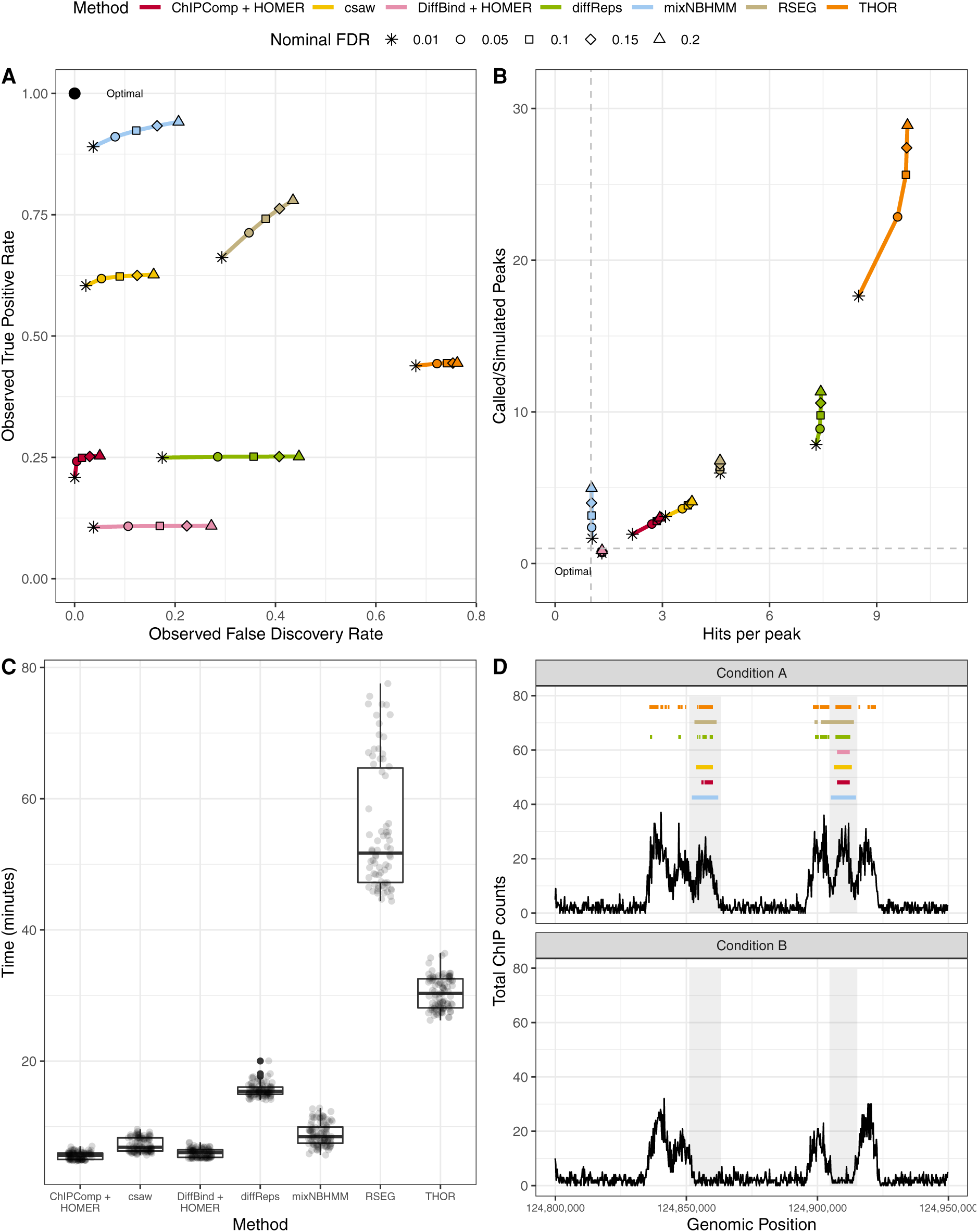
Sequencing read-based simulation from the csaw pipeline. (A): average observed sensitivity and FDR for various methods. (B): scatter plot of average ratio of called and simulated peaks (y-axis) and number of called peaks intersecting true differential regions (x-axis). (C): box plot of computing time (in minutes) for various algorithms. (D): an example of differential peak calls under a nominal FDR control of 0.05. Shaded areas indicate true differential peaks. This figure appears in color in the electronic version of this article.

**Figure 3:**
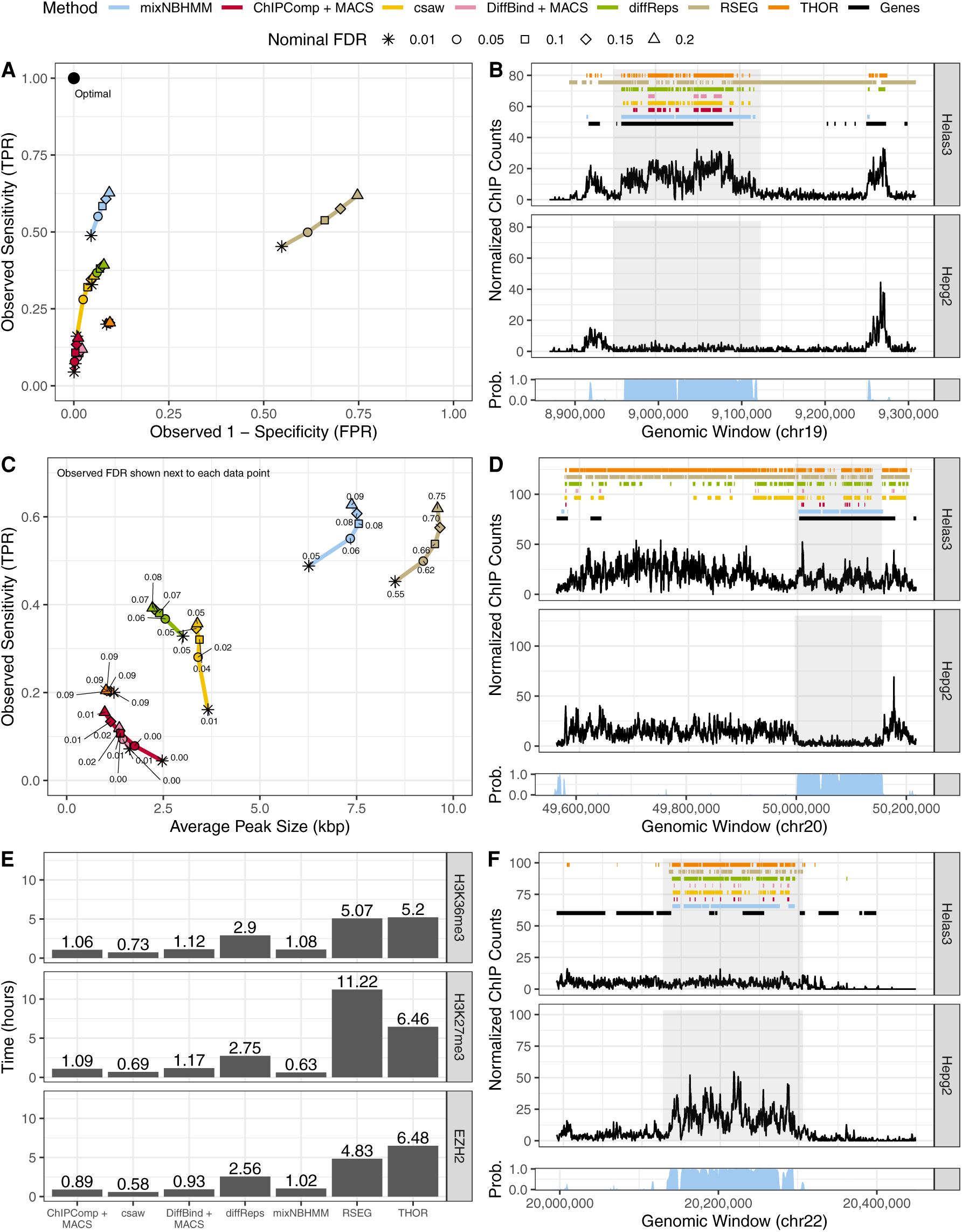
Analysis of broad ENCODE data. (A): ROC curves of H3K36me3 differential peak calls. (C): sensitivity (y-axis) and average H3K36me3 differential peak size (kbp; x-axis) of various methods under different nominal FDR thresholds (observed FDR annotated next to data points). (B), (D), and (F): example of peak calls from H3K36me3, H3K27me3, and EZH2, respectively, under a nominal FDR control of 0.05. Posterior probabilities of the HMM differential state are shown at the bottom of each panel. (E): computing time of genome-wide analysis from various methods. This figure appears in color in the electronic version of this article.

**Figure 4:**
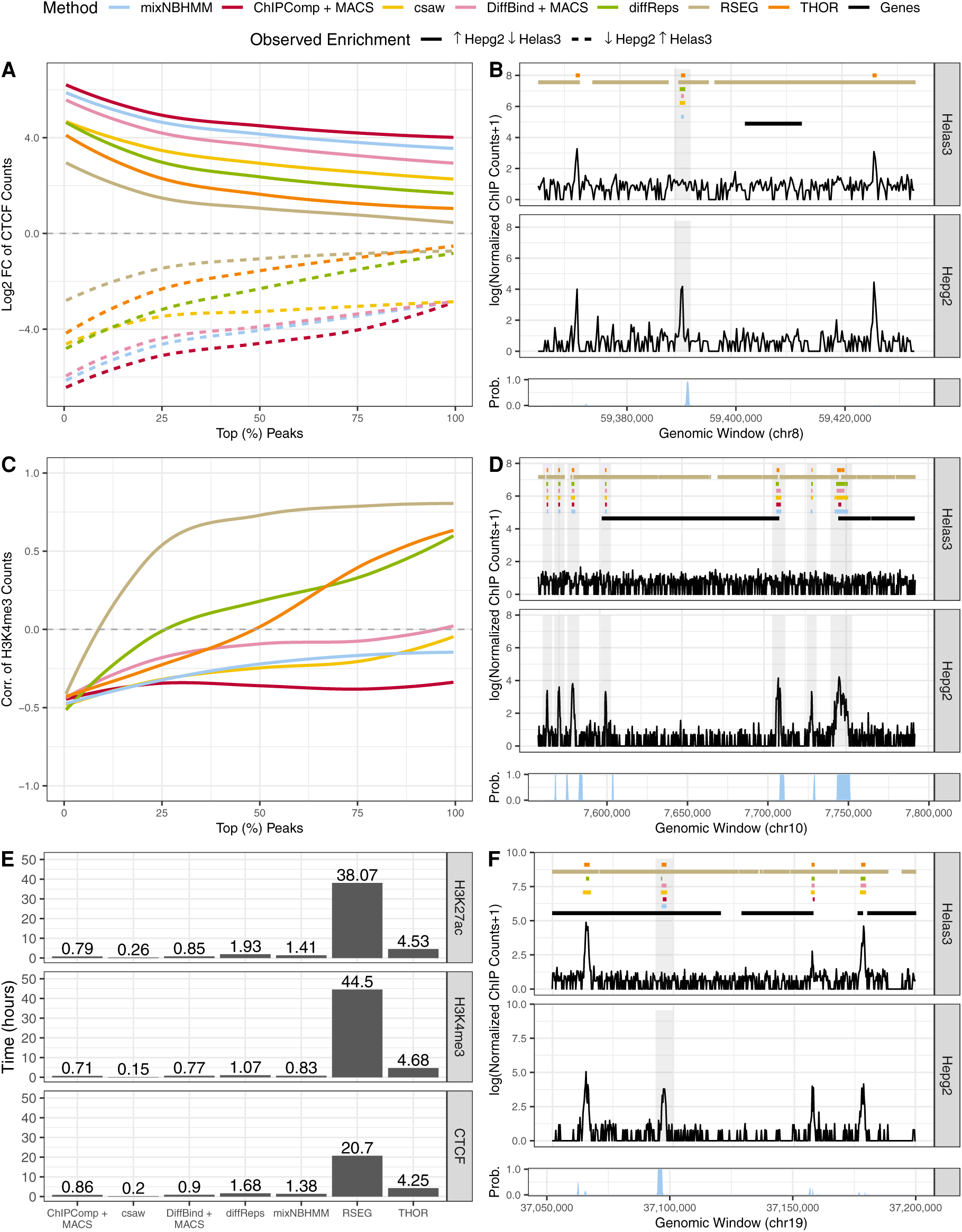
Analysis of short ENCODE data. (A) and (C): median LFC and correlation between cell lines of ChIP-seq counts from differential peaks for CTCF and H3K4me3, respectively. (B), (D), and (F): example of peak calls from CTCF, H3K4me3, and H3K27ac, respectively. Posterior probabilities of the HMM differential state are shown at the bottom of each panel. (E): computing time of genome-wide analysis from various methods. Results are shown under a nominal FDR control of 0.05. This figure appears in color in the electronic version of this article.

**Figure 5:**
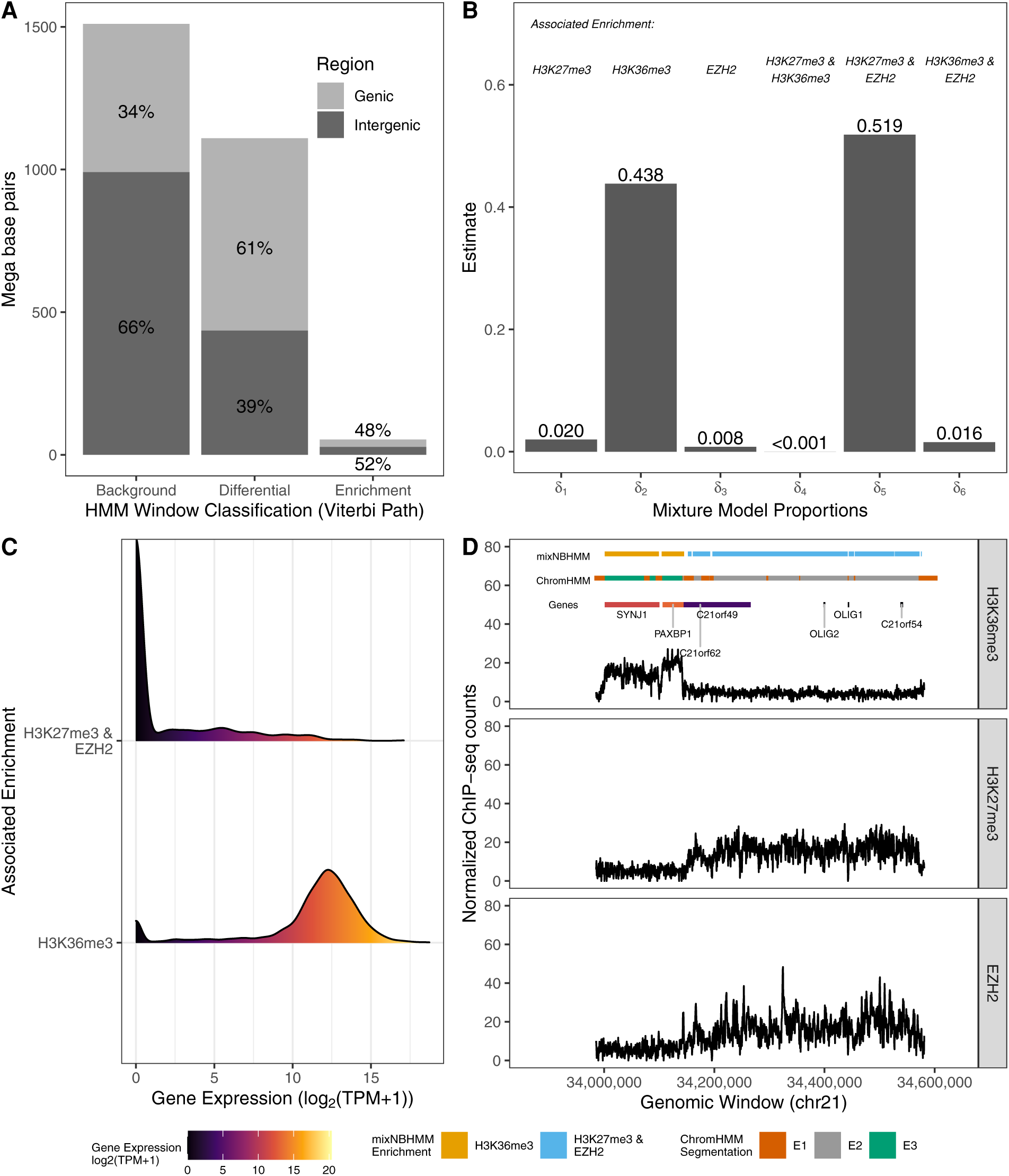
Genomic chromatin state segmentation and classification. (A): distribution of base pairs (y-axis) and the Viterbi sequence of states (x-axis). (B): estimated mixture probabilities and the associated differential combinatorial patterns. (C): density estimate from expression of genes intersecting differential peaks associated with the enrichment of H3K36me3 alone or the enrichment of H3K27me3 and EZH2 in consensus. (D): example of a genomic region with differential peaks and genes, colored according to their classification and expression levels, respectively. This figure appears in color in the electronic version of this article.

To obtain the parameter estimates 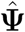, the EM algorithm iterates until the maximum absolute relative change in the parameter estimates three iterations apart is less than 10^−3^ for three consecutive iterations. To reduce the computation time, we make use of a RCEM algorithm with threshold 0.05. Briefly, the RCEM algorithm substantially reduces the dimensionality of the data during the M-step by randomly assigning a zero posterior probability to genomic windows unlikely to belong to each of the HMM states. The current estimation set up allows genomic windows exhibiting equal distribution of read counts to have their posterior probability aggregated during the M-step of the algorithm. Often, the distribution of read counts along the genome is highly concentrated on a particular set of values, such as 0, 1, and 2 for instance. Genomic windows exhibiting a particular pattern of counts across samples and conditions can have their posterior probability aggregated during the M-step, which further reduces the dimensionality of the objective function during the numerical optimization and leads to a fast gradient-based optimization.

Once the algorithm reaches convergence, the final set of HMM posterior probabilities can be used to segment the genome into consensus background, differential, or consensus enrichment windows. Approaches that control the total false discovery rate (FDR) via posterior probabilities (Efron et al., 2001) or that estimate the most likely sequence of hidden states (Viterbi, 1967) can be used for such purposes. Let 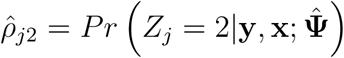 denote the estimated posterior probability that the *j*^*th*^ genomic window belongs to the differential HMM state, *j* = 1, …, *M*. For a cutoff of posterior probability *α*, the total FDR is 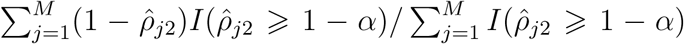, where *I*(·) is an indicator function. The posterior probability cutoff is then chosen by controlling the total FDR. Differential regions of enrichment are formed by merging adjacent windows that either meet a given FDR threshold level for the differential HMM state or belong to the same Viterbi’s predicted state. A discussion about these two approaches under the proposed model is presented in the Web Appendix B4. Additional details of proposed EM algorithm and the implemented code are available in the Web Appendices B5 and A3, respectively.

## 4. Simulation Studies

We evaluate the presented model in two independent simulation studies of broad epigenomic marks. In the first study (Section 4.1), we simulated read count-based data to assess the precision of the parameter estimation scheme, the performance of differential peak detection, and the accuracy of the classification of specific differential combinatorial patterns of enrichment within differential peaks. In the second simulation study (Section 4.2), we utilize the simulation pipeline presented in Lun and Smyth (2015) to generate synthetic ChIP-seq reads from *in silico* experiments with broad differential peaks. The aim of the second simulation study was to compare our model with other DPCs in a more realistic scenario with broad peaks, while also avoiding the assumption of a parametric model for the data.

### 4.1 Read Count Simulation

Read counts were simulated under different scenarios that varied regarding the type of histone modification mark (H3K36me3 and H3K27me3), genome length (*M*, 10^5^, 5 × 10^5^, and 10^6^ windows), number of conditions (*G*, 2, 3, and 4), and number of replicates per condition (*n*, 1, 2, and 4). We further assessed our model under different Signal-to-Noise Ratio (SNR) levels. We define the SNR as the ratio between the means of consensus enrichment and consensus background emission distributions. Mean and dispersion parameters used in this simulation study were estimated from ENCODE data and are presented in the Web Appendix C1 for all the scenarios. Different SNR levels were defined by decreasing the ratio of the means in decrements of 10% while maintaining the mean-variance relationship. Read counts were assumed to follow a NB distribution and were simulated using a first-order Markov chain with 2^*G*^ states, representing every combination of background and enrichment across *G* conditions. We aimed to assess whether our model was able to assign all 2^*G*^ − 2 simulated differential states to the differential HMM state, while maintaining a precise parameter estimation scheme and accurate classification of differential combinatorial patterns.

#### 4.1.1 Simulation Results

Table 1 shows the true values and the average relative bias of parameter estimates (and the 2.5^*th*^, 97.5^*th*^ percentiles) from a hundred simulated data sets relative to the scenario of H3K27me3 with 10^5^ genomic windows (see Web Appendix C1 for additional details). Results are shown for different levels of SNR, number of conditions, and number of replicates per condition. Overall, no significant differences regarding the relative bias of parameter estimates were observed across simulations under different genome lengths. Depending on the number of conditions, the observed relative bias and the range of the reported percentiles tended to decrease as more replicates were included in the analyses. This effect was particularly significant in scenarios with four conditions with respect to parameters *β*_3_ and *λ*_3_. In general, scenarios with higher SNR showed lower relative bias and variability of the parameter estimates in comparison to scenarios with lower SNR, regardless of the number of conditions or replicates per condition. In scenarios with lower SNR levels or higher number of conditions, these results also highlight the importance of experimental replicates to achieve precise parameter estimates. The proposed estimation approach via EM algorithm led to precise parameter estimates and was robust to a data generating mechanism that was different than the one assumed by the proposed model.

**Table 1:** Read count simulation. True values and average relative bias of parameter estimates (and 2.5^*th*^, 97.5^*th*^ percentiles) across a hundred simulated data sets are shown for H3K27me3 with 10^5^ genomic windows. Scenarios with observed SNR and 70% of observed SNR are shown.

| SNR | Conditions | Parameter | True | One Replicate |  | Two Replicates |  | Four Replicates |  |
| --- | --- | --- | --- | --- | --- | --- | --- | --- | --- |
| | | | | R. Bias | $(P_{2.5}, P_{97.5})$ | R. Bias | $(P_{2.5}, P_{97.5})$ | R. Bias | $(P_{2.5}, P_{97.5})$ |
| 70% of<br>Observed<br>SNR | Two | $\beta_1$ | 1.116 | 0.000 | (-0.004, 0.004) | 0.000 | (-0.002, 0.002) | 0.000 | (-0.002, 0.002) |
| | | $\beta_3$ | 0.808 | -0.001 | (-0.012, 0.008) | 0.000 | (-0.005, 0.006) | 0.000 | (-0.003, 0.004) |
| | | $\lambda_1$ | 1.281 | 0.000 | (-0.016, 0.013) | 0.000 | (-0.010, 0.010) | 0.000 | (-0.008, 0.007) |
| | | $\lambda_3$ | -0.232 | -0.001 | (-0.109, 0.110) | 0.000 | (-0.078, 0.067) | 0.000 | (-0.058, 0.048) |
| | Three | $\beta_1$ | 1.116 | 0.002 | (-0.005, 0.013) | -0.003 | (-0.007, 0.001) | -0.001 | (-0.002, 0.001) |
| | | $\beta_3$ | 0.808 | -0.156 | (-0.233, -0.098) | -0.014 | (-0.025, -0.005) | -0.001 | (-0.004, 0.003) |
| | | $\lambda_1$ | 1.281 | 0.005 | (-0.033, 0.032) | 0.004 | (-0.006, 0.014) | 0.001 | (-0.006, 0.007) |
| | | $\lambda_3$ | -0.232 | 0.898 | (0.679, 1.083) | 0.130 | (0.012, 0.242) | 0.015 | (-0.033, 0.061) |
| | Four | $\beta_1$ | 1.116 | 0.001 | (-0.005, 0.020) | -0.012 | (-0.017, -0.008) | -0.010 | (-0.014, -0.007) |
| | | $\beta_3$ | 0.808 | -0.145 | (-0.256, -0.120) | -0.087 | (-0.098, -0.075) | -0.032 | (-0.062, -0.015) |
| | | $\lambda_1$ | 1.281 | 0.016 | (-0.028, 0.033) | 0.028 | (0.016, 0.041) | 0.017 | (0.007, 0.028) |
| | | $\lambda_3$ | -0.232 | 0.947 | (0.819, 1.083) | 0.769 | (0.672, 0.877) | 0.366 | (0.211, 0.613) |
| Observed<br>SNR | Two | $\beta_1$ | 1.116 | 0.000 | (-0.004, 0.004) | 0.000 | (-0.003, 0.002) | 0.000 | (-0.002, 0.002) |
| | | $\beta_3$ | 1.165 | 0.000 | (-0.005, 0.005) | 0.000 | (-0.003, 0.003) | 0.000 | (-0.002, 0.003) |
| | | $\lambda_1$ | 1.281 | 0.000 | (-0.015, 0.018) | -0.001 | (-0.010, 0.008) | 0.001 | (-0.004, 0.007) |
| | | $\lambda_3$ | 0.124 | -0.012 | (-0.231, 0.174) | 0.008 | (-0.108, 0.129) | -0.008 | (-0.107, 0.085) |
| | Three | $\beta_1$ | 1.116 | -0.004 | (-0.008, 0.001) | -0.001 | (-0.003, 0.002) | 0.000 | (-0.002, 0.002) |
| | | $\beta_3$ | 1.165 | -0.010 | (-0.019, -0.002) | 0.000 | (-0.003, 0.003) | 0.000 | (-0.002, 0.002) |
| | | $\lambda_1$ | 1.281 | -0.001 | (-0.012, 0.012) | 0.000 | (-0.008, 0.009) | 0.000 | (-0.006, 0.007) |
| | | $\lambda_3$ | 0.124 | -0.422 | (-0.697, -0.068) | -0.026 | (-0.171, 0.083) | -0.003 | (-0.085, 0.081) |
| | Four | $\beta_1$ | 1.116 | -0.013 | (-0.019, -0.008) | -0.008 | (-0.014, -0.003) | 0.000 | (-0.002, 0.001) |
| | | $\beta_3$ | 1.165 | -0.111 | (-0.121, -0.101) | -0.005 | (-0.010, -0.001) | 0.000 | (-0.002, 0.002) |
| | | $\lambda_1$ | 1.281 | -0.011 | (-0.025, 0.001) | 0.009 | (-0.002, 0.018) | 0.001 | (-0.004, 0.006) |
| | | $\lambda_3$ | 0.124 | -3.526 | (-3.782, -3.239) | -0.438 | (-0.740, -0.217) | -0.009 | (-0.084, 0.059) |

### 4.2 Sequencing Read Simulation

We performed a second simulation study aiming to compare the proposed model with the current DPCs ChIPComp, csaw, DiffBind, diffReps, RSEG, and THOR. We used the simulation pipeline presented by Lun and Smyth (2015) where data were generated in a more general scheme without a particular read count model assumption. Here, sequencing reads from broad ChIP-seq experiments were generated for two conditions and two replicates per condition. For the differential peaks callers ChIPComp and DiffBind that require sets of candidate regions, we followed the analyses presented by Lun and Smyth (2015) and called peaks in advance using HOMER. Peaks were then used as input in the respective software for differential call. A hundred simulated data sets were generated and peaks were called by all the methods under multiple nominal FDR thresholds. For our method and RSEG, window-based posterior probabilities were used to control the total FDR as described in Section 3.2.

#### 4.2.1 Simulation Results

Figure 2 shows the main results of our second simulation study. Out of 100 simulated data sets, RSEG either failed to analyze the data due to internal errors or called the entire genome as differential in 26 and in 3 instances, respectively. Similar issues have been previously reported in other studies (Starmer and Magnuson, 2016). We observed that our method showed the highest observed sensitivity among all DPCs, regardless of the nominal FDR thresholding level, while maintaining a moderate observed FDR (Figure 2A). Methods such as diffReps, RSEG, and THOR showed higher observed FDR levels than the nominal threshold due to the excessive number of differential peaks called outside true differential regions (shaded area in Figure 2D). While diffReps and THOR called an excessive number of short and discontiguous peaks, RSEG called regions that were usually wider than the observed differential enrichment regions. These results are further illustrated in Figure 2B, where we present the average ratio of the number of called and simulated peaks (y-axis) and the average number of called peaks intersecting true differential regions (x-axis). Regarding the computation time, the HMM-based algorithms RSEG and THOR appeared to be the most computationally intensive and required longer amounts of time to analyze the data. In Figure 2C, we present the box plots of computing time (in minutes) across a hundred simulated data sets for all benchmarked methods. While still being an HMM-based algorithm, our method was among the fastest tools for differential peak detection due to the implemented strategies to improve the computation time of the EM algorithm (Section 3.2). Figure 2D shows an example of a genomic region with simulated data and called peaks from various methods using nominal FDR threshold 0.05. As shown, our method was able to consistently cover most of true differential regions with broad peaks while exhibiting a limited number of false discoveries (see Web Appendix C2 for additional results).

## 5. Application to ENCODE Data

We applied our method to ChIP-seq data from the ENCODE Consortium (Section 2) to detect differential regions of enrichment of several epigenomic marks across distinct cell lines. First, we analyzed broad data from the histone modifications H3K36me3 and H3K27me3 as well as data from the enhancer EZH2 (Section 5.1). Secondly, we assessed the performance of the presented model on ChIP-seq experiments from the transcription factor CTCF and the histone modifications H3K27ac and H3K4me3 characterized by short peaks (Section 5.2). For H3K27ac and H3K4me3, enrichment peaks are usually deposited on the promoter regions of actively transcribed genes and several studies have associated their role with gene transcription (Creyghton et al., 2010; Lauberth et al., 2013). The transcription factor CTCF is a protein that binds to short DNA motifs and is responsible for several cellular processes that include the regulation of the chromatin 3D structure and mRNA splicing (Shukla et al., 2011). Two technical replicates for each epigenomic mark were used in the analysis. Using RNA-seq data, we assessed the practical significance of our results by associating the detection and classification of differential combinatorial patterns from called peaks of H3K36me3, H3K27me3, and EZH2 with the direction of gene expression (Section 5.3; see Web Appendix D1 for the RNA-seq data processing).

We compared the genome-wide performance of the presented model with the current DPCs ChIPComp, csaw, DiffBind, diffReps, RSEG, and THOR. For the methods that require a set of candidate regions to be specified *a priori*, ChIPComp and DiffBind, peaks were called in advance using MACS (Zhang et al., 2008) and used as input to the software for differential call. We benchmarked methods regarding the coverage of differentially transcribed gene bodies, the number and average size of differential peak calls, log_2_ fold change (LFC) of read counts, Spearman correlation of log_2_-transformed read counts between cell lines, and computation time. Metrics for sensitivity and specificity were defined on the window level and based on the coverage of differentially transcribed gene bodies by called peaks (Web Appendix D1). For broad marks, read counts were computed using non-overlapping windows of 500bp. For the remaining short marks, we computed read counts using non-overlapping windows of 250bp. Results presented in this section pertain to the analysis of two cell lines, namely Helas3 and Hepg2. A discussion about the choice of the window size is presented in the Web Appendix D2. Results from the analysis of more than two cell lines are presented in the Web Appendix D3. Data accessing code, data pre-processing steps, method-specific parameters, and code to replicate the presented results are detailed in the Web Appendix A.

### 5.1 Analysis of ChIP-seq Data From Broad Marks

Methods were benchmarked regarding the coverage of differentially transcribed gene bodies. The histone modification H3K36me3 is known to be associated with gene transcription and enriched regions of this mark are usually deposited on transcribed gene bodies. Hence, the location of differential peaks of H3K36me3 is expected to agree with the location of differentially expressed genes. Following the analysis presented by Steinhauser et al. (2016), we defined a set of protein coding genes exhibiting |LFC| *>* 2 of ChIP-seq read counts between the two analyzed cell lines as true differentially transcribed genes. Results using different threshold levels are presented in the Web Appendix D1 and agree with those presented here. Protein-coding genes with total read count across cell lines under the 25^*th*^ percentile were excluded from the analysis. Normalization by the median log-ratios of each replicate over the geometric mean was performed to avoid spurious differences due to sequencing depth.

In Figure 3A, we show receiver operating characteristic (ROC) curves for various methods and different nominal levels of FDR threshold for differential peaks of H3K36me3. Similar to the analysis of broad histone modification marks presented by Xing et al. (2012), we computed the observed true positive rates (false positive rates) on the window-level as the proportion of windows called as differential out of the total number of windows associated (not associated) with differentially transcribed genes (Web Appendix D1). Our method had the best overall performance among all DPCs as its differential peaks were able to cover most of differentially transcribed gene bodies while still maintaining a low number of false positives. Methods that tended to call short peaks, such as ChIPComp and DiffBind, were the ones with the lowest sensitivity among all methods. ChIPComp and DiffBind have been previously shown to be dependent on the set of candidate peaks and to perform best in scenarios with short peaks (Figure 3B; Steinhauser et al. (2016); Lun and Smyth (2015)). In Figure 3C, we show the observed sensitivity (y-axis) and the average differential peak size (kbp; x-axis) for various methods under different nominal FDR levels (the observed FDR is annotated next to each data point). Our model and RSEG, two HMM-based methods, tended to call broader differential peaks and exhibited better sensitivity than other methods. Yet, differential peaks called by RSEG often did not correspond to differential regions of enrichment (Figure 3D), a behavior that has been noted by others (Starmer and Magnuson, 2016) and also seen in simulated data (Figure 2). Our HMM-based method with a non-linear normalization scheme via model offsets allowed us to maintain a low observed FDR and a higher sensitivity than other DPCs. In Figure 3F, we show examples of differential peak calls for the enhancer EZH2. Our method was among the fastest algorithms due to our computational scheme, taking approximately 1 hour to analyze genome-wide data (Figure 3E).

### 5.2 Analysis of ChIP-seq Data From Short Marks

We further evaluated the performance of the proposed method on data sets characterized by short peaks, namely the histone modifications H3K4me3 and H3K27ac and the transcription factor CTCF. The goal of our analysis was to assess whether our method was robust to different types of data and still able to call short differential regions of enrichment. In these scenarios, differential peaks are usually observed in isolated genomic regions and exhibit a high SNR. It has been shown that certain HMM-based approaches, including RSEG, have low accuracy under short histone modification marks and TF data (Hocking et al., 2016). As we show, the model proposed in this article performs comparably to the evaluated DPCs known to perform best in short data (ChIPComp, DiffBind; Steinhauser et al. (2016)) and appeared to be more efficient regarding the computation time in certain scenarios.

We calculated the LFC and the Spearman correlation between cell lines Helas3 and Hepg2 based on ChIP-seq read counts mapped onto differential peaks called by each method. Read counts were previously normalized by the median log-ratios of each replicate over the geometric mean to avoid spurious differences due to sequencing depth. As these marks are characterized by short peaks, ideal methods would show high absolute LFC and negative correlation between read counts mapped on differential peaks. Figure 4 shows the main results from our analysis using short data sets. In Figures 4 A and C we show the median LFC and the Spearman correlation of ChIP-seq counts for differential CTCF and H3K4me3 peak calls (sorted by the absolute LFC), respectively, under a nominal FDR control of 0.05. We present separate curves regarding the signal of observed enrichment to better characterize the direction of change. The results show that the HMM-based methods RSEG and THOR were among those with the lowest absolute LFC among all methods, which confirms their sub optimal performance in the scenario of short peaks (Hocking et al., 2016). In addition, we observed that ChIPComp had the best performance overall as it was able to call differential peaks with the highest absolute LFC and the lowest correlation between read counts of the two analyzed cell lines. Our model was able to properly call truly short differential peaks (Figures 4 B, D, and F) and was comparable to the non-HMM based methods regarding the computation time (Figure 4E), jointly calling differential peaks in less than 1.5 hour.

### 5.3 Genomic Segmentation and Classification of Chromatin States

Lastly, we analyzed data from the cell line Helas3 to segment its genome regarding the activity of marks H3K36me3, H3K27me3, and EZH2. We considered each mark as a separate experimental condition (*G* = 3) and sought to jointly classify local chromatin states in Helas3 based upon the presence or absence of enrichment from each mark. It is known that EZH2 catalyzes the methylation of H3K27me3, a repressive mark, and H3K36me3 is associated with transcribed genes (Section 2). Hence, we expected regions of enrichment in consensus for these marks to be rare and differential regions to be mostly represented by either transcribed chromatin states (enrichment for H3K36me3 alone) or repressed chromatin states (enrichment co-occurrence for H3K27me3 and EZH2). The analyses presented in this section highlight the applicability of our method in the context of genomic segmentation (Ernst and Kellis, 2012), a distinct problem not tackled by current DPCs.

First, we segmented the genome using the Viterbi sequence of most likely HMM states to understand the distribution of genomic regions associated with consensus background, differential, and consensus enrichment states (Figure 5). While the majority of the genomic regions exhibited no enrichment for any of the analyzed marks, regions of consensus enrichment were rare and represented only 2% of the analyzed genome (Figure 5A), as expected. Consensus background and differential regions mostly covered intergenic (66%) and protein-coding genic regions (61%), respectively. Differential genomic windows were mostly representing either transcribed chromatin states or repressed chromatin states (Figure 5B). All differential combinatorial patterns expected to be rare had associated mixture proportion estimates less than 0.02 (see Web Appendix B3 for a discussion on pruning rare states).

These results suggest that protein-coding genes overlapping differential regions should be either silenced (e.g. genes associated with repressed chromatin states) or highly expressed (e.g. genes associated with transcribed chromatin states). To assign combinatorial patterns to differential peaks, we chose the combination pertaining to the most frequent mixture component across windows by using the maximum estimated mixture model posterior probability, *Pr*(*W*_*jl*_ = 1|*Z*_*j*_ = 2, **y**_..*j*_, **x**; **Ψ**^(*t*)^), *j* = 1, …, *M*. For genes overlapping differential regions associated with either transcribed or repressed chromatin states, we computed the distribution of transcripts per million (TPM) from matching RNA-seq experiments. Genes associated with transcribed chromatin states had a significantly higher distribution of TPM counts than genes associated with repressed chromatin states (Figure 5C). We detected broad differential regions of enrichment and the classification of differential combinatorial patterns agreed with their biological roles as well as the expression levels of associated gene bodies. Figure 5D shows an example of a genomic region with differential enrichment for the analyzed marks. We compare our results to ChromHMM with 3 states, a method developed for chromatin segmentation. Our method offers the benefit of simultaneously detecting differential peaks and classifying the combinatorial pattern of enrichment through mixture model posterior probabilities even in the context of genomic segmentation with highly diverse epigenomic marks (see Web Appendix D4). By using the BIC for model selection (Web Appendix B3), one can choose the number of biologically relevant mixture components to be included in the model, a task that may not be as straightforward in methods such as ChromHMM (see Supplementary Figure 4 in Ernst and Kellis (2012)).

## 6. Discussion

We presented a flexible and efficient statistical model designed to call differential regions of enrichment from ChIP-seq experiments with multiple replicates and multiple conditions. Our model has three main advantages over current methods tailored for differential peak detection. First, it uses an HMM-based approach that accounts for the local dependency of ChIP-seq counts and is able to precisely detect broad and short differential regions of enrichment. Second, it utilizes a GLM-based framework with model offsets that account for potential non-linear biases in the data as well as a constrained parametrization across HMM states and mixture components. Our implementation of the RCEM algorithm led to genome-wide analyses of ChIP-seq data under a computational time comparable to some of the fastest current methods and was at least 5 times faster than current HMM-based algorithms. Lastly, our method allows the simultaneous detection and classification of differential combinatorial patterns of enrichment from its embedded mixture model and the associated posterior probabilities under any number of conditions. Our software has been implemented into an R package and is available for download (Web Appendix A).

## Appendix

The *Q*-function of the EM algorithm is defined as *Q(***Ψ**|**Ψ**^(*t*)^) = *Q*_0_(***π, γ***|**Ψ**^(*t)*^) +*Q*_1_ (***ψ***_1_|**Ψ**^(*t*)^)+ *Q*_2_ (***δ, Ψ***_2_|**Ψ**^(*t*)^)+ *Q*_3_ (***ψ***_3_|**Ψ**^(*t*)^), such that

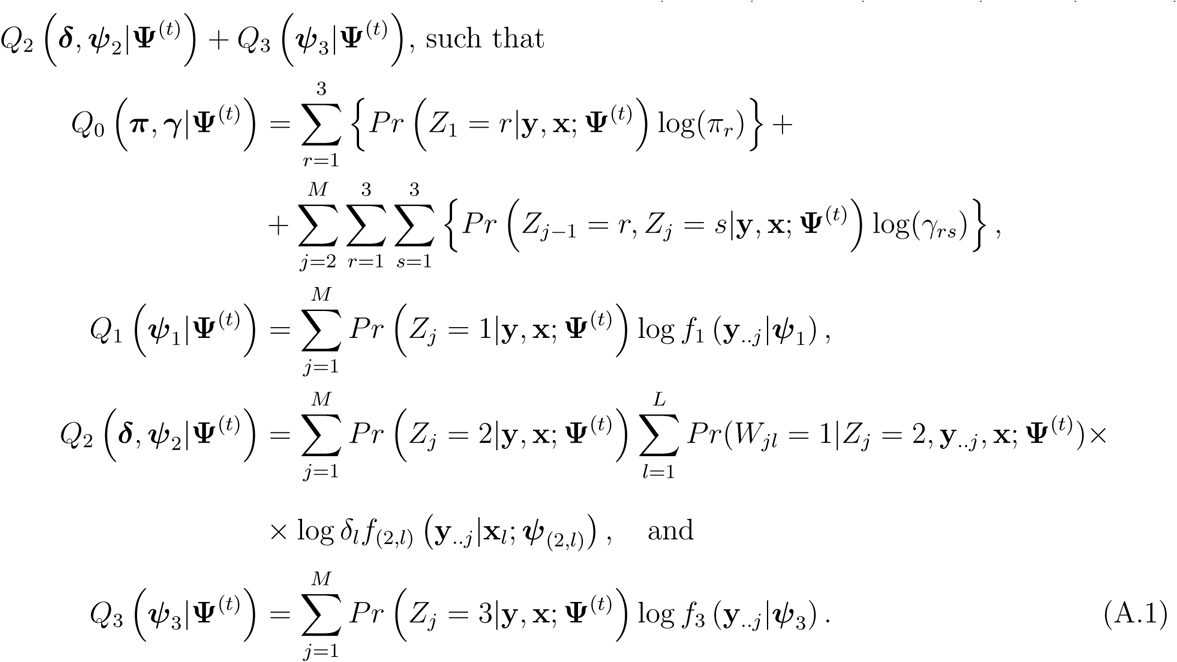

The forward probabilities are defined as 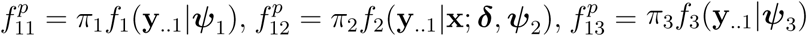, and, for 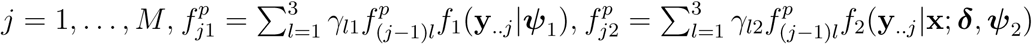 and 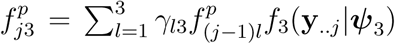. Conversely, the backward probabilities are defined as 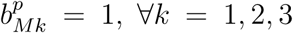 and, for 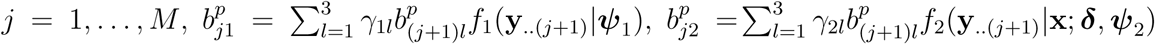 and 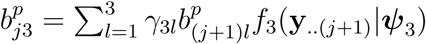. Then, we have

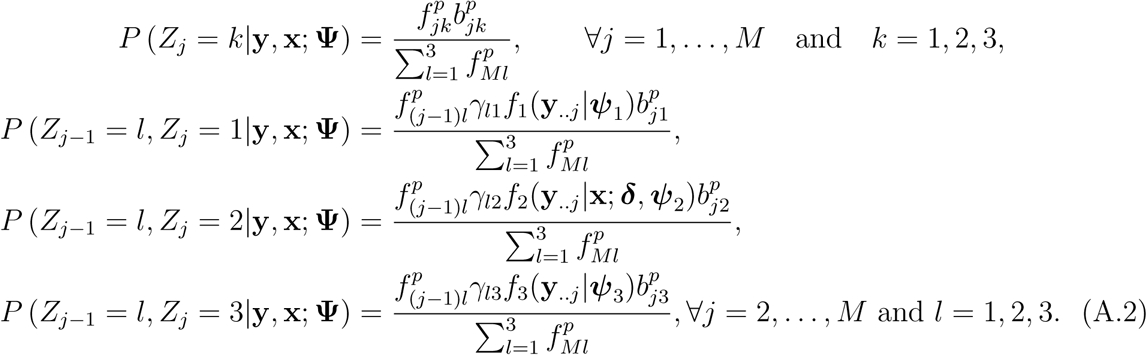

## Notes

### Competing Interest Statement

The authors have declared no competing interest.

