## Supplementary Materials for "Efficient Detection and Classification of Epigenomic Changes Under Multiple Conditions"

May 29, 2020

### Supporting Information A

In this appendix, we present a summary of data accession codes, data pre-processing, and parameter specifications from benchmarked methods.

#### A1 Data accession codes

Web Table 1: GEO sample accession codes of the analyzed data from the ENCODE Consortium

| Cell Line | H3K27me3 | H3K36me3 | EZH2 | H3K4me3 | H3K27ac | CTCF | RNA-seq |
|---|---|---|---|---|---|---|---|
| H1hesc | GSM733748 | GSM733725 | GSM1003524 | GSM733657 | GSM733718 | GSM733672 | GSM758566 |
| HelaS3 | GSM733696 | GSM733711 | GSM1003520 | GSM733682 | GSM733684 | GSM733785 | GSM765402 |
| Hepg2 | GSM733754 | GSM733685 | GSM1003487 | GSM733737 | GSM733743 | GSM733645 | GSM758575 |
| Huvec | GSM733688 | GSM733757 | GSM1003518 | GSM733673 | GSM733691 | GSM733716 | GSM758563 |

#### A2 Data Processing

First, we removed PCR duplicates from the BAM files using SAMTools (Li et al., 2009) and converted the resulting indexed and sorted files to BED format using BEDTools (Quinlan and Hall, 2010), as RSEG only accepts such a format as input. Then, the fragment length of each ChIP-seq experiment was estimated using csaw and its functions *correlateReads* and *maximizeCcf*. Finally, using the estimated fragment length, read counts from all cell lines were tabulated for their ChIP replicates using fixed-step and non overlapping windows of size 250bp, 500bp, 750bp, and 1000bp through the R package *bamsignals* (Mammana and Helmuth, 2016). For all methods using window-based approaches (csaw, ChIPComp, diffReps, RSEG, THOR, and mixNBHMM), we assessed their performance with different window sizes. See Section D2 for a discussion about results with different window sizes.

All the methods considered in the data applications and simulation study outputted a set of differential genomic regions/windows that were used for benchmark purposes. THOR output a list of differential peaks in BED6+4 format (*narrowPeak*) with adjusted p-values. RSEG output a WIG file with genomic windows and their posterior probabilities for differential enrichment. diffReps output an annotated TXT file with differential regions of enrichment and their adjusted p-values. DiffBind output a TXT file with differential regions of enrichment and their respective multiple testing corrected FDR. diffReps output a TXT file with differential regions of enrichment and their p-values. csaw output a TSV file with differential regions of enrichment and their FDR adjusted p-values. For a fair FDR thresholding comparison, we control the total FDR and output the differential regions of enrichment based on the set of posterior probabilities as described in the main text. For a comparison between the Viterbi and the FDR thresholding approach, see Section B4.

The following parametrization was used when calling peaks from the benchmarked methods:

- THOR: *rgt-THOR 'config' –name 'name' -b 'bp' –pvalue 1.0 –output-dir 'output'*,
- RSEG: *rseg-diff -verbose -mode 3 -out 'output' -score 'score' -chrom 'chrom' -bin-size 'bp' -deadzones 'deadzonee' -duplicates 'sample1' 'sample2'*,
- ChIPComp: *ChIPComp(makeCountSet(conf,design,filetype="bam",species="hg19",binsize=bp))*,
- diffReps: *diffReps.pl –gname hg19 –report 'output' –treatment 'sample1' –control 'sample2' –btr 'control1' –bco 'control2' –window 'bp' –pval 1 –nsd 'marktype' –meth 'nb'*,
- DiffBind: *dba.report(dba.analyze(dba.contrast(dba.count(dba(sampleSheet = conf)), categories=DBA_CONDITION,minMembers=2)),th=1)*,

such that bp $= \{250, 500, 750, 100\}$ and $marktype =' broad'$ if H3K27me3, H3K36me3, or EZH2, or $marktype =' sharp'$ otherwise.

For DiffBind under 3 conditions (Figure 1B, main text), the set of differential peaks included all peaks deemed to be differential by DiffBind under an FDR control of 0.05 simultaneously for all three pairwise contrast tests between the cell lines Helas3, Hepg2, and Huvec. In the particular genomic position shown in Figure 1B, no differential peaks were reported by DiffBind.

For csaw, we used the following setup:

```
# List of bam files
bam.files = list.files(path=paste0(tmpdir,'/chip'),pattern='*.bam$',
                       full.names=T)
bam.files

# Design matrix
design <- model.matrix(~factor(cell.type))
colnames(design) <- c("intercept", "cell.type")
design

# Parameters (PCR duplicates already removed and quality score filtered)
param <- readParam(dedup = F)
param

# Estimating the average fragment length (rescaling all to 200bp)
x = lapply(bam.files,correlateReads,param=param,max.dist=250)
multi.frag.lens = list(unlist(lapply(x,maximizeCcf)),200)
multi.frag.lens
```

```
# Counting reads (for a window size of 250bp, for instance)
data <- windowCounts(bam.files,width = 250,ext = multi.frag.lens,
                     param = param,filter = 20)
data

# Filtering data
data.large <- windowCounts(bam.files,width=2500,bin=T,param=param)

bin.ab <- scaledAverage(data.large, scale=median(getWidths(data.large))/
                          median(getWidths(data)))

threshold <- median(bin.ab) + log2(2)

keep.global <- aveLogCPM(asDGEList(data)) >  threshold

sum(keep.global)

# Creating filtered data
filtered.data <- data[keep.global,]

# Testing for DB (assuming composition bias is negligble,
# i.e. cell lines should exhibit a balanced number of DB regions)

y <- DGEList(assay(filtered.data), lib.size = filtered.data$totals)
y$samples$norm.factors <- 1
y$offset <- NULL
y <- estimateDisp(y, design)
```

```
fit <- glmQLFit(y, design, robust = TRUE)
out <- glmQLFTest(fit,contrast = contrast)
tabres <- topTags(out, nrow(out))$table
tabres <- tabres[order(as.integer(rownames(tabres))),]

merged <- mergeWindows(rowRanges(filtered.data), tol=tol,
                       max.width=max.width)
tabneg <- combineTests(merged$id, tabres)
```

For ChIPComp and DiffBind, candidate peaks were called in advance using MACS with the following syntax:

- MACS: *macs2 callpeak -f BAM -g 2.80e+09 -B 'options' -t 'sample' -c 'control' –outdir 'output' -n 'filename'*

such that *options* = {–broad –broad-cutoff 0.1} if H3K27me3, H3K36me3, or EZH2, or *options* = {-q 0.01} otherwise.

#### A3 Software

mixNBHMM was implemented in a R package that is available on the GitHub repository https://github.com/plbaldoni/mixNBHMM.

mixNBHMM is a package with a differential peak caller to detect differential enrichment regions from multiple ChIP-seq experiments with replicates. The main function of the package is *mixNBHMM*(). The package allows the user to specify a set of parameters that control the Expectation-Maximization (EM) algorithm. These parameters include, for instance, the convergence (and termination) criteria of the algorithm and the threshold value for the rejection controlled EM algorithm. These parameters can be defined by the function *controlEM*(). Please refer to the package documentation (e.g. *?mixNBHMM::mixNBHMM*) for additional details and the complete help manual.

#### A4 Code

The necessary code to replicate the results presented in the main article and in the supplementary material can be downloaded from https://github.com/plbaldoni/mixNBHMMPaper.

### Supporting Information B

In this appendix, we present additional methodological results and an overview of the EM algorithm proposed in the main article.

#### B1 Overview of the Proposed Hidden Markov Model

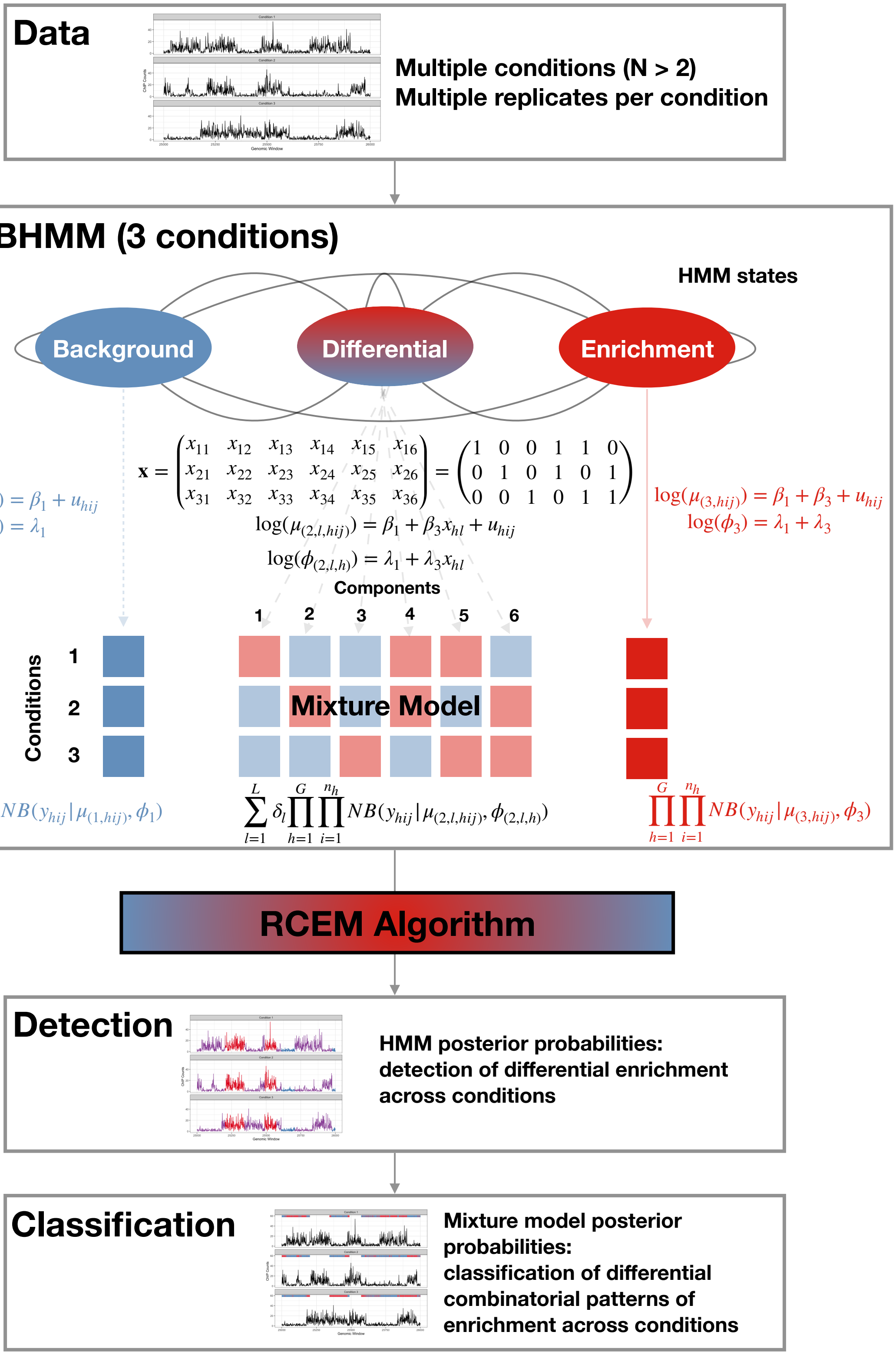


Web Figure 1: Overview of the presented HMM. Given the ChIP-seq data collected under multiple conditions (with or without experimental replicates) , the mixture model embedded in the differential HMM component accounts for all combinatorial patterns of enrichment across multiple groups. Posterior probabilities from the HMM and the mixture model allow the detection and classification, respectively, of epigenomic changes across conditions.

### B2 Adjustments for nuisance effects

#### B22 Normalization for non-linear biases via model offsets

In our analyses, we observed that the magnitude of the local differences in read counts between conditions changed with the average of local read coverage. Here, we accounted for these trended differences to avoid calling spurious differential peaks due to the different magnitude of library sizes across groups. Specifically, we implemented an approach similar to the non-linear normalization method used by csaw as follows (Lun and Smyth, 2015). First, we create a reference sample of read counts formed by the geometric mean of read counts from all replicates and conditions. Then, we fitted a loess curve on the difference between the read counts of each sample and the reference on the average of those two quantities. A similar approach was first implemented by (Lun and Smyth, 2015) and is available in their software. Here, we add a continuity correction of 1 to avoid discarding genomic windows with zero counts. Using the smoothed curve as the model offset, we observed better results than a simple correction via either the total sum of read counts or cell-specific median log ratio. The rationale behind this approach is to create a reference library in which each genomic window is the geometric mean of counts across all conditions and replicates, and then read counts are properly adjusted by accounting for the smoothed differences between each individual library and the reference library. A useful way to evaluate the performance of this normalization method is to compare samples with respect to their adjusted read counts. For example, plotting the ratios between counts and the calculated offsets $y_{hij}/exp(u_{hij})$ for all samples in the study. In Figure 2 we show an example of a genomic region from three analyzed cell lines and their respective MA plot, unadjusted ChIP counts, and offset-adjusted ChIP counts. After accounting for the offset, the read counts from Helas3 are adjusted to its larger library size with respect to the other under sequenced cell lines.

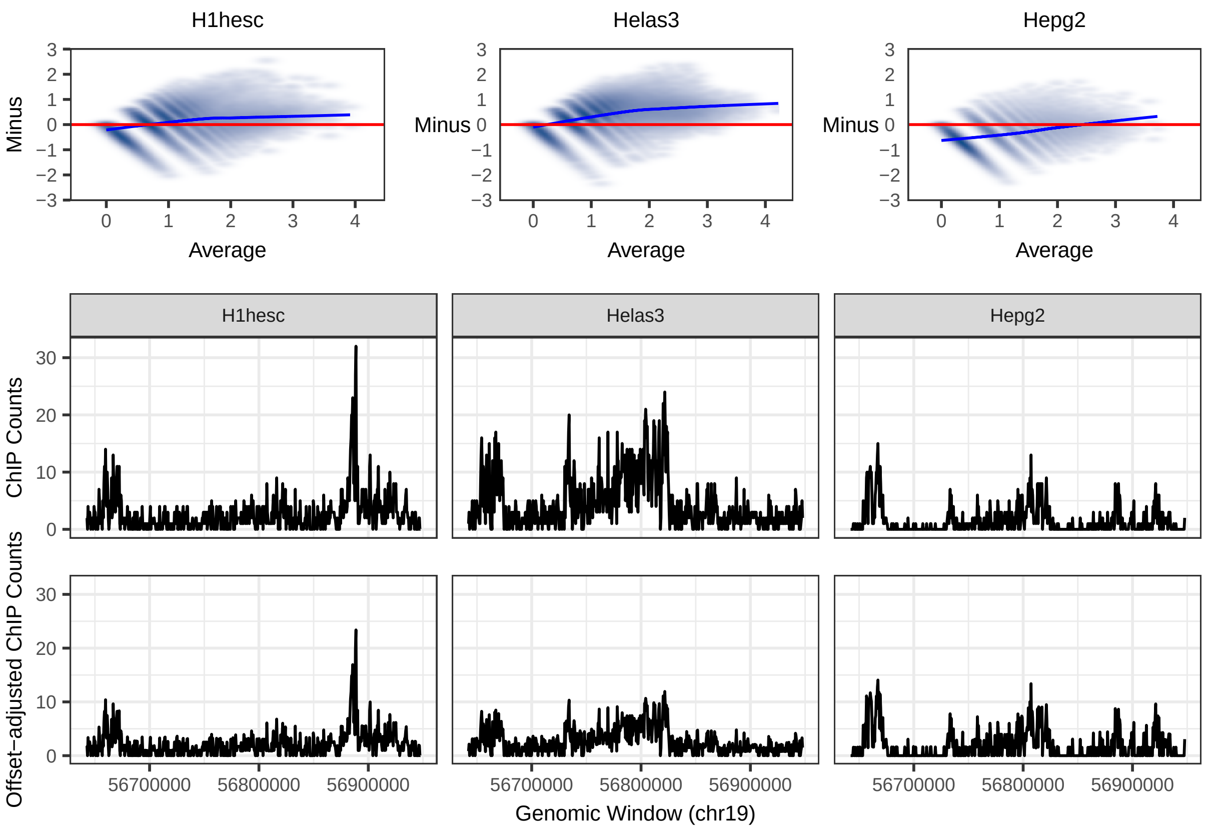


Web Figure 2: MA plot of read counts from three distinct analyzed cell lines (top), unadjusted ChIP read counts (center), and offset-adjusted ChIP read counts (bottom) from a given genomic region on chromosome 19. The blue line in the MA plots shows the offset created via loess smoothing.

#### B22 Input control adjustment in differential peak calling

Our implementation (Web Appendix A4) allows the optional inclusion of continuous covariates in the model with state-specific parametrization. The main purpose of the inclusion of such covariates in the model is the adjustment for input control (or any other continuous variable, such as autoregressive counts) that can be helpful in distinguishing background from enrichment signal. Several methods for differential peak calling allow the inclusion of input control in their computational framework (Stark and Brown, 2011; Shen et al., 2013; Chen et al., 2015). However, Lun and Smyth (2015) point out that "(...) controls are mostly irrelevant when testing for DB between ChIP samples.". To evaluate this claim, we ran an analysis of real data and simulated data while accounting for the input control effect.

To asses whether accounting for input control effect leads to an improvement in performance, we utilized the smoothing technique proposed by Chen et al. (2015) to account for input controls and autoregressive counts. Specifically, we fitted generalized additive models (GAM, instead of loess smoothing) in the data normalization step while accounting for input control (or autoregressive counts) as a covariate. The resulting fitted curve was then used in the analysis as model offsets.

First, we analyzed real data by smoothing the input control effect and autoregressive counts with a two-step approach. Specifically, we first called peaks without the inclusion of extra covariates in the model, and then utilized the called differential peaks from the first step to smooth the covariates for each HMM predicted state. Predicted smoothing curves from the GAM approach were then passed as model offsets in a second step of analysis. As claimed by Lun and Smyth (2015), we observed minor differences in the results that would justify their inclusion in the analysis. Results from the histone modification mark H3K36me3 are presented in Figure 3.

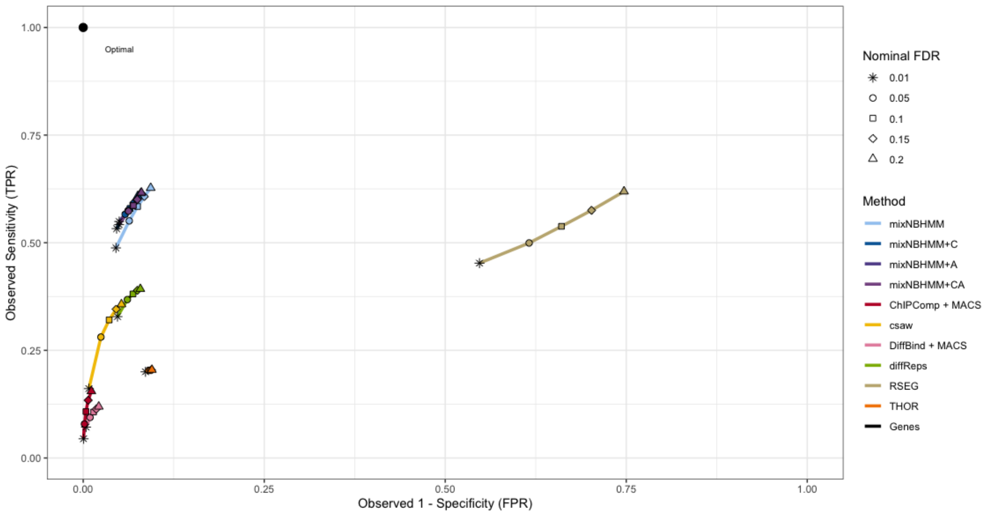


Web Figure 3: ROC curves for H3K36me3 utilizing no input controls (mixNBHMM), input control only (mixNBHMM + C), autoregressive counts only (mixNBHMM + A), and smoothing of both input controls and autoregressive counts (mixNBHMM + CA)

Next, we reasoned that our approach of modeling input control effect with state-specific parametrization could not be ideal, since independent controls were available for every sample and there could exist sample-specific effects not captured by our model. We then attempted to verify the utility of including input control into the differential binding analysis by simulating data where ChIP-counts were generated such that their log-mean had a linear relationship with input controls (Figure 4). We then fitted three different models that differed regarding the inclusion of input control: a model without control, a model with control, and a model with controls where the smoothing was calculated separately for each latent HMM state. Again, results did not show significant improvement by including the effect of control in the analysis.

Overall, we did not observe a significant improvement in performance by including input control in differential peak calling. Although several methods do offer the option of including controls in their analysis pipeline, we did not find that their inclusion was justifiable under our modelling assumptions. Our findings are in agreement with Lun and Smyth (2015).

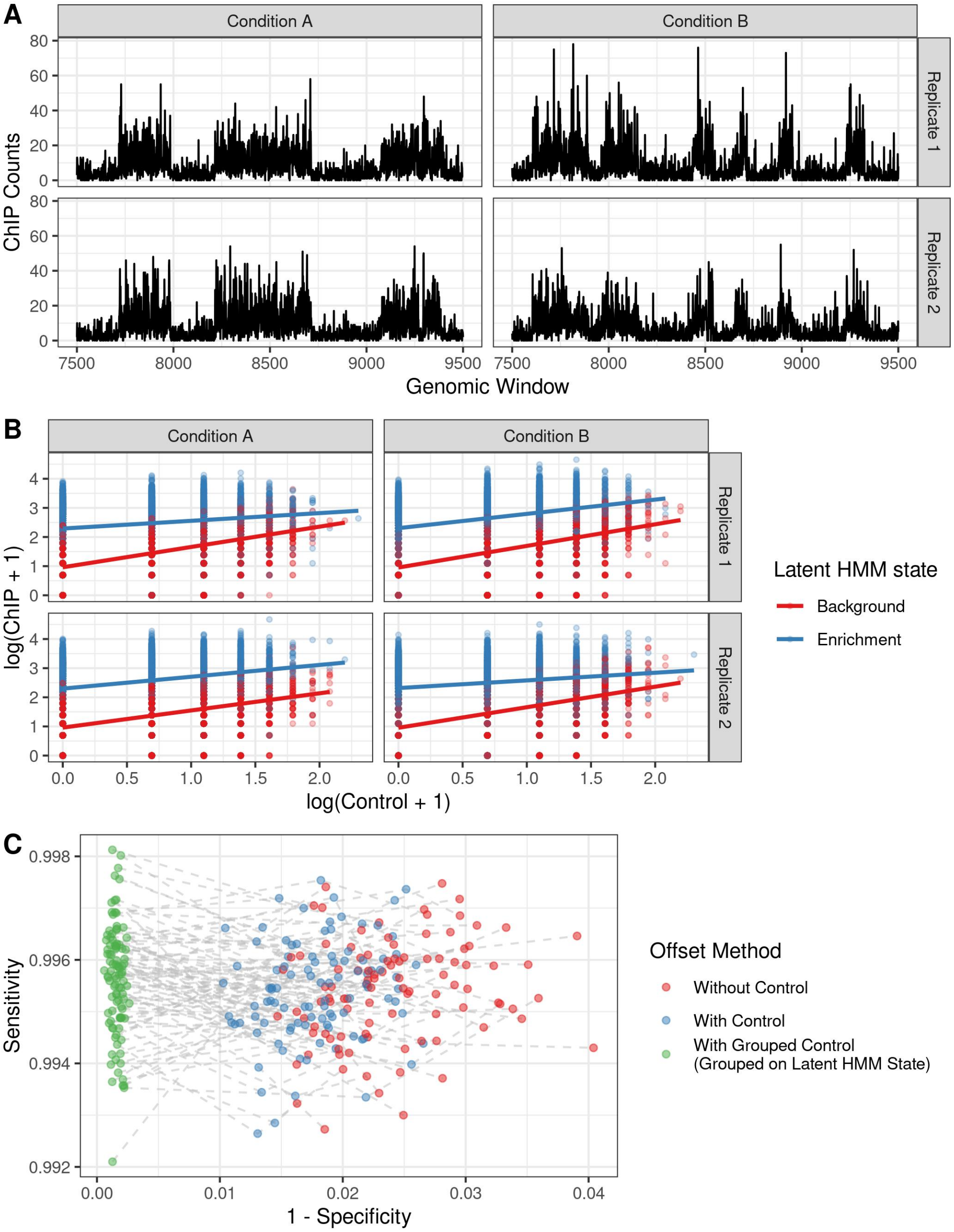


Web Figure 4: Results from simulated data (A) where the log-means of ChIP-seq counts were generated as a linear function of input controls (B). Sensitivity/specificity analyses did not show significant improvement by including the effect of control in the offset scheme.

### B3 Bayesian Information Criterion (BIC) for Hidden Markov Models

The BIC for hidden Markov models has been discussed by Zucchini et al. (2017). For the presented three-state HMM, one can calculate the BIC as

$$BIC = -2\log\left(\sum_{r=1}^{3} f_{Mr}^{p}\right) + (11+L)\log\left(M\sum_{h=1}^{G} n_h\right), \tag{B.1}$$

where $f_{Mr}^{p}$ is the forward probability pertaining to the $r^{th}$ state calculated at the (last) $M^{th}$ genomic window (as detailed in the Appendix of the main text), $L$ is the number of mixture components, $G$ is the number of conditions, and $n_h$ is the number of replicates pertaining to condition $h$. The number of model parameters to be estimated is $(11+L)$: 6 transition probabilities, 2 initial probabilities, 4 model coefficients pertaining to the emission distributions, and $L-1$ prior probabilities from the mixture model.

As shown in the main text, the proposed HMM is robust to situations where certain combinatorial patters are rare. However, if pruning rare combinatorial patterns is still of interest, such a task can be performed by making use of the BIC. For the analysis of $G$ experimental conditions with a given BIC threshold $\epsilon$, say $\epsilon = 0.01$, and $L = 2^G - 2$ mixture components, one can prune rare combinatorial patterns by the following algorithm.

1. Fit the three-state HMM with $L$ mixture components (model 0) and compute the model BIC, $\text{BIC}_0$, as in Equation B.1.

2. Fit a reduced three-state HMM with $L-1$ mixture components (model 1) by excluding the component associated with the rarest combinatorial pattern of enrichment. Compute its BIC, $\text{BIC}_1$.

3. Calculate $\Delta\text{BIC} = (\text{BIC}_1 - \text{BIC}_0)/\text{BIC}_0$. If $|\Delta\text{BIC}| \leq \epsilon$, set $L \leftarrow L-1$ and return to 1. If $|\Delta\text{BIC}| > \epsilon$, stop and set the model 0 as the final model.

In scenarios where the number of mixture components is smaller than $2^G - 2$, the implemented method initializes the EM algorithm by clustering genomic windows with respect to the posterior probabilities of enrichment obtained from a initial run of a two-state HMM to classify genomic windows into background and enrichment windows. Such an initialization improves the overall computation time by reducing the time to convergence of the presented EM algorithm.

We applied the above approach in real data where the goal was to reduce the number of rare mixture components. Specifically, we reanalyzed the data presented in Section 5.3 of the main text by refitting the presented model with reduced number of combinatorial patterns. Figure 5 presents the BIC from various models regarding the number of mixture components on data from epigenomic marks H3K37me3, H3K36me3, and EZH2. As shown, models with more than 2 mixture components exhibited values of BIC quite close to each other. Conversely, the model with a single differential component had an excessively large BIC. These results suggest that, according to the BIC, the parsimonious model with only 2 mixture components would be the one chosen. As detailed in the main text, the analyzed data sets are characterized by only 2 combinatorial patterns of enrichment, which are associated with the enrichment of H3K36me3 alone, and the enrichment of H3K27me3 and EZH2 in consensus. Hence, choosing the model with 2 components via BIC agrees with the biological roles of the analyzed marks.

We also compare the results of our method with ChromHMM, an algorithm developed for chromatin segmentation. In Figure 6, panels A-D, we present results of ChromHMM with 3 (ideal), 4, 5, and 6 states. By using the BIC for model selection, one could easily choose the number of biologically relevant mixture components to be included in the model, a task that may not be as straightforward in methods such as ChromHMM (see Supplementary Figure 4 in Ernst and Kellis (2012)). Our method offers the benefit of simultaneously detecting differential peaks and classifying the combinatorial pattern of enrichment through mixture model posterior probabilities even in the context of genomic segmentation. Despite the

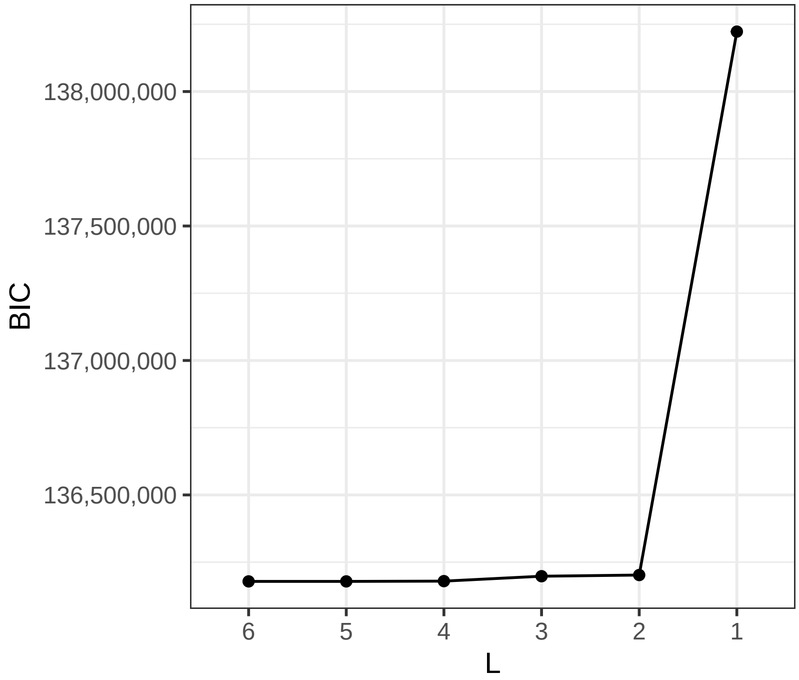


Web Figure 5: BIC from various models regarding their number of mixture components on epigenomic marks H3K36me3, H3K27me3, and EHZ2 (Section 5.3 of the main text)

choice of the number of mixture components via BIC, mixNBHMM appeared to be robust in scenarios with rare combinatorial patters. This is so because we utilize a constrained parametrization in which only 4 GLM-specific parameters need to be estimated regardless of the number of mixture components, and in the genome-wide analysis of ChIP-seq data one often has enough data to estimate such quantities.

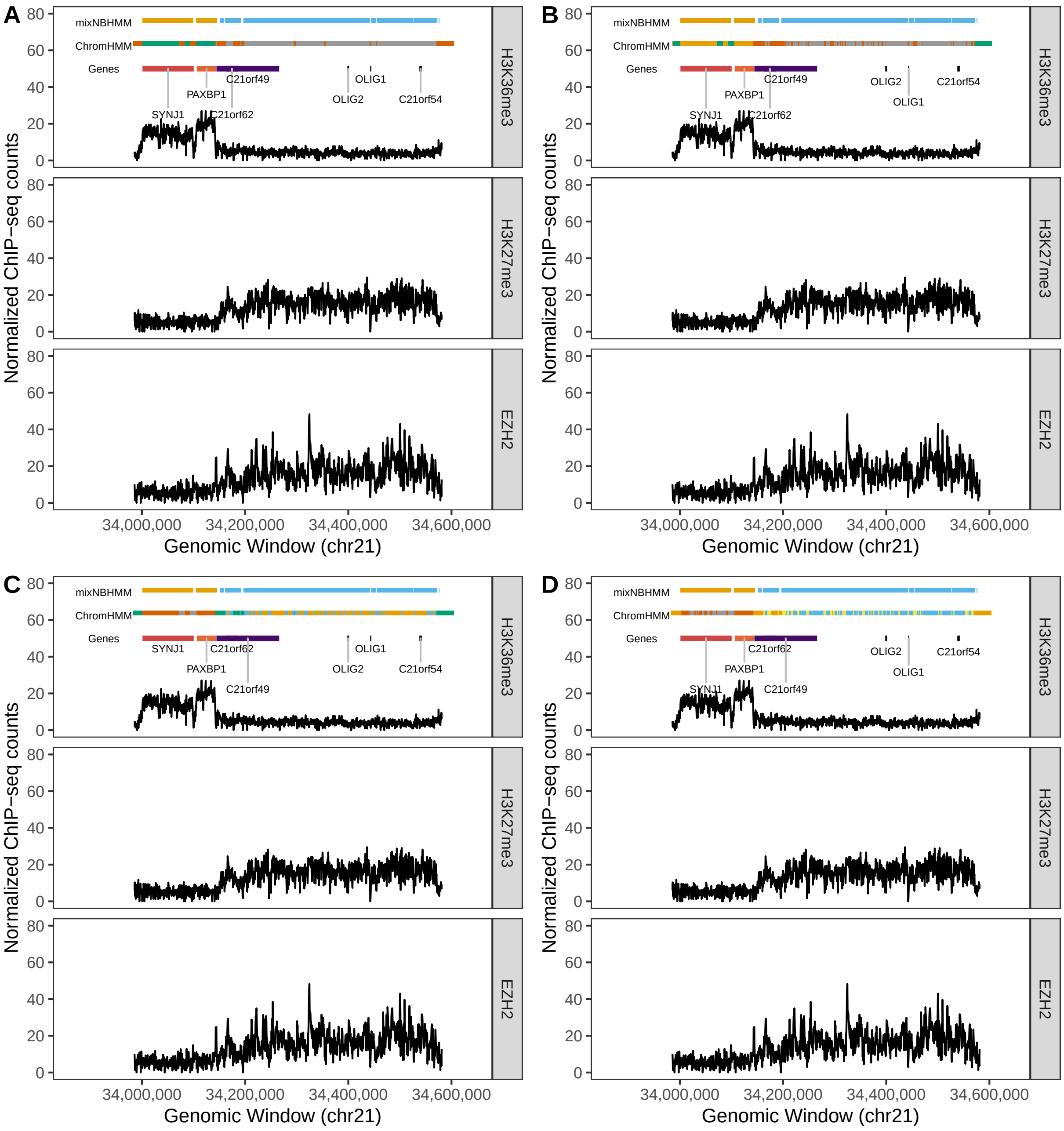


Web Figure 6: Comparative analysis of mixNBHMM and ChromHMM with 3 (ideal), 4, 5, and 6 states (panels A-D).

### B4 Study on FDR Control and the Viterbi Algorithm

In this section, we present results comparing the FDR controlling approach with the Viterbi algorithm. We evaluated the FDR approach using cutoffs 0.01, 0.05, 0.10, 0.15, and 0.20. We compared the results between the two approaches using window sizes of 250bp, 500bp, 750bp, and 1000bp. Overall, we observed that the Viterbi sequence of states led to similar results than the sequences based on FDR control cutoffs across all choices of window size. Specifically, we observed that the sensitivity and specificity of the sequence of Viterbi states were close to those from FDR control, in particular for FDR control 0.10. These results are shown in Figures 7, 8, 9, and 10. These facts are also reflected by the length and number of called peaks. In Figures 11 and 12 we show examples of peak calls of H3K26me3 and H3K27me3, respectively, from all FDR control cutoffs and the Viterbi sequence of states. Overall, we observe minor differences regarding the size of peak calls of the Viterbi and FDR control sequences across different choices of window sizes. These differences were mainly present in the data for H3K27me3, which is known to be a histone mark that expands through broader domains than H3K36me3. Finally, it is worth noting that the Viterbi algorithm gives us a way to call peaks that does not depend on the choice of the FDR cutoff.

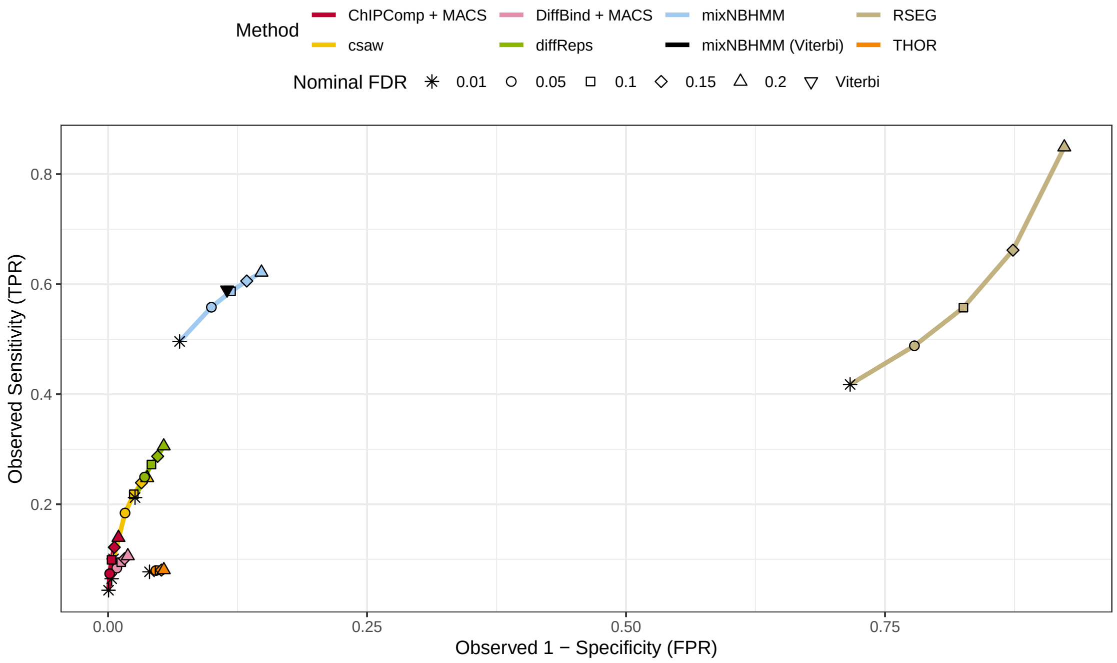


Web Figure 7: FDR-based results from broad marks (250bp) and Viterbi-based result from mixNBHMM.

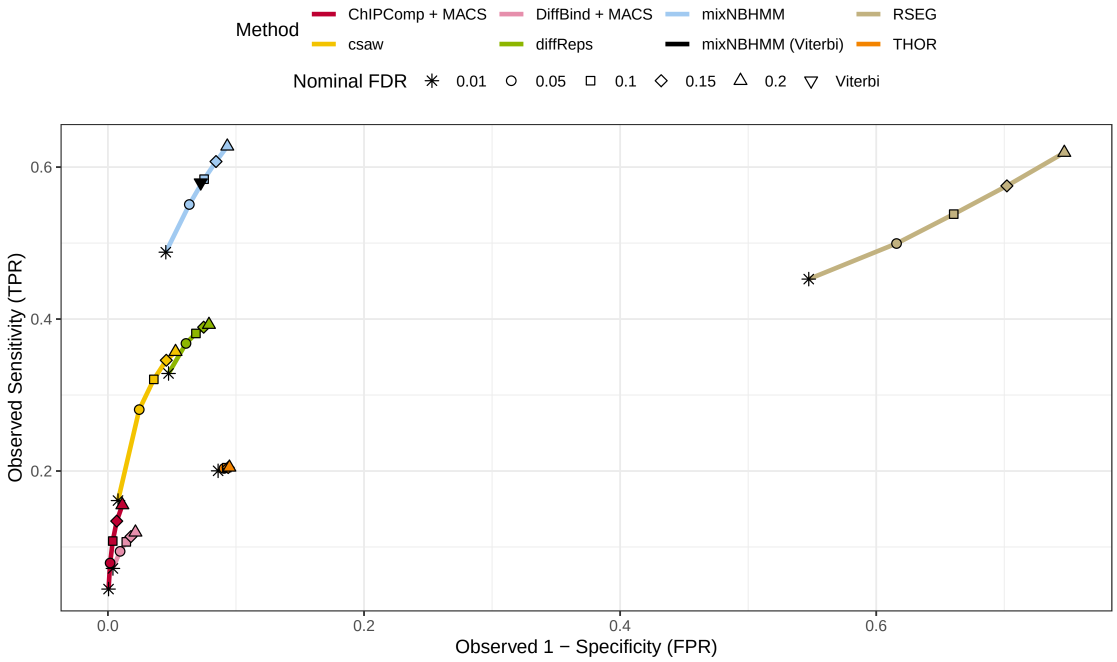


Web Figure 8: FDR-based results from broad marks (500bp) and Viterbi-based result from mixNBHMM.

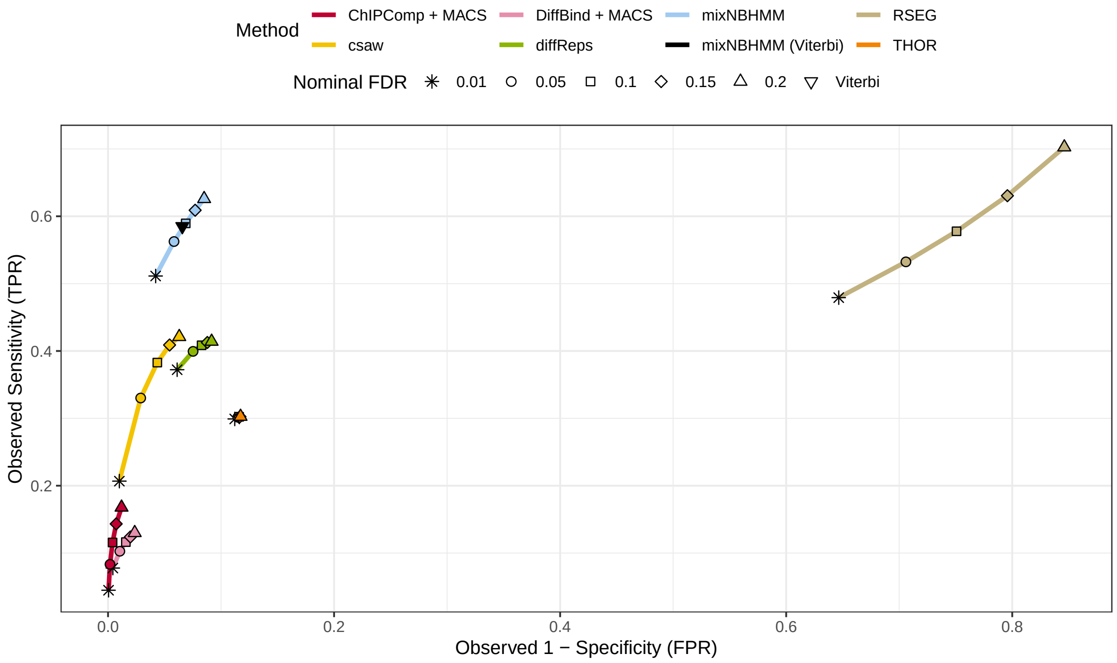


Web Figure 9: FDR-based results from broad marks (750bp) and Viterbi-based result from mixNBHMM.

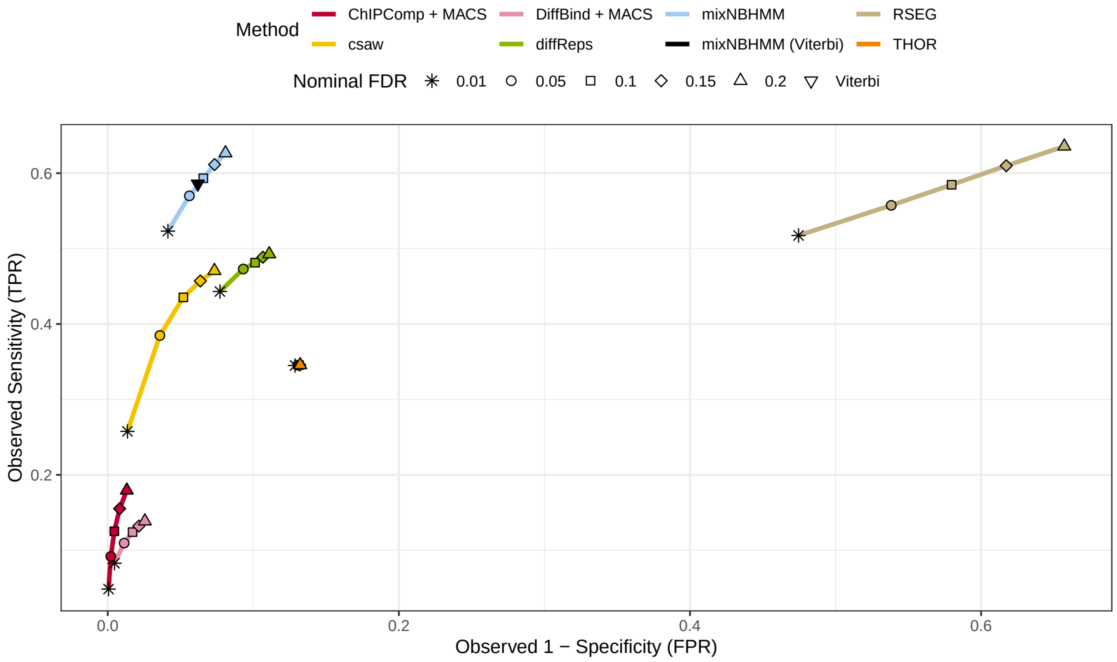


Web Figure 10: FDR-based results from broad marks (1000bp) and Viterbi-based result from mixNBHMM.

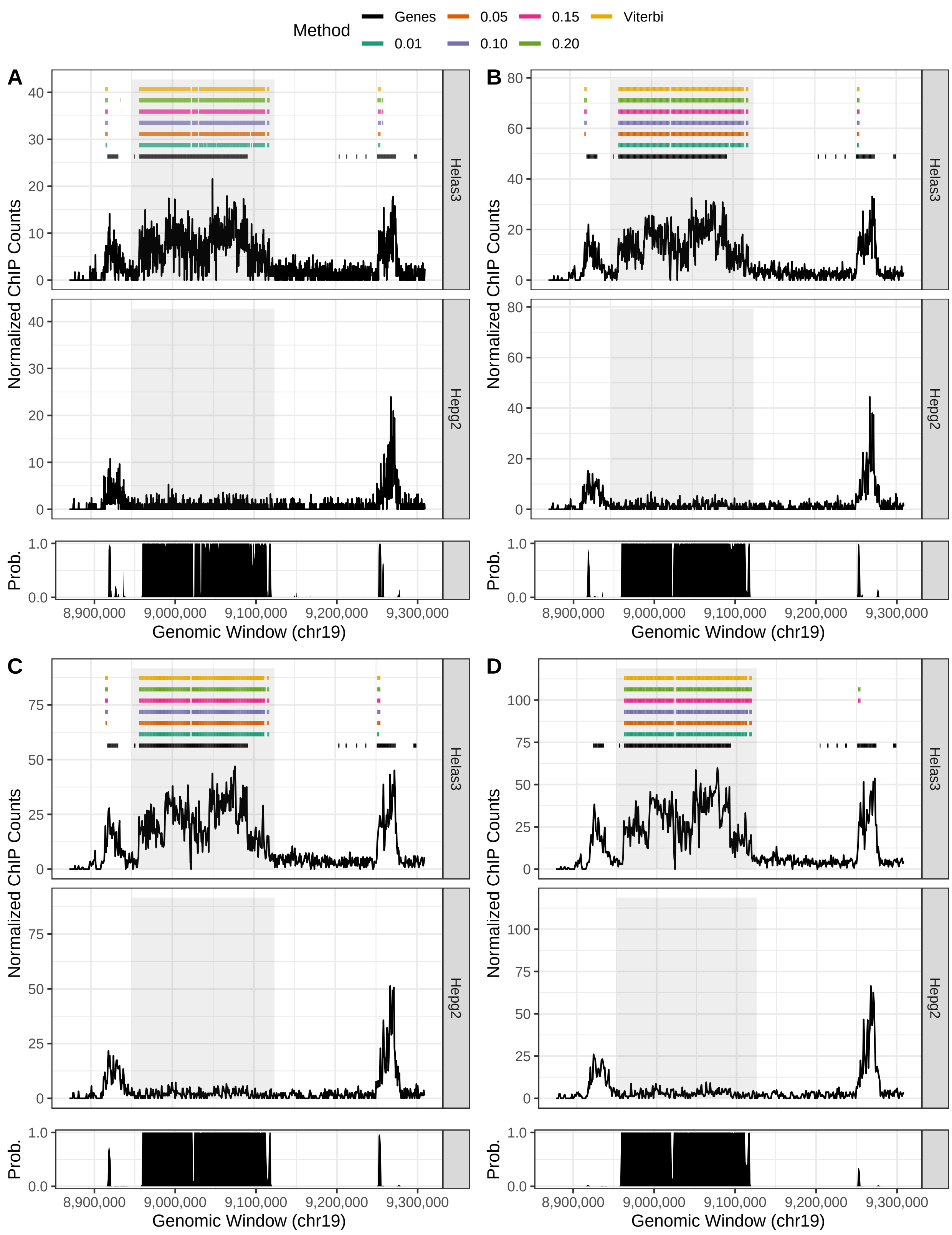


Web Figure 11: FDR- and Viterbi-based peak calls from H3K36me3 with 250bp (A), 500bp, (B), 750bp (C), and 1000bp (D).

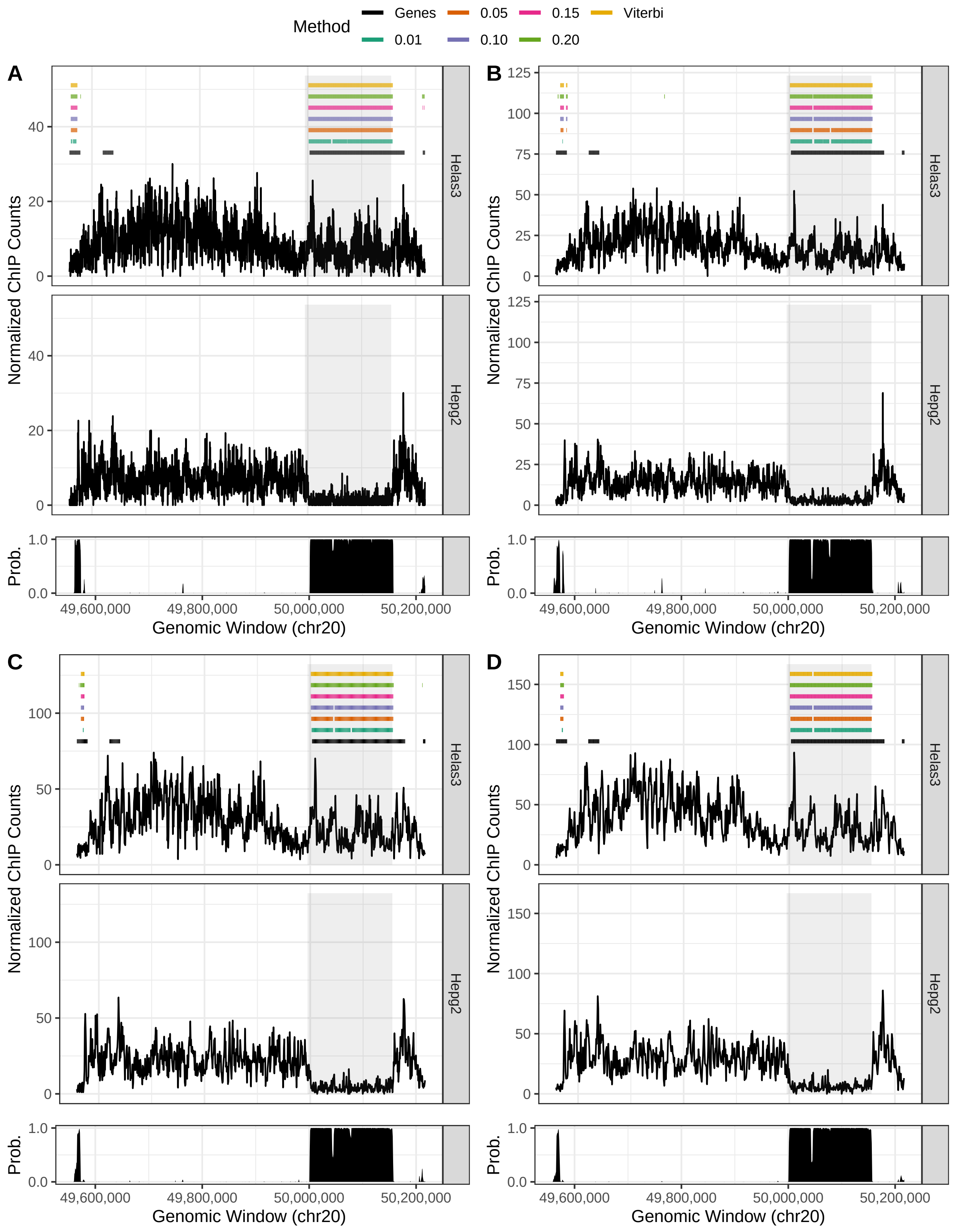


Web Figure 12: FDR- and Viterbi-based peak calls from H3K27me3 with 250bp (A), 500bp, (B), 750bp (C), and 1000bp (D).

### B5 HMM Emission Distributions

For the consensus background ($r = 1$) and consensus enrichment ($r = 3$) states, the emission distribution function is

$$\begin{aligned} f_r(\mathbf{y}_{..j}|\boldsymbol{\psi}_r) &= \prod_{h=1}^{G}\prod_{i=1}^{n_h} f_r(y_{hij}|\boldsymbol{\psi}_r), \quad r \in \{1,3\} \quad \text{and} \quad y_{hij} \in \{0,1,2,\ldots\}, \\ &= \prod_{h=1}^{G}\prod_{i=1}^{n_h} \Pr(Y_{hij} = y_{hij}|Z_j = r; \boldsymbol{\psi}_r), \\ &= \prod_{h=1}^{G}\prod_{i=1}^{n_h} \frac{\Gamma(y_{hij}+\phi_r)}{y_{hij}!\Gamma(\phi_r)} \left(\frac{\phi_r}{\mu_{(r,hij)}+\phi_r}\right)^{\phi_r} \left(\frac{\mu_{(r,hij)}}{\mu_{(r,hij)}+\phi_r}\right)^{y_{hij}}. \end{aligned} \tag{B.2}$$

For the differential state ($r = 2$), the emission distribution is

$$\begin{aligned} f_2(\mathbf{y}_{..j}|\mathbf{x};\boldsymbol{\delta},\boldsymbol{\psi}_2) &= \sum_{l=1}^{L} \delta_l f_{(2,l)}\left(\mathbf{y}_{..j}|\mathbf{x}_l;\boldsymbol{\psi}_{(2,l)}\right), \quad y_{hij} \in \{0,1,2,\ldots\}, \\ &= \sum_{l=1}^{L} \delta_l \prod_{h=1}^{G}\prod_{i=1}^{n_h} Pr\left(Y_{hij} = y_{hij}|Z_j = 2, \mathbf{x}_l; \boldsymbol{\psi}_{(2,l)}\right), \\ &= \sum_{l=1}^{L} \delta_l \prod_{h=1}^{G}\prod_{i=1}^{n_h} \frac{\Gamma\left(y_{hij}+\phi_{(2,l,h)}\right)}{y_{hij}!\Gamma\left(\phi_{(2,l,h)}\right)} \left(\frac{\phi_{(2,l,h)}}{\mu_{(2,l,hij)}+\phi_{(2,l,h)}}\right)^{\phi_{(2,l,h)}} \times \\ &\times \left(\frac{\mu_{(2,l,hij)}}{\mu_{(2,l,hij)}+\phi_{(2,l,h)}}\right)^{y_{hij}}. \end{aligned} \tag{B.3}$$

Apart from the offset $u_{hij}$, we will assume that replicates from the same (different) condition share common (distinct) mean and dispersion parameters under every mixing probability distribution $f_{(2,l)}$. To define all possible combinations of background and enrichment across $G$ conditions, we consider the following sets of singletons $A_1$, pairs $A_2$, ..., and $(G-1)$-tuples

$A_{G-1}$ such that

$$
\begin{aligned}
A_1 &= \left\{a^{(1)} \mid a^{(1)} \in \mathbb{G}_+ \text{ and } a^{(1)} \leq G\right\}, \\
A_2 &= \left\{(a_1^{(2)}, a_2^{(2)}) \mid (a_1^{(2)}, a_2^{(2)}) \in \mathbb{G}_+^2 \text{ and } a_1^{(2)} < a_2^{(2)} \leq G\right\}, \\
&\vdots \\
A_{G-1} &= \left\{(a_g^{(G-1)})_{g=1}^{G-1} \mid (a_g^{(G-1)})_{g=1}^{G-1} \in \mathbb{G}_+^{G-1} \text{ and } a_1^{(G-1)} < \ldots < a_{G-1}^{(G-1)} \leq G\right\}.
\end{aligned}
$$

The union of all sets $A = \cup_{k=1}^{G-1} A_k$ contains an exhaustive list of $L = 2^G - 2$ elements that determines the differential pattern across $G$ conditions such that each element of $A$ indicates which of the $G$ conditions are enriched. For instance, if $G = 3$, $A_1 = \{1, 2, 3\}$ and $A_2 = \{(1, 2), (1, 3), (2, 3)\}$ define the six possible combinations of enrichment and background across three conditions. Then, we define a bijective mapping $A \to S_1, \ldots, S_L$ and let $x_{hl} = I(h \in S_l)$ indicate whether the read count of genomic window $j$ from replicate $i$ of condition $h$ is enriched in the mixture component $l$. We model the log-mean $\mu_{(2,l,hij)}$ and log-dispersion $\phi_{(2,l,h)}$ of mixture $l$ from the emission distribution (B.3) as

$$
\begin{aligned}
\log(\mu_{(2,l,hij)}) &= \beta_1 + \beta_3 x_{hl} + u_{hij}, \quad \text{and} \\
\log(\phi_{(2,l,h)}) &= \lambda_1 + \lambda_3 x_{hl}.
\end{aligned}
$$

According to this parametrization, $\beta_1$ and $\lambda_1$ are the baseline log-mean and log-dispersion parameters of the read count distribution from replicates of conditions that are not enriched under the mixing distribution $l$. Conversely, $\beta_1 + \beta_3$ and $\lambda_1 + \lambda_3$ are the baseline log-mean and log-dispersion parameters of the read count distribution from replicates of conditions enriched under the mixing distribution $l$. This choice of parametrization ensures that windows exhibiting differential enrichment across conditions share means and dispersions that are common between the remaining non differential HMM states.

A pseudo code of the presented EM algorithm is below.

1. Initialize $\boldsymbol{\pi}^{(0)}, \boldsymbol{\gamma}^{(0)}, \boldsymbol{\delta}^{(0)}, \beta_1^{(0)}, \beta_3^{(0)}, \lambda_1^{(0)}, \lambda_3^{(0)}$, such that $\sum_{r=1}^{3} \pi_r^{(0)} = 1$ and $\sum_{s=1}^{3} \gamma_{rs} = 1$.

2. E step ($t \geq 1$),

   (a) Calculate $Pr\left(Z_j = r|\mathbf{y}, \mathbf{x}; \boldsymbol{\Psi}^{(t-1)}\right)$ and $Pr\left(Z_{j-1} = r, Z_j = s|\mathbf{y}, \mathbf{x}; \boldsymbol{\Psi}^{(t-1)}\right)$ for all $r$ and $s$ in $\{1, 2, 3\}$ and $j = 1, \ldots, M$ via Forward-Backward algorithm as detailed in Appendix B of the main article

   (b) Calculate $Pr(W_{jl} = 1|Z_j = 2, \mathbf{y}_{..j}, \mathbf{x}; \boldsymbol{\Psi}^{(t-1)})$ for all $l$ $\{1, \ldots, L\}$ and $j = 1, \ldots, M$ as $f_{(2,l)}(\mathbf{y}_{..j}|\mathbf{x}_l; \boldsymbol{\psi}_{(2,l)}^{(t-1)})\delta_l^{(t-1)} / \sum_{k=1}^{L} f_{(2,k)}(\mathbf{y}_{..j}|\mathbf{x}_k; \boldsymbol{\psi}_{(2,k)}^{(t-1)})\delta_k^{(t-1)}$

3. M step ($t \geq 1$),

   (a) Maximize Equation (5) (main text) with respect to the initial and transition probabilities to obtain for all $r$ and $s$ in $\{1, 2, 3\}$

$$\pi_r^{(t)} = Pr\left(Z_1 = r|\mathbf{y}, \mathbf{x}; \boldsymbol{\Psi}^{(t-1)}\right)$$

$$\gamma_{rs}^{(t)} = \sum_{j=2}^{M} Pr(Z_{j-1} = r, Z_j = s|\mathbf{y}, \mathbf{x}; \boldsymbol{\Psi}^{(t-1)}) / \sum_{j=2}^{M} Pr(Z_{j-1} = r|\mathbf{y}, \mathbf{x}; \boldsymbol{\Psi}^{(t-1)})$$

   (b) Maximize Equation (5) with respect to $\boldsymbol{\delta}$ to obtain $\boldsymbol{\delta}^{(t)}$ such that $\sum_{l=1}^{L} \delta_l^{(t)} = 1$.

   (c) Conditionally upon $\boldsymbol{\delta}^{(t)}$, maximize Equation (5) with respect to $\beta_1, \beta_3, \lambda_1, \lambda_3$ to obtain $\beta_1^{(t)}, \beta_3^{(t)}, \lambda_1^{(t)}, \lambda_3^{(t)}$,

   (d) Iterate between (b) and (c) until convergence.

4. Iterate between 2. and 3. until convergence.

### Supporting Information C

In this appendix, we present an overview of the simulation scenarios and additional results.

#### C1 Read Count Simulation

In this section, we present additional results from the read count based simulation.

For each scenario, we simulated a hundred data sets as follows. First, we generated a sequence of hidden states with length $M$ from a first-order Markov chain with $2^G$ states representing every combination of background and enrichment across $G$ conditions. Here, we let states 1 and $2^G$ represent consensus background and consensus enrichment, respectively. Transition probabilities were chosen such that long sequences of enrichment regions (average length of 50 genomic windows) and equal proportions of differential states were generated, while the majority of the simulated states pertained to consensus background. Secondly, read counts were simulated for $n$ replicates from $G$ conditions following NB distributions with mean and dispersion parameters that led to a given SNR and respected the differential pattern of enrichment determined by the hidden state. For each scenario, this setup led to a simulated matrix of counts of dimension $M \times (n \times G)$. The aim of this study was to assess whether the presented model was able to aggregate all $2^G - 2$ simulated differential hidden states into its differential HMM state while maintaining a precise parameter estimation scheme. The model parameters used in this simulated study were estimated from ENCODE data pertaining to H3K36me3 and H3K27me3 histone modification marks and a shown in

Web Table 2: Read count simulation. True values and average relative bias of parameter estimates (and $2.5^{th}$, $97.5^{th}$ percentiles) across a hundred simulated data sets are shown for H3K27me3 with $10^6$ genomic windows. Scenarios with observed (obs.) SNR and 70% of observed SNR are shown.

| | | | | One Replicate | | Two Replicates | | Four Replicates | |
|---|---|---|---|---|---|---|---|---|---|
| SNR | Conditions | Parameter | True | R. Bias | ($P_{2.5}$,$P_{97.5}$) | R. Bias | ($P_{2.5}$,$P_{97.5}$) | R. Bias | ($P_{2.5}$,$P_{97.5}$) |
| 70% of Observed SNR | Two | $\beta_1$ | 1.116 | 0.000 | (-0.001, 0.001) | 0.000 | (-0.001, 0.001) | 0.000 | (-0.001, 0.001) |
| | | $\beta_3$ | 0.808 | 0.000 | (-0.003, 0.003) | 0.000 | (-0.002, 0.002) | 0.000 | (-0.002, 0.001) |
| | | $\lambda_1$ | 1.281 | 0.000 | (-0.005, 0.005) | 0.000 | (-0.004, 0.002) | 0.000 | (-0.002, 0.002) |
| | | $\lambda_3$ | -0.232 | -0.005 | (-0.034, 0.029) | -0.002 | (-0.030, 0.021) | -0.001 | (-0.016, 0.013) |
| | Three | $\beta_1$ | 1.116 | -0.002 | (-0.003, -0.000) | -0.003 | (-0.004, -0.002) | -0.001 | (-0.001, 0.000) |
| | | $\beta_3$ | 0.808 | -0.122 | (-0.129, -0.114) | -0.012 | (-0.015, -0.009) | -0.001 | (-0.002, 0.000) |
| | | $\lambda_1$ | 1.281 | 0.017 | (0.011, 0.022) | 0.004 | (0.001, 0.007) | 0.000 | (-0.002, 0.002) |
| | | $\lambda_3$ | -0.232 | 0.857 | (0.805, 0.913) | 0.113 | (0.089, 0.142) | 0.010 | (-0.005, 0.023) |
| | Four | $\beta_1$ | 1.116 | 0.000 | (-0.002, 0.001) | -0.012 | (-0.013, -0.011) | -0.011 | (-0.012, -0.009) |
| | | $\beta_3$ | 0.808 | -0.135 | (-0.141, -0.128) | -0.088 | (-0.091, -0.085) | -0.032 | (-0.054, -0.023) |
| | | $\lambda_1$ | 1.281 | 0.018 | (0.012, 0.024) | 0.028 | (0.025, 0.031) | 0.017 | (0.014, 0.023) |
| | | $\lambda_3$ | -0.232 | 0.924 | (0.869, 0.977) | 0.770 | (0.744, 0.798) | 0.367 | (0.276, 0.542) |
| Observed SNR | Two | $\beta_1$ | 1.116 | 0.000 | (-0.001, 0.001) | 0.000 | (-0.001, 0.001) | 0.000 | (-0.001, 0.001) |
| | | $\beta_3$ | 1.165 | 0.000 | (-0.002, 0.001) | 0.000 | (-0.001, 0.001) | 0.000 | (-0.001, 0.001) |
| | | $\lambda_1$ | 1.281 | 0.000 | (-0.004, 0.004) | 0.000 | (-0.003, 0.003) | 0.000 | (-0.002, 0.002) |
| | | $\lambda_3$ | 0.124 | 0.004 | (-0.057, 0.066) | -0.004 | (-0.047, 0.038) | -0.003 | (-0.035, 0.034) |
| | Three | $\beta_1$ | 1.116 | -0.003 | (-0.005, -0.002) | 0.000 | (-0.001, 0.000) | 0.000 | (-0.001, 0.001) |
| | | $\beta_3$ | 1.165 | -0.007 | (-0.010, -0.005) | 0.000 | (-0.001, 0.001) | 0.000 | (-0.001, 0.001) |
| | | $\lambda_1$ | 1.281 | -0.001 | (-0.006, 0.003) | 0.001 | (-0.002, 0.004) | 0.000 | (-0.002, 0.002) |
| | | $\lambda_3$ | 0.124 | -0.317 | (-0.404, -0.223) | -0.023 | (-0.057, 0.012) | -0.003 | (-0.025, 0.024) |
| | Four | $\beta_1$ | 1.116 | -0.014 | (-0.015, -0.012) | -0.007 | (-0.008, -0.006) | 0.000 | (-0.001, 0.000) |
| | | $\beta_3$ | 1.165 | -0.111 | (-0.114, -0.109) | -0.003 | (-0.004, -0.002) | 0.000 | (-0.001, 0.001) |
| | | $\lambda_1$ | 1.281 | -0.012 | (-0.016, -0.007) | 0.007 | (0.004, 0.010) | 0.000 | (-0.001, 0.002) |
| | | $\lambda_3$ | 0.124 | -3.520 | (-3.589, -3.439) | -0.360 | (-0.446, -0.282) | -0.005 | (-0.025, 0.017) |

the tables presented in this section.

Overall, while results were similar regarding different number of genome sizes and histone modification marks, the number of replicates and SNR level played major roles in the performance of the model. In scenarios with either low SNR or high number of conditions, more replicates were needed to achieve better model performance. However, under the observed SNR, two replicates were sufficient to achieve a small relative bias regarding the estimation of most of the model parameters throughout all the evaluated scenarios. This is in agreement with the ENCODE Consortium guidelines, which recommends two technical or biological replicates to achieve best performance in ChIP-seq experiments.

Next, we assessed the sensitivity of our method to detect simulated differential regions of enrichment. First, differential regions were defined from the HMM posterior probabili-

Web Table 3: Read count simulation. True values and average relative bias of parameter estimates (and $2.5^{th}$, $97.5^{th}$ percentiles) across a hundred simulated data sets are shown for H3K36me3 with $10^6$ genomic windows. Scenarios with observed (obs.) SNR and 70% of observed SNR are shown.

| | | | | One Replicate | | Two Replicates | | Four Replicates | |
|---|---|---|---|---|---|---|---|---|---|
| SNR | Conditions | Parameter | True | R. Bias | ($P_{2.5}$,$P_{97.5}$) | R. Bias | ($P_{2.5}$,$P_{97.5}$) | R. Bias | ($P_{2.5}$,$P_{97.5}$) |
| 70% of Observed SNR | Two | $\beta_1$ | 0.756 | 0.000 | (-0.002, 0.002) | 0.000 | (-0.001, 0.001) | 0.000 | (-0.001, 0.001) |
| | | $\beta_3$ | 1.304 | 0.000 | (-0.002, 0.002) | 0.000 | (-0.001, 0.002) | 0.000 | (-0.001, 0.001) |
| | | $\lambda_1$ | 1.173 | 0.000 | (-0.006, 0.006) | 0.000 | (-0.004, 0.003) | -0.001 | (-0.003, 0.002) |
| | | $\lambda_3$ | -0.467 | 0.000 | (-0.015, 0.020) | 0.000 | (-0.012, 0.012) | -0.001 | (-0.010, 0.010) |
| | Three | $\beta_1$ | 0.756 | 0.003 | (0.001, 0.005) | -0.002 | (-0.003, -0.000) | 0.000 | (-0.001, 0.001) |
| | | $\beta_3$ | 1.304 | -0.115 | (-0.122, -0.106) | -0.001 | (-0.002, -0.000) | 0.000 | (-0.001, 0.000) |
| | | $\lambda_1$ | 1.173 | 0.028 | (0.020, 0.036) | 0.002 | (-0.001, 0.006) | 0.000 | (-0.003, 0.002) |
| | | $\lambda_3$ | -0.467 | 0.693 | (0.651, 0.730) | 0.017 | (0.006, 0.030) | 0.000 | (-0.007, 0.008) |
| | Four | $\beta_1$ | 0.756 | -0.009 | (-0.012, -0.007) | -0.020 | (-0.021, -0.018) | -0.001 | (-0.002, 0.000) |
| | | $\beta_3$ | 1.304 | -0.107 | (-0.109, -0.103) | -0.073 | (-0.075, -0.070) | 0.000 | (-0.001, 0.001) |
| | | $\lambda_1$ | 1.173 | 0.052 | (0.045, 0.058) | 0.046 | (0.042, 0.050) | 0.001 | (-0.002, 0.003) |
| | | $\lambda_3$ | -0.467 | 0.736 | (0.715, 0.757) | 0.573 | (0.558, 0.589) | 0.004 | (-0.004, 0.011) |
| Observed SNR | Two | $\beta_1$ | 0.756 | 0.000 | (-0.002, 0.002) | 0.000 | (-0.002, 0.002) | 0.000 | (-0.001, 0.001) |
| | | $\beta_3$ | 1.661 | 0.000 | (-0.001, 0.001) | 0.000 | (-0.001, 0.001) | 0.000 | (-0.001, 0.001) |
| | | $\lambda_1$ | 1.173 | 0.000 | (-0.005, 0.006) | 0.000 | (-0.004, 0.004) | 0.000 | (-0.002, 0.003) |
| | | $\lambda_3$ | -0.111 | 0.002 | (-0.063, 0.085) | 0.004 | (-0.046, 0.055) | -0.001 | (-0.034, 0.031) |
| | Three | $\beta_1$ | 0.756 | -0.003 | (-0.005, -0.001) | 0.000 | (-0.002, 0.001) | 0.000 | (-0.001, 0.001) |
| | | $\beta_3$ | 1.661 | -0.001 | (-0.003, -0.000) | 0.000 | (-0.001, 0.001) | 0.000 | (-0.000, 0.001) |
| | | $\lambda_1$ | 1.173 | 0.002 | (-0.003, 0.007) | 0.000 | (-0.004, 0.004) | 0.000 | (-0.002, 0.003) |
| | | $\lambda_3$ | -0.111 | 0.164 | (0.084, 0.236) | 0.008 | (-0.044, 0.048) | 0.004 | (-0.022, 0.036) |
| | Four | $\beta_1$ | 0.756 | -0.030 | (-0.032, -0.027) | -0.001 | (-0.003, 0.000) | 0.000 | (-0.001, 0.001) |
| | | $\beta_3$ | 1.661 | -0.089 | (-0.092, -0.087) | 0.000 | (-0.001, 0.001) | 0.000 | (-0.001, 0.000) |
| | | $\lambda_1$ | 1.173 | 0.014 | (0.008, 0.020) | 0.002 | (-0.002, 0.006) | 0.000 | (-0.002, 0.002) |
| | | $\lambda_3$ | -0.111 | 4.934 | (4.824, 5.048) | 0.044 | (0.007, 0.093) | 0.003 | (-0.021, 0.030) |

Web Table 4: Read count simulation. True values and average relative bias of parameter estimates (and $2.5^{th}$, $97.5^{th}$ percentiles) across a hundred simulated data sets are shown for H3K36me3 with $10^5$ genomic windows. Scenarios with observed (obs.) SNR and 70% of observed SNR are shown.

| | | | | One Replicate | | Two Replicates | | Four Replicates | |
|---|---|---|---|---|---|---|---|---|---|
| SNR | Conditions | Parameter | True | R. Bias | $(P_{2.5},P_{97.5})$ | R. Bias | $(P_{2.5},P_{97.5})$ | R. Bias | $(P_{2.5},P_{97.5})$ |
| 70% of Observed SNR | Two | $\beta_1$ | 0.756 | 0.000 | (-0.006, 0.007) | 0.000 | (-0.005, 0.004) | 0.000 | (-0.003, 0.003) |
| | | $\beta_3$ | 1.304 | 0.000 | (-0.006, 0.005) | 0.000 | (-0.003, 0.004) | 0.000 | (-0.003, 0.003) |
| | | $\lambda_1$ | 1.173 | 0.000 | (-0.016, 0.018) | 0.002 | (-0.009, 0.015) | 0.000 | (-0.008, 0.008) |
| | | $\lambda_3$ | -0.467 | -0.001 | (-0.060, 0.064) | 0.004 | (-0.031, 0.040) | -0.001 | (-0.026, 0.023) |
| | Three | $\beta_1$ | 0.756 | 0.003 | (-0.004, 0.010) | -0.002 | (-0.007, 0.001) | 0.000 | (-0.003, 0.002) |
| | | $\beta_3$ | 1.304 | -0.119 | (-0.195, -0.086) | -0.001 | (-0.005, 0.002) | 0.000 | (-0.002, 0.002) |
| | | $\lambda_1$ | 1.173 | 0.025 | (-0.003, 0.047) | 0.003 | (-0.010, 0.015) | 0.000 | (-0.007, 0.007) |
| | | $\lambda_3$ | -0.467 | 0.694 | (0.586, 0.888) | 0.017 | (-0.020, 0.058) | 0.000 | (-0.024, 0.025) |
| | Four | $\beta_1$ | 0.756 | -0.010 | (-0.017, -0.003) | -0.019 | (-0.024, -0.015) | -0.001 | (-0.003, 0.002) |
| | | $\beta_3$ | 1.304 | -0.107 | (-0.119, -0.096) | -0.074 | (-0.080, -0.067) | 0.000 | (-0.002, 0.002) |
| | | $\lambda_1$ | 1.173 | 0.055 | (0.034, 0.080) | 0.044 | (0.028, 0.058) | 0.001 | (-0.006, 0.010) |
| | | $\lambda_3$ | -0.467 | 0.746 | (0.667, 0.835) | 0.571 | (0.517, 0.627) | 0.005 | (-0.017, 0.030) |
| Observed SNR | Two | $\beta_1$ | 0.756 | 0.000 | (-0.007, 0.006) | 0.000 | (-0.004, 0.004) | 0.000 | (-0.003, 0.003) |
| | | $\beta_3$ | 1.661 | 0.000 | (-0.004, 0.004) | 0.000 | (-0.003, 0.003) | 0.000 | (-0.002, 0.002) |
| | | $\lambda_1$ | 1.173 | 0.001 | (-0.014, 0.018) | 0.001 | (-0.011, 0.012) | 0.001 | (-0.009, 0.010) |
| | | $\lambda_3$ | -0.111 | 0.018 | (-0.195, 0.219) | 0.010 | (-0.128, 0.158) | 0.011 | (-0.091, 0.126) |
| | Three | $\beta_1$ | 0.756 | -0.004 | (-0.010, 0.003) | 0.000 | (-0.004, 0.005) | 0.000 | (-0.002, 0.003) |
| | | $\beta_3$ | 1.661 | -0.001 | (-0.005, 0.003) | 0.000 | (-0.003, 0.002) | 0.000 | (-0.001, 0.002) |
| | | $\lambda_1$ | 1.173 | 0.003 | (-0.013, 0.020) | -0.001 | (-0.013, 0.011) | -0.001 | (-0.007, 0.006) |
| | | $\lambda_3$ | -0.111 | 0.187 | (-0.024, 0.451) | -0.005 | (-0.164, 0.128) | -0.007 | (-0.097, 0.097) |
| | Four | $\beta_1$ | 0.756 | -0.030 | (-0.038, -0.022) | -0.002 | (-0.005, 0.003) | 0.000 | (-0.002, 0.002) |
| | | $\beta_3$ | 1.661 | -0.088 | (-0.094, -0.081) | 0.000 | (-0.002, 0.002) | 0.000 | (-0.001, 0.001) |
| | | $\lambda_1$ | 1.173 | 0.013 | (-0.007, 0.033) | 0.002 | (-0.010, 0.014) | 0.000 | (-0.007, 0.007) |
| | | $\lambda_3$ | -0.111 | 4.870 | (4.528, 5.273) | 0.052 | (-0.106, 0.209) | -0.001 | (-0.077, 0.092) |

ties pertaining to the differential state by controlling the total FDR as defined in Section 3.2 (main text). For different nominal FDR threshold levels, the model sensitivity was estimated as the proportion of windows correctly assigned as differential out of the total number of simulated differential windows. Additionally, the observed FDR was calculated as the proportion of genomic windows incorrectly called as differential out of the total number of called differential windows. In Figures 13-15, panel A shows the average observed true positive rate (y-axis) and the observed FDR (x-axis) for different nominal FDR levels across a hundred simulated data. Results are shown for different levels of SNR, number of conditions, and number of replicates per condition. Overall, we observed that the number of replicates per condition played a major role on the sensitivity levels of the model, in which scenarios with two and four replicates had the best results regardless of the number of conditions and SNR levels. For scenarios with either high number of conditions or low SNR levels, more replicates were needed to achieve higher sensitivity.

Finally, we used the estimated mixture model posterior probabilities $Pr(W_{jl} = 1|Z_j = 2, \mathbf{y}_{..j}, \mathbf{x}; \hat{\boldsymbol{\Psi}})$, $j = 1, \ldots, M$, to classify the differential combinatorial patterns of enrichment of detected differential windows. To this end, we first calculated the maximum estimated mixture model posterior probability across all $L$ components to determine the most likely differential combinatorial pattern from genomic windows assigned to be part of the differential state. Then, we compared the window-based classification with the true window-based simulated states from the Markov Chain (states $2, \ldots, G-1$). In Figures 13-15, panel B shows the confusion matrices of classified (x-axis) and simulated (y-axis) differential windows for a scenario with three conditions. Differential combinatorial patterns of enrichment are represented by the sequences of letters 'E' (enrichment) and 'B' (background), such that each letter corresponds to the status of a given condition. The number of windows (averaged over a all simulated data sets) is shown as entries of the matrices and represented by the color scale. Darker colors on the diagonal entries indicate better agreement between simulated and classified patterns. By utilizing the posterior probabilities from the mixture model, we observed a good performance when classifying the differential combinatorial pattern of enrichment from differential windows. Results were best under scenarios with higher number of replicates or SNR.

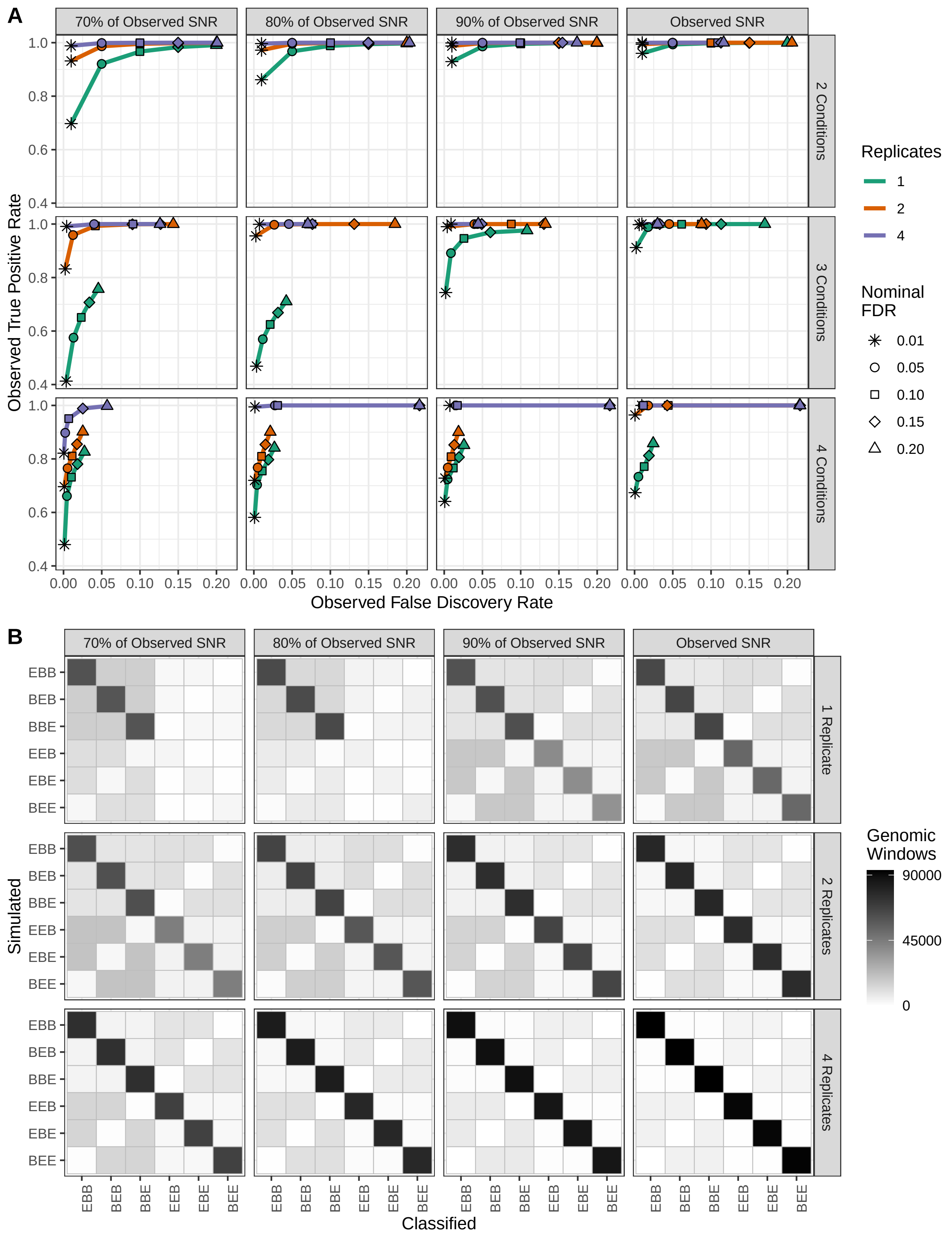


Web Figure 13: Read count simulation. (A): average observed TPR and FDR for different nominal FDR threshold levels for various scenarios in simulated data of H3K27me3 with $10^6$ genomic windows. (B): confusion matrices are shown for the 3 conditions scenario. On x- and y-axes the labels indicate the differential classified and simulated patterns, respectively. For instance, EBE denotes enrichment in conditions 1 and 3 only. Darker colors on the diagonal of the matrix indicate better agreement.

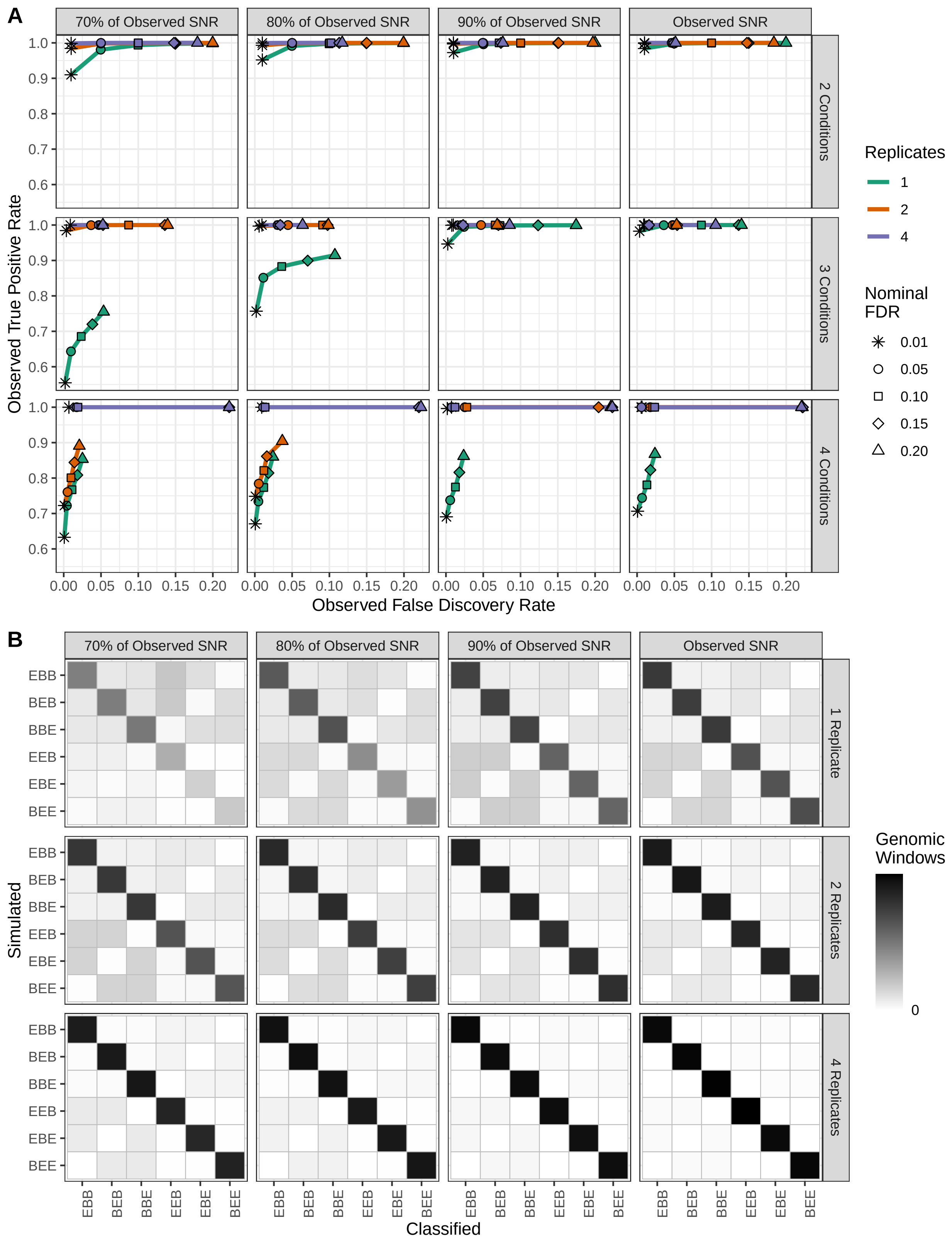


Web Figure 14: Read count simulation. (A): average observed TPR and FDR for different nominal FDR threshold levels for various scenarios in simulated data of H3K36me3 with $10^5$ genomic windows. (B): confusion matrices are shown for the 3 conditions scenario. On x- and y-axes the labels indicate the differential classified and simulated patterns, respectively. For instance, EBE denotes enrichment in conditions 1 and 3 only. Darker colors on the diagonal of the matrix indicate better agreement.

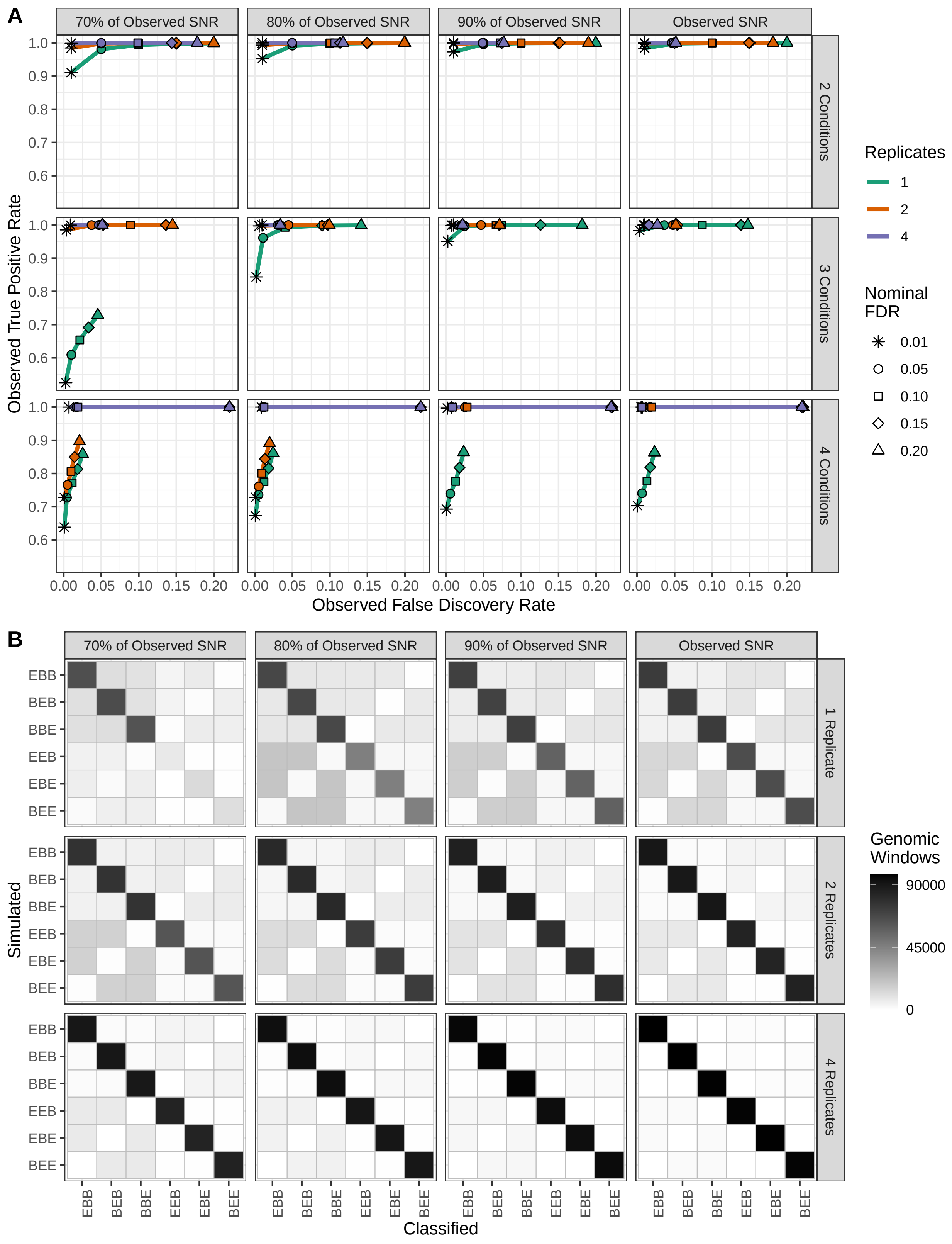


Web Figure 15: Read count simulation. (A): average observed TPR and FDR for different nominal FDR threshold levels for various scenarios in simulated data of H3K36me3 with $10^6$ genomic windows. (B): confusion matrices are shown for the 3 conditions scenario. On x- and y-axes the labels indicate the differential classified and simulated patterns, respectively. For instance, EBE denotes enrichment in conditions 1 and 3 only. Darker colors on the diagonal of the matrix indicate better agreement.

### C2 Sequencing Read Simulation

In this section, we present additional results from the sequencing read simulation using different FDR control threshold for differential peak call.

We performed a second simulation study aiming to compare the proposed model with the current DPCs ChIPComp, csaw, DiffBind, diffReps, RSEG, and THOR, since none of them was designed to accept matrices of window-based read counts as input. To this purpose, we revisited the simulation study presented by Lun and Smyth, 2015 where data were generated in a more general scheme without a particular read count model assumption. Here, sequencing reads from ChIP-seq experiments were generated for two conditions and two replicates per condition. For the differential peaks callers ChIPComp and DiffBind that require sets of candidate regions, we followed the analyses presented by Lun and Smyth, 2015 and called peaks in advance using HOMER. Those were then used as input in the respective software for differential call. For ChIPComp and DiffBind, we attempted to call peaks in advance using MACS. However, MACS exhibited issues to work on simulated data as it was not able to detect peaks. Here, we adapted their simulation pipeline aiming to generate broad and diffuse differential regions of enrichment. Specifically, a total of 4000 binding sites spaced 60-65 kbp apart were simulated such that each site was formed by three neighboring peaks 10kb wide. For each peak, the number of sequencing reads were simulated from a NB distribution with mean 300 and dispersions sampled from an inverse chi-squared distribution with 20 degrees of freedom. Reads were positioned on peak sites to form smooth binding profiles. To introduce differential enrichment, one or two peaks from 1000 binding sites were randomly chosen and reads pertaining to them were removed from the sequencing library in one of the two conditions. This process was repeated for the second condition, which led to a total of 2000 differential peaks. Background reads were added to the library to maintain the realistic aspects of the experiment. A hundred simulated data sets were generated and peaks were called by all the methods under multiple nominal FDR thresholds. For our method and RSEG, window-based posterior probabilities were used to control the total FDR. For illustration purposes, we refer to the method presented in this article as mixNBHMM.

Specifically, panel D from Figures 16, 17, 18, and 19 show example of peak calls using FDR control cutoffs 0.01, 0.10, 0.15, and 0.20, respectively. Overall, results were consistent across different choices of FDR control. This fact is also shown by the observed sensitivity and false discovery rate from all methods (panels A). We observed a clear distinction of methods regarding the observed sensitivity and specificity such that RSEG, diffReps, and THOR exhibited the highest observed FDR and ChIPComp and DiffBind exhibited the lowest sensitivity.

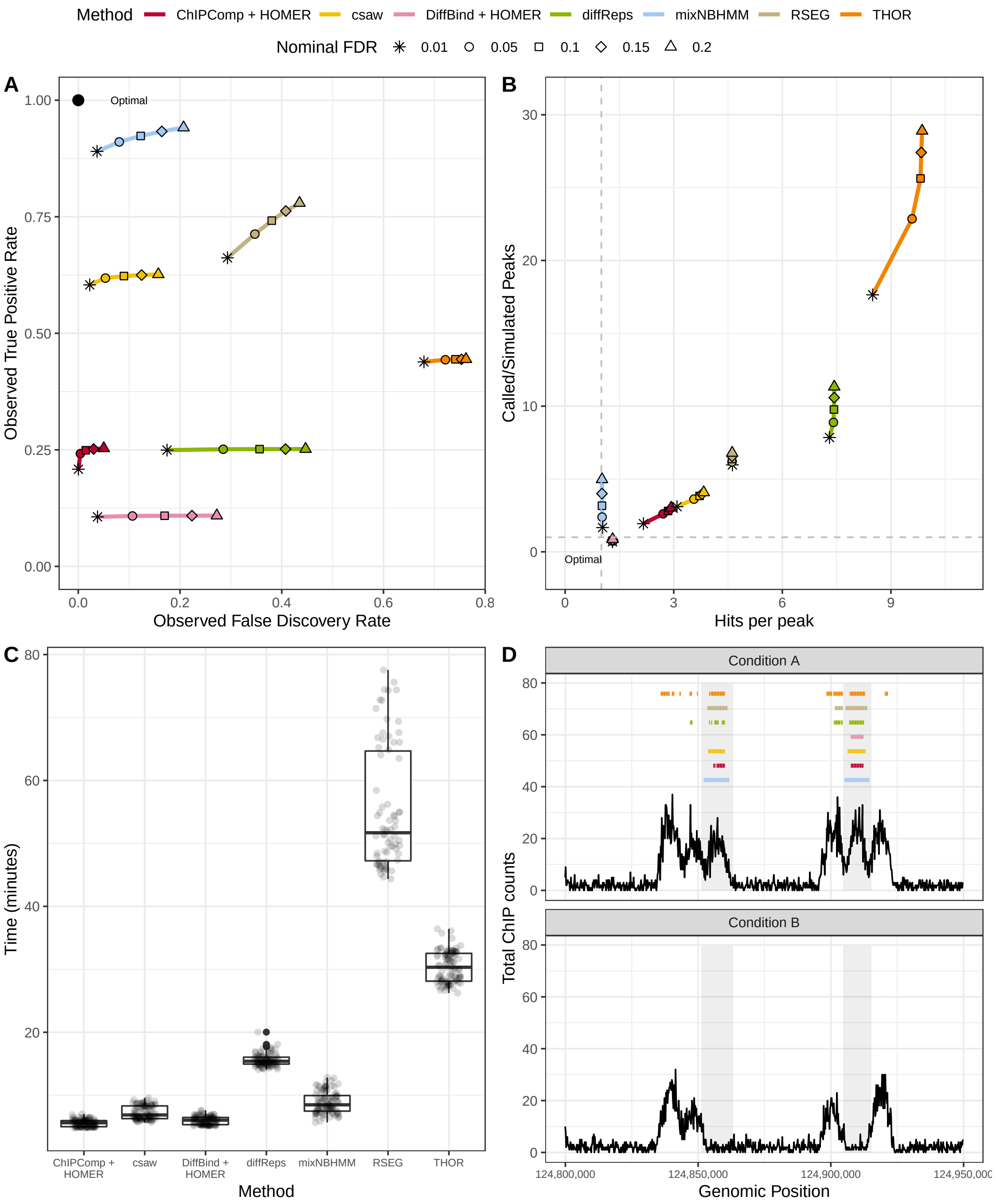


Web Figure 16: Sequencing read simulation. (A): average observed sensitivity and FDR for various methods under different nominal FDR levels. (B): scatter plot of average ratio of called and simulated peaks (y-axis) and average number of called peaks intersecting true differential regions (x-axis) for different nominal FDR levels. (C): box plot of computing time (in minutes) for various algorithms. (D): an example of differential peak calls under a nominal FDR threshold 0.01. Shaded areas indicate true differential peaks.

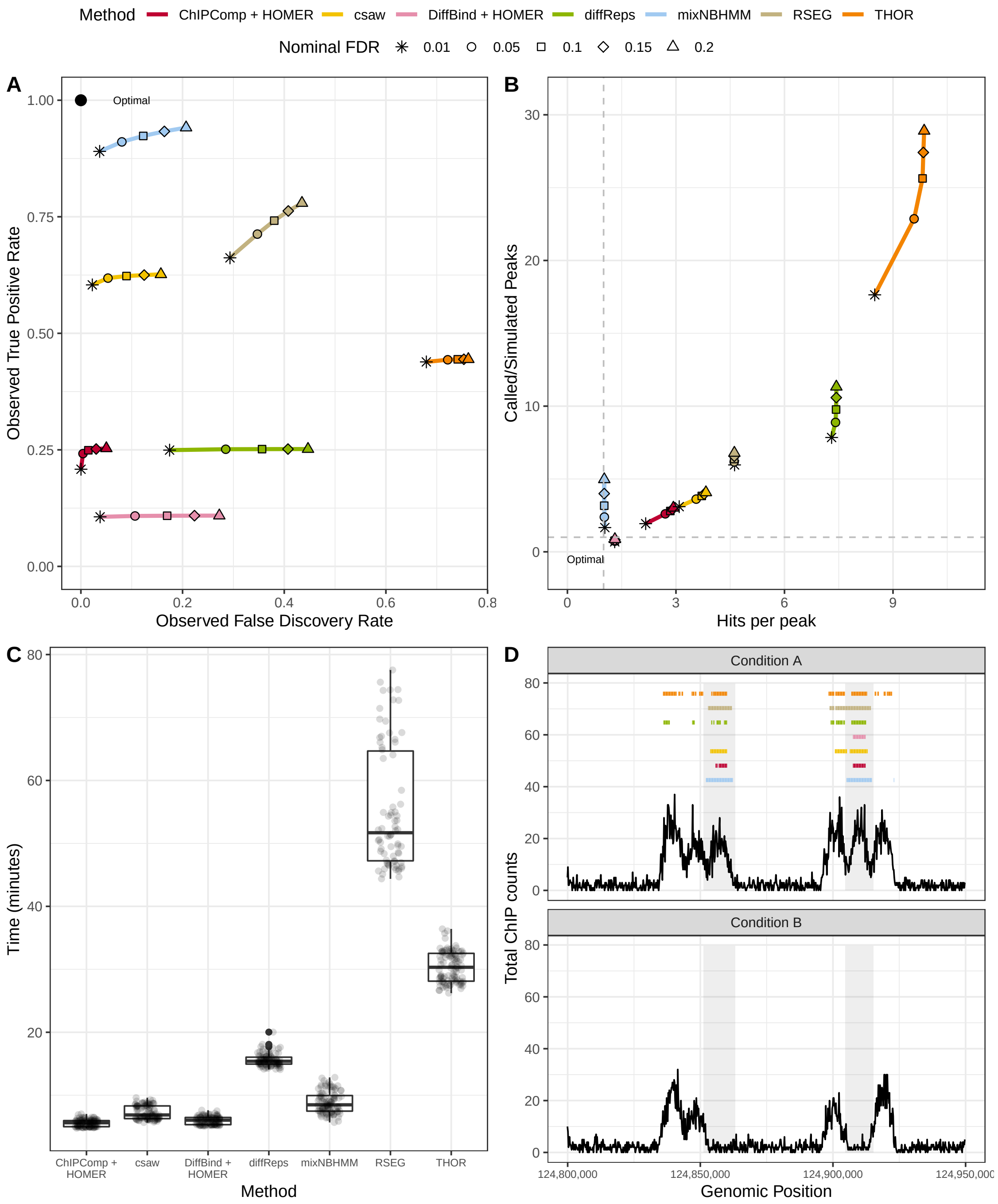


Web Figure 17: Sequencing read simulation. (A): average observed sensitivity and FDR for various methods under different nominal FDR levels. (B): scatter plot of average ratio of called and simulated peaks (y-axis) and average number of called peaks intersecting true differential regions (x-axis) for different nominal FDR levels. (C): box plot of computing time (in minutes) for various algorithms. (D): an example of differential peak calls under a nominal FDR threshold 0.10. Shaded areas indicate true differential peaks.

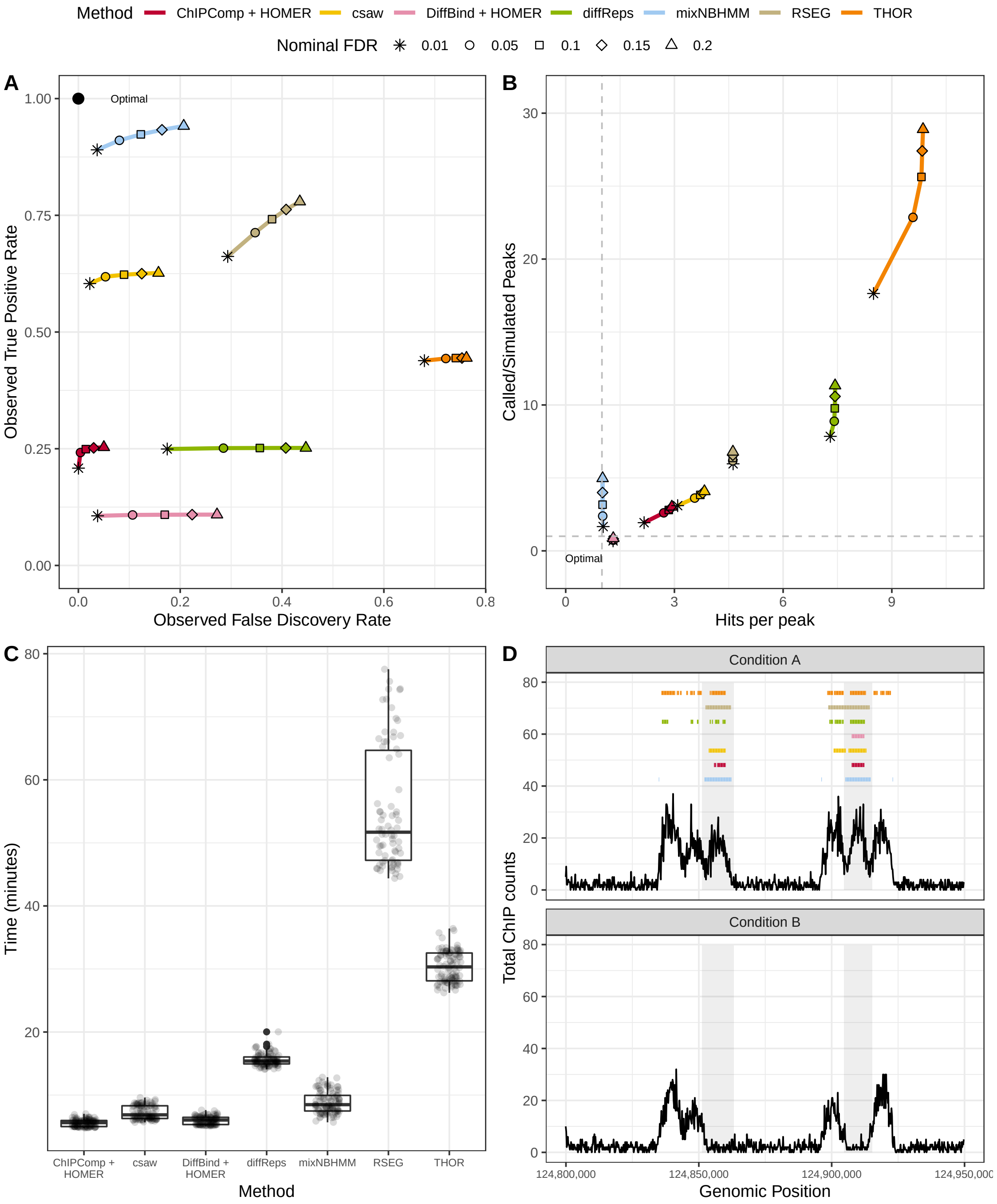


Web Figure 18: Sequencing read simulation. (A): average observed sensitivity and FDR for various methods under different nominal FDR levels. (B): scatter plot of average ratio of called and simulated peaks (y-axis) and average number of called peaks intersecting true differential regions (x-axis) for different nominal FDR levels. (C): box plot of computing time (in minutes) for various algorithms. (D): an example of differential peak calls under a nominal FDR threshold 0.15. Shaded areas indicate true differential peaks.

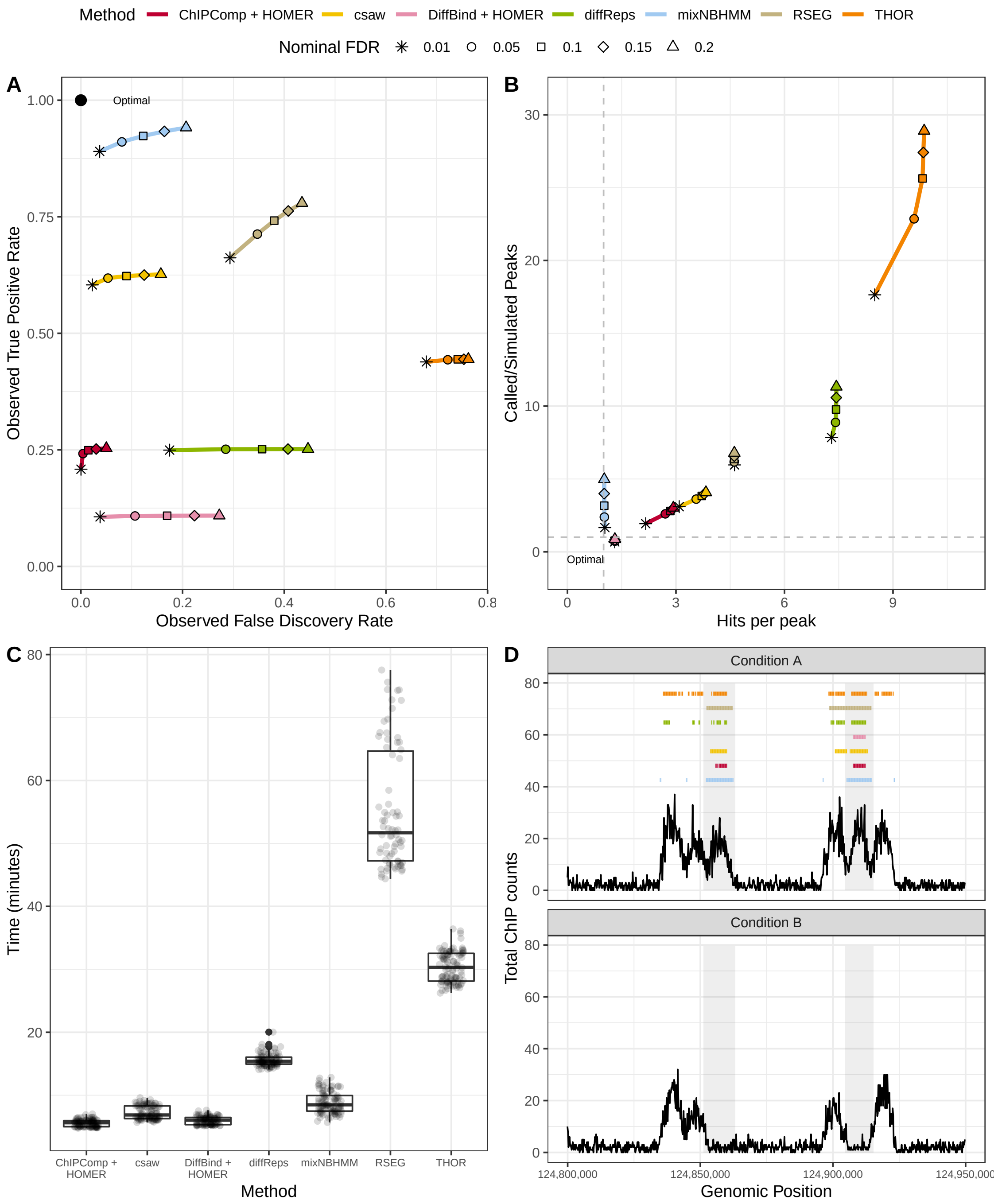


Web Figure 19: Sequencing read simulation. (A): average observed sensitivity and FDR for various methods under different nominal FDR levels. (B): scatter plot of average ratio of called and simulated peaks (y-axis) and average number of called peaks intersecting true differential regions (x-axis) for different nominal FDR levels. (C): box plot of computing time (in minutes) for various algorithms. (D): an example of differential peak calls under a nominal FDR threshold 0.20. Shaded areas indicate true differential peaks.

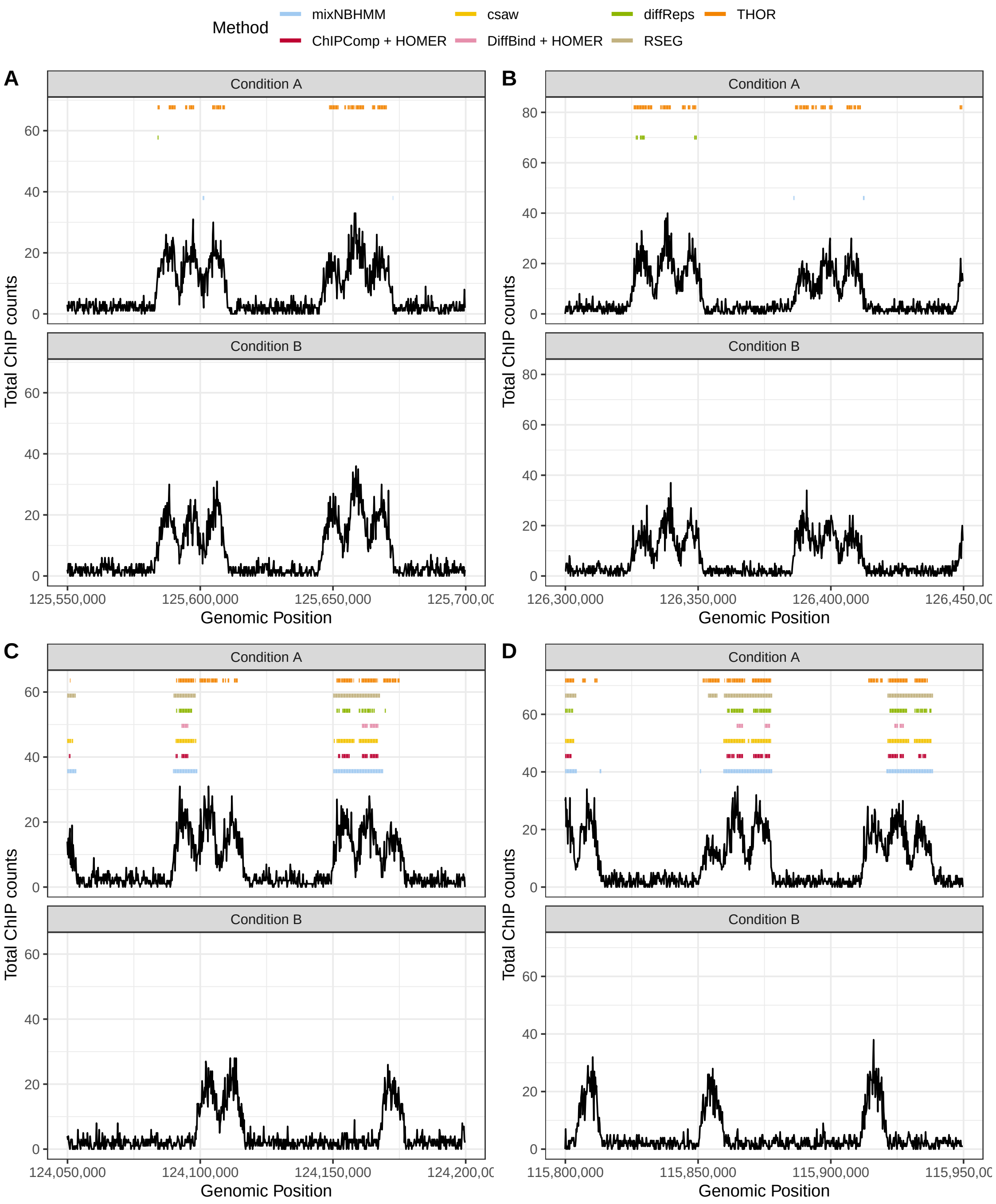


Web Figure 20: Sequencing read simulation. Examples of differential called peaks from the various methods under an FDR control of 0.05.

### Supporting Information D

In this appendix, we present an overview of the application on data from the ENCODE Consortium.

#### D1 RNA-seq data and gene transcription

Under the three cell lines scenario (presented in the main text), we used RNA-seq experimental data from the ENCODE Consortium on Helas3, Hepg2, and Huvec human cells to quantify the expression levels of genes as follows. First, we used Salmon (Patro et al., 2017) to quantify transcript expression from cell-specific RNA-seq experiments. We then calculated, using the R package *tximport* (Soneson et al., 2015), estimated counts using abundance estimates (transcripts per million, TPM) scaled up to the average transcript length over samples and library size. This step ensures that counts computed from Salmon are not correlated with the average transcript length. Secondly, we calculated the number of ChIP-seq reads from H3K4me3 and H3K27ac overlapping gene promoters. Promoter regions extend around the transcription start site 2000 base pairs upstream and 200 base pair downstream. Finally, read counts from RNA-seq and ChIP-seq were normalized for sequencing depth using DESeq2.

For the two cell line scenario, differentially transcribed genes were defined through log2 fold changes of H3K36me3 ChIP-seq read counts. We observed several cases in which differentially expressed genes (defined by RNA-seq data) did not exhibit differential enrichment for H3K36me3. However, due to the activating roles of H3K36me3 on gene transcription, we follow the ideas presented by Steinhauser et al., 2016 and Ji et al., 2013 and defined differentially transcribed genes based on log2 fold changes of H3K36me3 ChIP-seq read counts.

Sensitivity and specificity metrics were calculated on the window-level as follows. For non-overlapping genomic windows $b_1, \ldots, b_M$, let $g_j = I(b_j \in \text{ differentially transcribed gene})$ and $d_j = I(b_j \in \text{ differential peak})$ denote the indicators of genomic windows being associated with either differential gene bodies or differential peaks, respectively, for $j = 1, \ldots, M$, for a given method and nominal FDR level. Then, the observed sensitivity (TPR), specificity (1-FPR), and FDR were calculated as follows:

$$\text{Sensitivity} = \frac{\sum_{j=1}^{M} g_j d_j}{\sum_{j=1}^{M} g_j}$$

$$\text{Specificity} = \frac{\sum_{j=1}^{M} (1 - g_j)(1 - d_j)}{\sum_{j=1}^{M} (1 - g_j)}$$

$$\text{FDR} = \frac{\sum_{j=1}^{M} (1 - g_j) d_j}{\sum_{j=1}^{M} d_j}$$

We present additional results using log2 FC 1, 1.5, 2.5, and 3 in Figure 21. Overall the presented model outperformed all current methods regardless of the choice of FC cutoff, as we maintained the highest sensitivity for a limited observed false discovery rate. RSEG showed significant higher observed false discovery rate in all the scenarios.

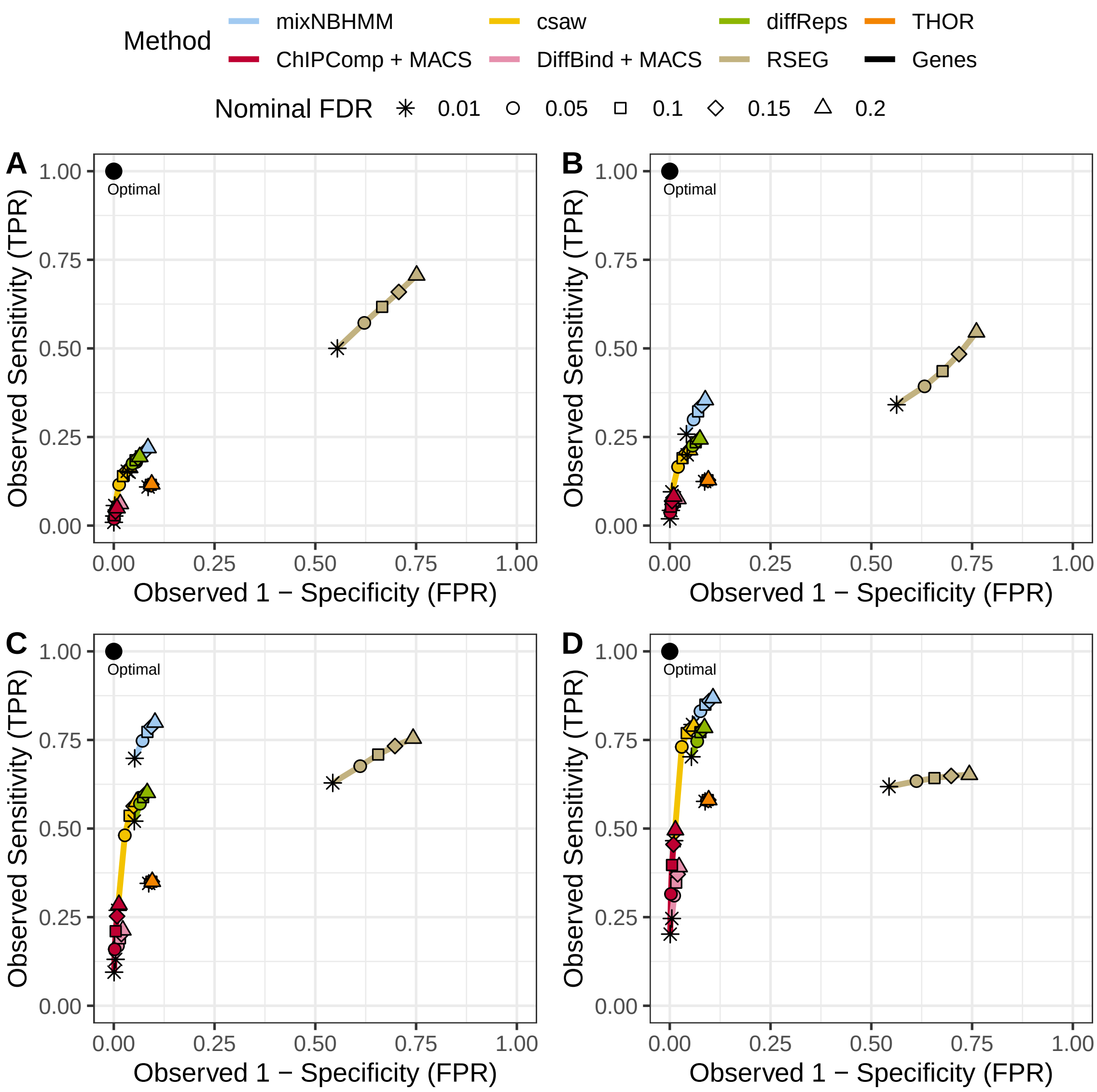


Web Figure 21: ROC curves for H3K36me3 differential peaks covering differentially transcribed gene bodies defined if having absolute $LFC > 1$ (A), $LFC > 1.5$ (B), $LFC > 2.5$ (C), or $LFC > 3$ (D).

#### D2 Study on Window Sizes

We compared methods regarding the metrics used in this paper to analyze broad marks using window sizes of 250bp, 500bp , 750bp, and 1000bp. Overall, results were consistent across different window sizes. The HMM-based methods RSEG and THOR appeared to be highly sensitive to the window size and showed better results for 1000bp. However, the presented model appeared to be robust to all scenarios as it consistently called a small number of broad peaks overall (Figures 22, 23, and 24, panels C). The methods csaw, ChIPComp, diffReps, and DiffBind exhibited fragmented and narrow differential peaks regardless of the window size. The performance of all methods appeared to be consistent to the choice of window size under sharp data set. The presented model was robust in all the scenarios as it was able to detect large differences between read counts of the analyzed cell lines.

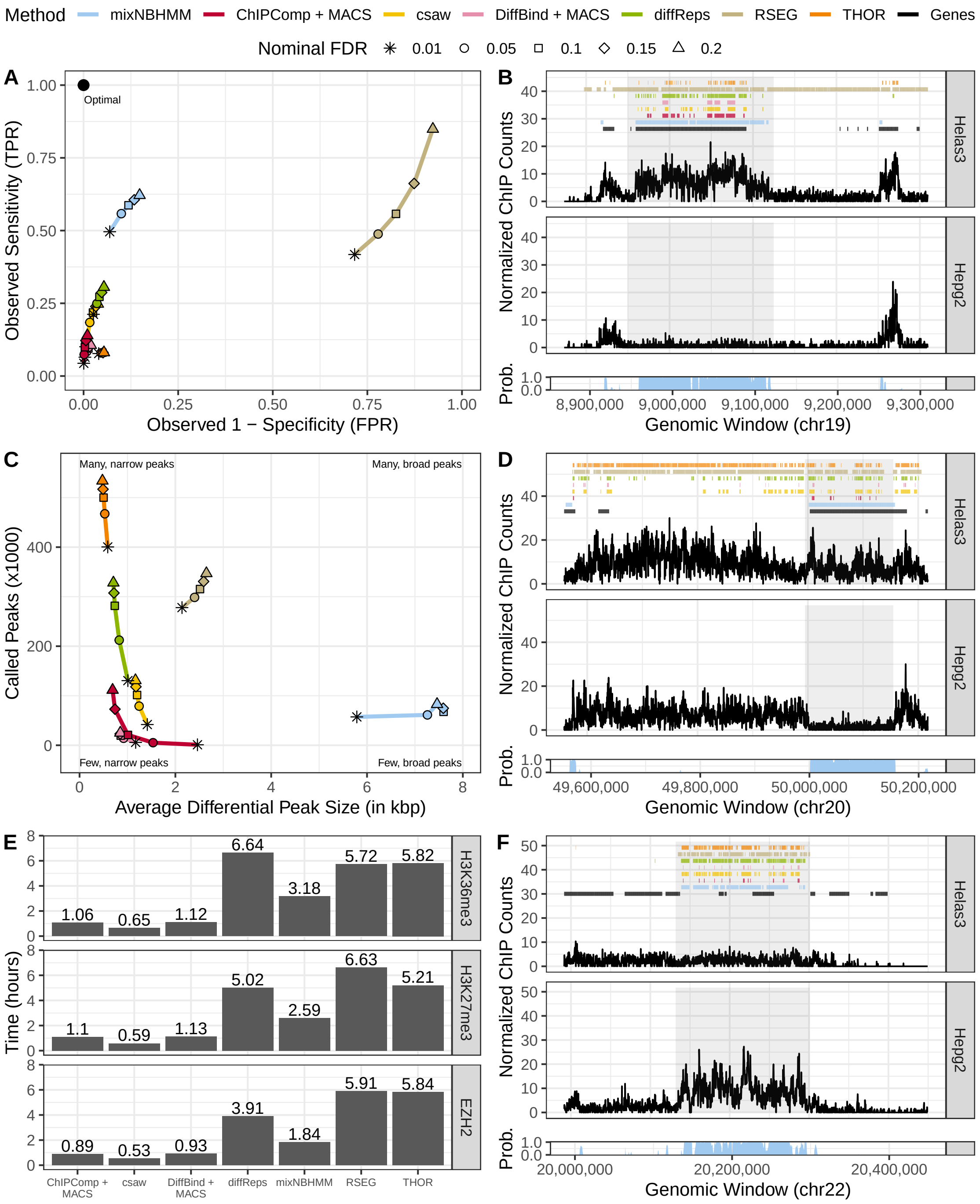


Web Figure 22: Results from broad marks (250bp). (A): ROC curves of differentially transcribed gene coverage based on H3K36me3 differential peak calls. (C): average number (y-axis) and size (x-axis) of H3K27me3 called peaks for various methods and different nominal FDR thresholds. (B), (D), and (F): example of peak calls from H3K36me3, H3K27me3, and EZH2, respectively. (E): computing time of genome-wide analysis of broad marks from various methods.

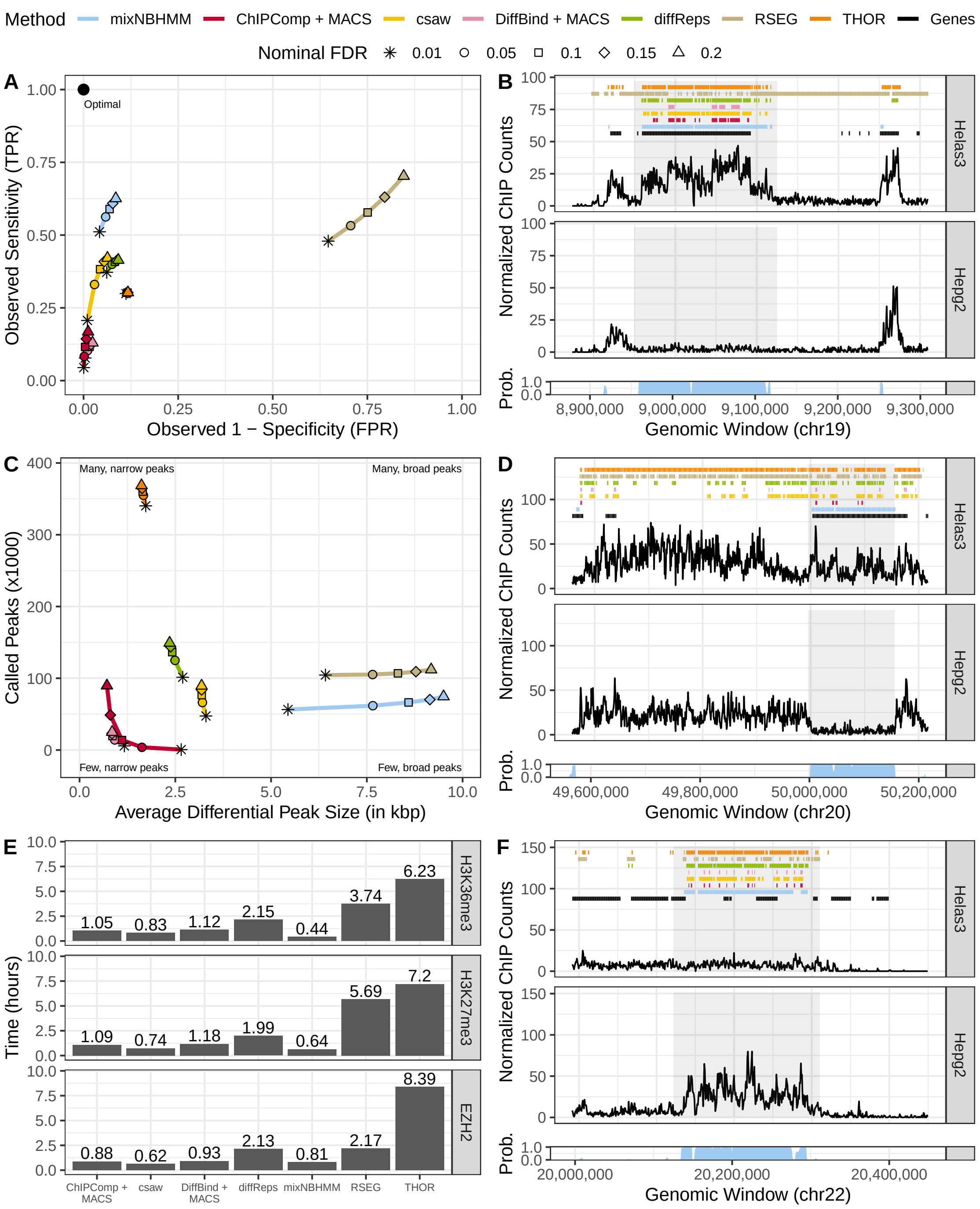


Web Figure 23: Results from broad marks (750bp). (A): ROC curves of differentially transcribed gene coverage based on H3K36me3 differential peak calls. (C): average number (y-axis) and size (x-axis) of H3K27me3 called peaks for various methods and different nominal FDR thresholds. (B), (D), and (F): example of peak calls from H3K36me3, H3K27me3, and EZH2, respectively. (E): computing time of genome-wide analysis of broad marks from various methods.

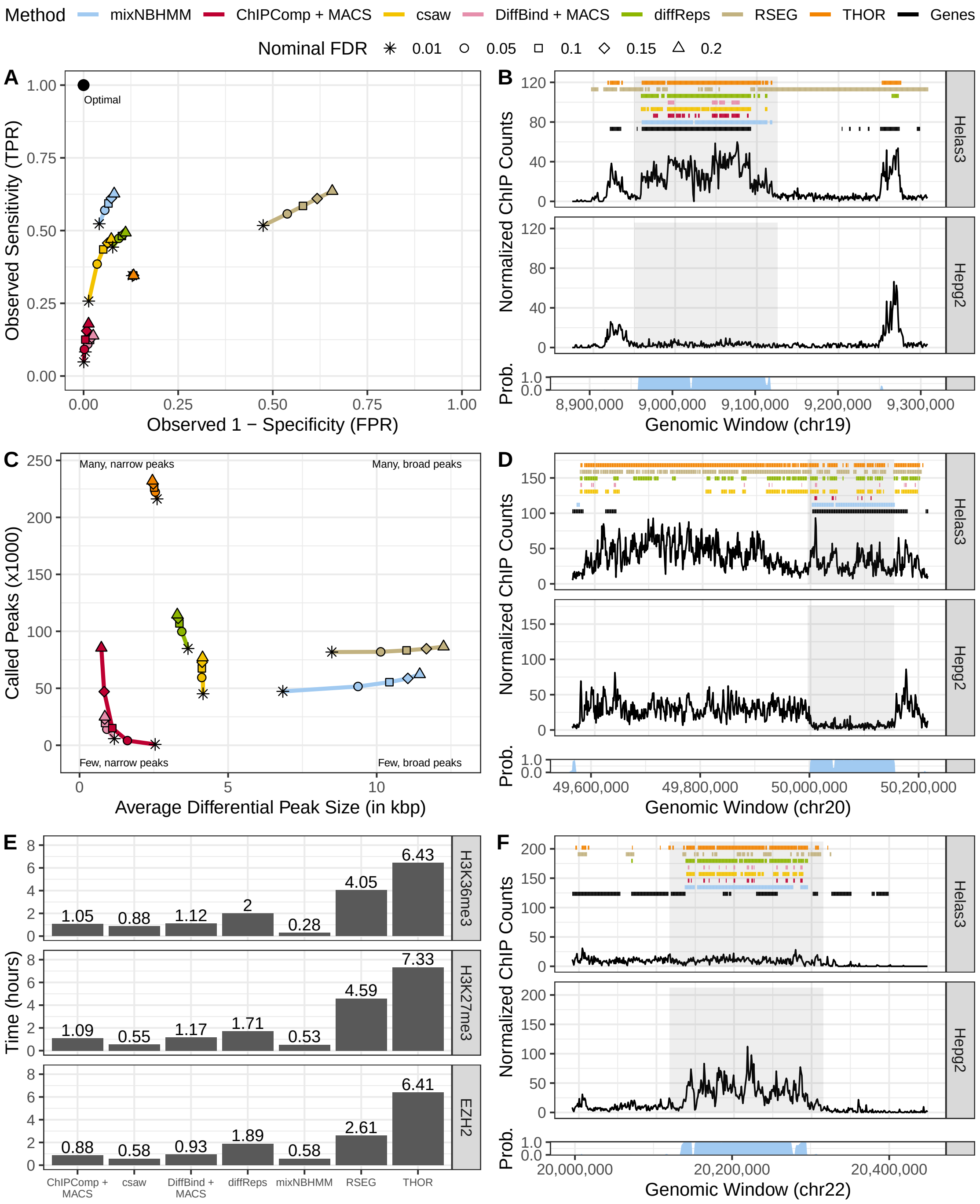


Web Figure 24: Results from broad marks (1000bp). (A): ROC curves of differentially transcribed gene coverage based on H3K36me3 differential peak calls. (C): average number (y-axis) and size (x-axis) of H3K27me3 called peaks for various methods and different nominal FDR thresholds. (B), (D), and (F): example of peak calls from H3K36me3, H3K27me3, and EZH2, respectively. (E): computing time of genome-wide analysis of broad marks from various methods.

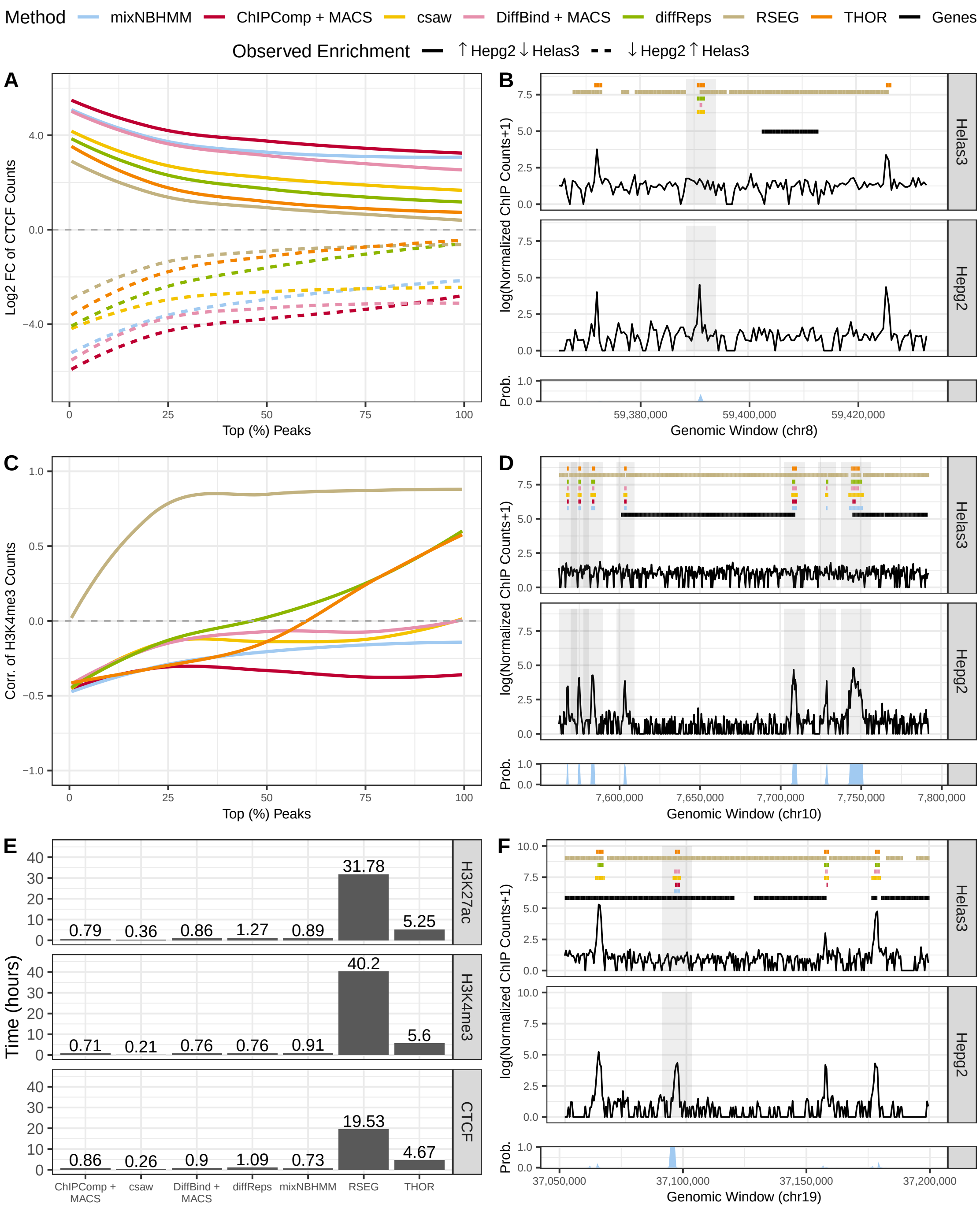


Web Figure 25: Results from sharp marks (500bp). (A) and (C): median $\log_2 FC$ and correlation between cell lines of ChIP-seq counts from differential called peaks (FDR 0.05, sorted by the absolute LFC) for CTCF and H3K4me3, respectively. (B), (D), and (F): example of broad peak calls from CTCF, H3K4me3, and H3K27ac, respectively. (E): computing time of genome-wide analysis of sharp marks from various methods.

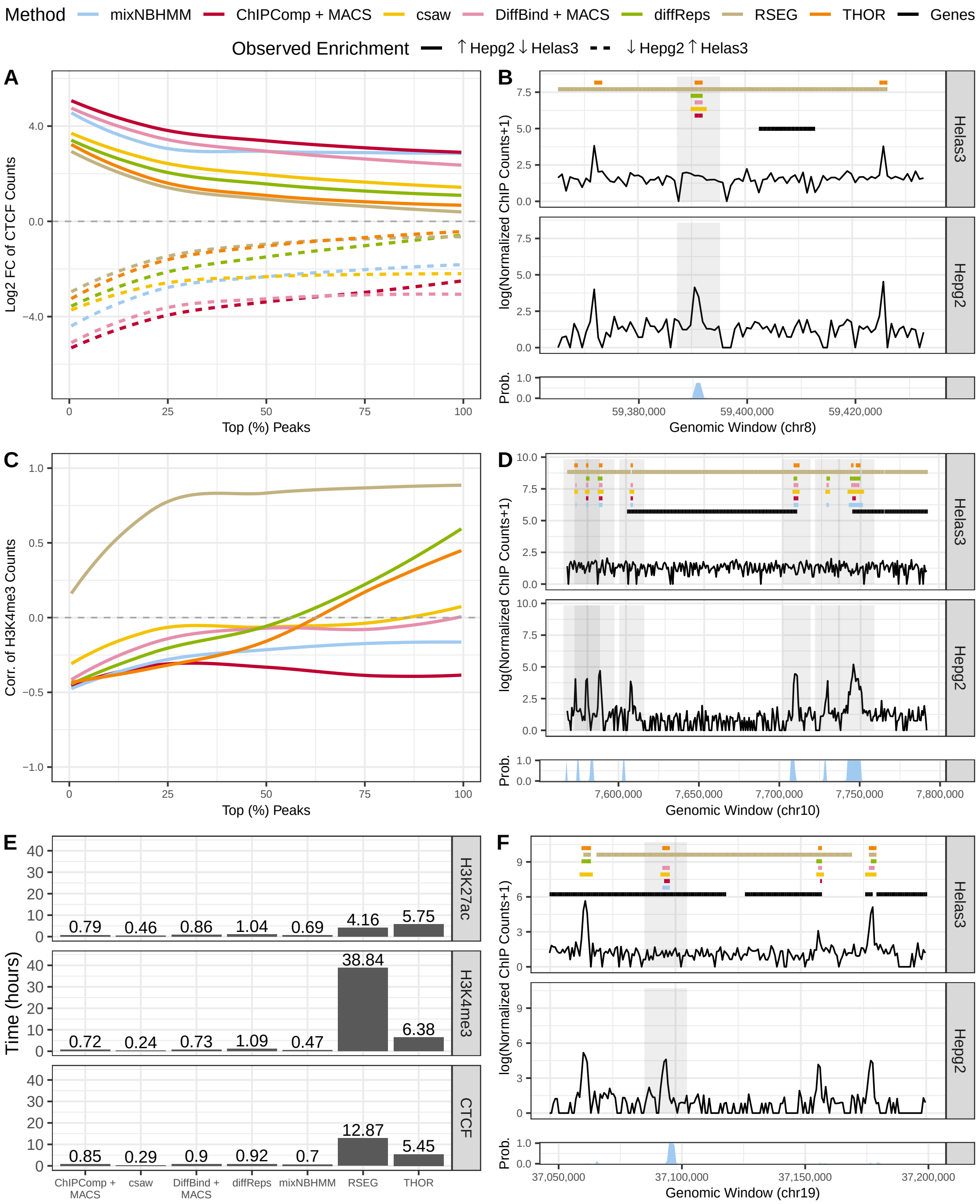


Web Figure 26: Results from sharp marks (750bp). (A) and (C): median $\log_2 FC$ and correlation between cell lines of ChIP-seq counts from differential called peaks (FDR 0.05, sorted by the absolute LFC) for CTCF and H3K4me3, respectively. (B), (D), and (F): example of broad peak calls from CTCF, H3K4me3, and H3K27ac, respectively. (E): computing time of genome-wide analysis of sharp marks from various methods.

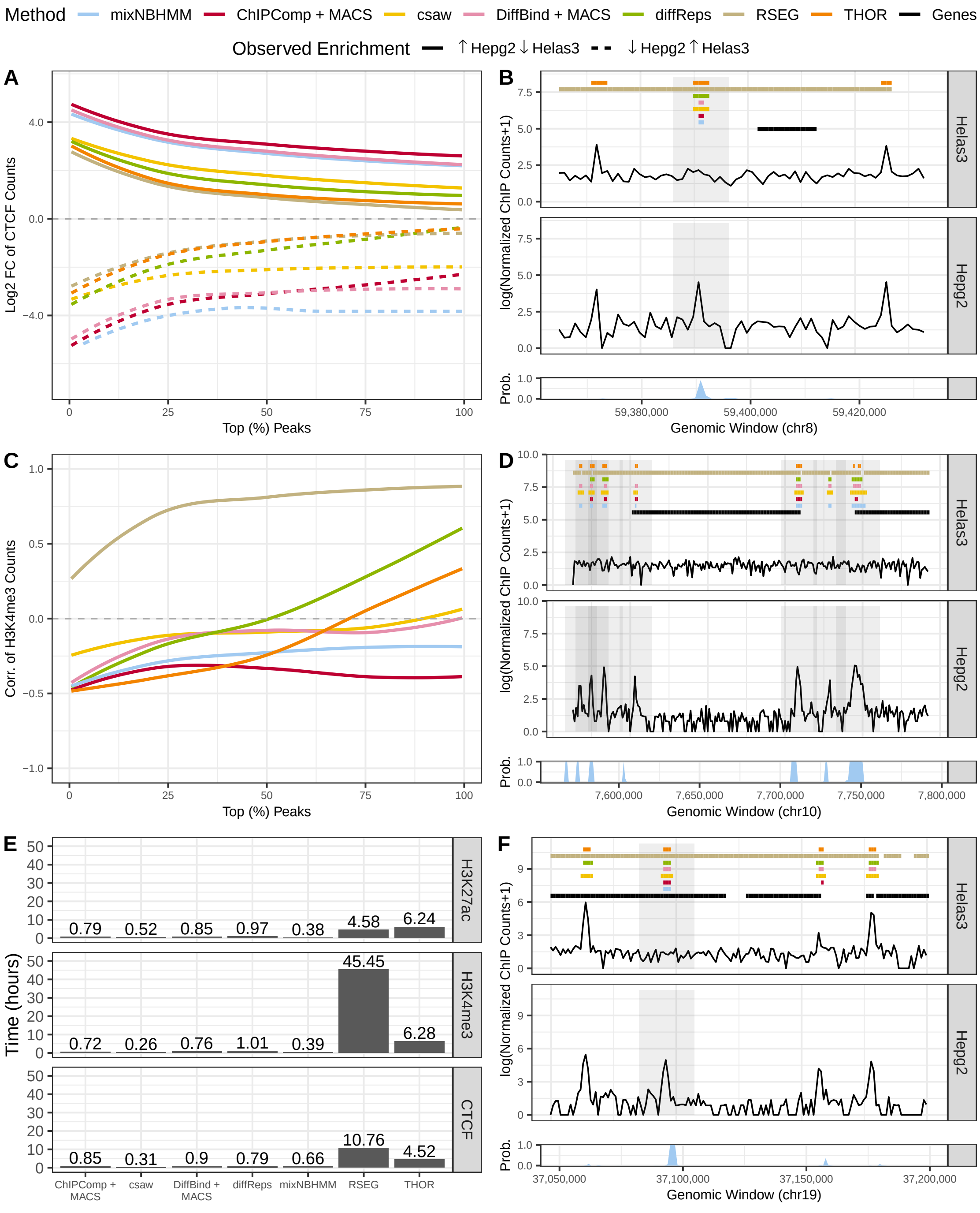


Web Figure 27: Results from sharp marks (1000bp). (A) and (C): median $\log_2 FC$ and correlation between cell lines of ChIP-seq counts from differential called peaks (FDR 0.05, sorted by the absolute LFC) for CTCF and H3K4me3, respectively. (B), (D), and (F): example of broad peak calls from CTCF, H3K4me3, and H3K27ac, respectively. (E): computing time of genome-wide analysis of sharp marks from various methods.

### D3 Classification of Combinatorial Patterns of Protein-DNA Binding Activity and Their Associations With Gene Expression

Lastly, we jointly analyzed data from cell lines Helas3, Hepg2, and Huvec to assess the performance and biological relevance of the classification of differential combinatorial patterns of enrichment from H3K4me3 and H3K27ac. These epigenomic marks have been associated with gene transcription and are often deposited on promoter regions of actively transcribed gene bodies. First, we evaluated the classification performance of differential combinatorial patterns of enrichment across cell lines by making use of the estimated mixture model posterior probabilities. Given the differential HMM state, these posterior probabilities allow us assess whether read counts from genomic windows are likely to be generated from a particular combinatorial pattern of enrichment. Secondly, we compared the biological role of the classified patterns with gene expression data. To this end, RNA-seq and ChIP-seq read counts were mapped onto gene bodies and gene promoters, respectively. For the former, we used counts from abundance estimates scaled using the average transcript length over samples (Soneson et al., 2015). Using DESeq2 (Love et al., 2014), normalization between samples was performed to avoid spurious differences due to sequencing depth in both RNA-seq and ChIP-seq data. Then, pairwise LFC of read counts between cell lines were calculated. Genes with zero counts or with low total read count across cell lines were excluded from the analysis (25% of the protein-coding genes). Finally, we used the maximum window-based posterior probability of the presented mixture model to classify the combinatorial patterns of enrichment from the resulting genes whose promoters overlapped differential peaks.

Figure 29 shows the main results from our analyses under a nominal FDR control of 0.05. At the top, we present ternary density plots of differential *peaks* and their associated ChIP-seq read counts. The colors indicate the classified differential pattern of enrichment of peaks based on window-based posterior probabilities of the mixture model. We observed that the classification pattern agreed with the observed direction of enrichment from ChIP-seq data (vertices and opposite sides of the triangle vertices). At the bottom, we show the scatter plots of LFCs between cell lines of *gene* bodies (RNA-seq counts) and *gene* promoters (ChIP-seq counts), along with their Pearson correlation, for some of the classified combinatorial patterns of enrichment. As expected from the activating roles of H3K27ac and H3K4me3, we observed a high agreement between ChIP-seq and RNA-seq LFCs. Here, the colors indicate the classified differential pattern of activation of genes overlapping differential called peaks. These results show that the classified pattern of epigenomic change across the three cell lines agreed with the observed direction of gene expression and highlight the novelties of the presented model. We were able to accurately classify the differential pattern of enrichment of both called peaks (ternary plots) and genes overlapping called peaks (scatter plots) under more than two conditions. The embedded mixture model and its posterior probabilities generalizes the problem of differential peak calling for multiple conditions and provide a tool that is not yet available in any other method presented in the literature.

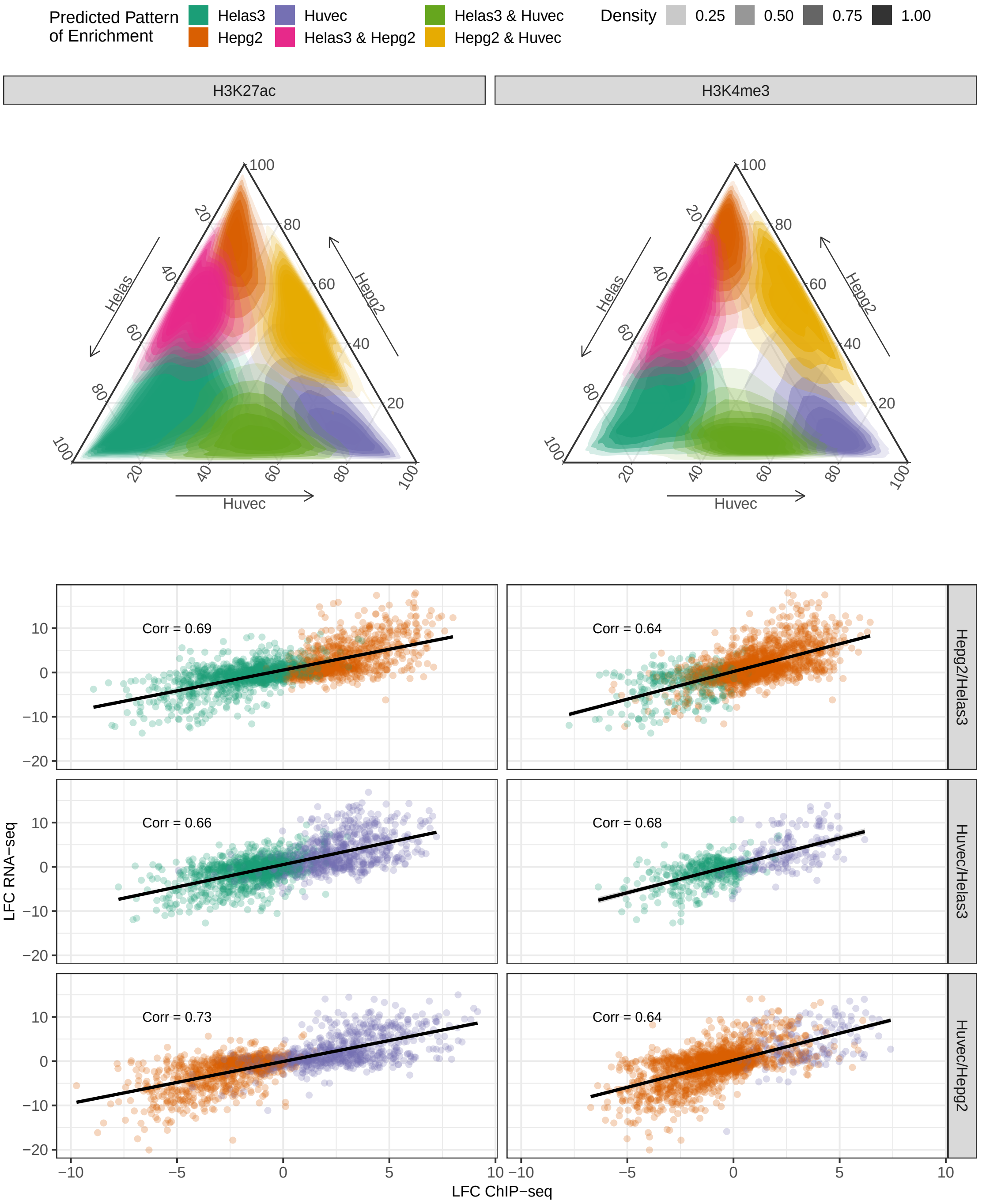


Web Figure 28: Ternary plots (top) of ChIP-seq counts mapped on differential peaks and scatter plots (bottom) of $\log_2 FC$s (LFC) between cell lines of ChIP-seq (x-axis) and RNA-seq counts (y-axis) mapped on genes intersecting differential peaks. Colors indicate the classified combinatorial pattern of peak enrichment (ternary plots) and gene activation (scatter plots). Correlations indicate high agreement between ChIP-seq and RNA-seq $\log_2 FC$s for these activating marks.

#### D4 Genomic Segmentation Analysis of H3K36me3 and CTCF

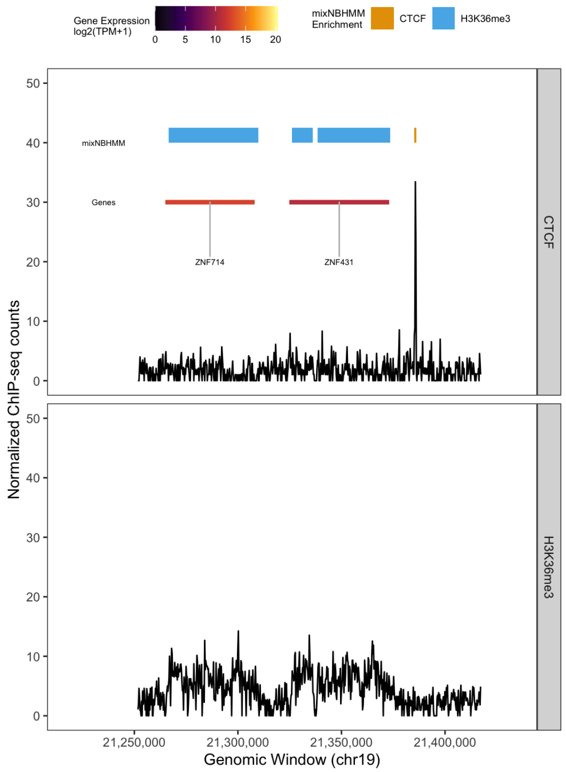


Web Figure 29: Genomic segmentation analysis of H3K36me3 and CTCF in Helas3 cell line. The chosen model parametrization and the normalization for non-linear biases via model offsets allow the segmentation of highly diverse epigenomic marks. The implemented hidden Markov model is able to properly account for the differences in length of enrichment regions between CTCF (short) and H3K36me3 (broad).
